# Plant pathogens convergently evolved to counteract redundant nodes of an NLR immune receptor network

**DOI:** 10.1101/2021.02.03.429184

**Authors:** Lida Derevnina, Mauricio P. Contreras, Hiroaki Adachi, Jessica Upson, Angel Vergara Cruces, Rongrong Xie, Jan Sklenar, Frank L.H. Menke, Sam T. Mugford, Dan MacLean, Wenbo Ma, Saskia Hogenhout, Aska Goverse, Abbas Maqbool, Chih-Hang Wu, Sophien Kamoun

## Abstract

In plants, NLR (nucleotide-binding domain and leucine-rich repeat-containing) proteins can form receptor networks to confer hypersensitive cell death and innate immunity. One class of NLRs, known as NRCs (NLR required for cell death), are central nodes in a complex network that protects against multiple pathogens and comprises up to half of the NLRome of solanaceous plants. Given the prevalence of this NLR network, we hypothesized that pathogens convergently evolved to secrete effectors that target NRC activities. To test this, we screened a library of 167 bacterial, oomycete, nematode and aphid effectors for their capacity to suppress the cell death response triggered by the NRC-dependent disease resistance proteins Prf and Rpi-blb2. Among five of the identified suppressors, one cyst nematode protein and one oomycete protein suppress the activity of autoimmune mutants of NRC2 and NRC3, but not NRC4, indicating that they specifically counteract a subset of NRC proteins independently of their sensor NLR partners. Whereas the cyst nematode effector SPRYSEC15 binds the nucleotide-binding domain of NRC2 and NRC3, the oomycete effector AVRcap1b suppresses the response of these NRCs via the membrane trafficking-associated protein NbTOL9a (Target of Myb 1-like protein 9a). We conclude that plant pathogens have evolved to counteract central nodes of the NRC immune receptor network through different mechanisms. Coevolution with pathogen effectors may have driven NRC diversification into functionally redundant nodes in a massively expanded NLR network.

## INTRODUCTION

Our view of the pathogenicity mechanisms of plant parasites (i.e. pathogens and pests) has significantly broadened over the years. Parasites as diverse as bacteria, oomycetes, nematodes and aphids turned out to be much more sophisticated manipulators of their host plants than anticipated. Indeed, it is now well established that these parasites secrete an arsenal of proteins, termed effectors, that modulate plant responses, such as innate immunity, to enable host infection and colonization. As a consequence, deciphering the biochemical activities of effectors to understand how parasites successfully colonize and reproduce has become a major conceptual paradigm in the field of molecular plant pathology (1, 2). In fact, effectors have emerged as molecular probes that can be utilized to unravel novel components and processes of the host immune system (2).

Most effectors studied to date suppress immune pathways induced by pathogen-associated molecular patterns (PAMPs). This so-called PAMP or PRR-triggered immunity (PTI) response is mediated by cell-surface pattern recognition receptors (3, 4) (Figure S1A). A subset of effectors, however, have avirulence (AVR) activity and inadvertently activate intracellular immune receptors of the nucleotide-binding, leucine-rich repeat (NLR) class of proteins, a response known as effector- or NLR-triggered immunity (ETI/NTI) (5-7). In plants, recognition of AVR effectors by NLRs can occur either directly or indirectly following various mechanistic models (8, 9). NTI is usually accompanied by a localised form of programmed cell death known as the hypersensitive response that hinders disease progression (6, 7, 10, 11). NLR-mediated immunity activated by AVR effectors can also be suppressed by other effectors (12). However, in sharp contrast to the widely studied PTI-suppressing effectors, the mechanisms by which effectors suppress NLR responses remain poorly understood (3, 13, 14). Therefore, understanding how effectors suppress NLR functions should provide important insights into the black box of how these immune receptors activate cell death and innate immunity, one of the major unsolved questions in the field of plant pathology (15).

NLRs are multidomain proteins with a NB-ARC (nucleotide-binding domain shared with APAF-1, various R-proteins and CED-4) and at least one additional domain (16). The C-terminus of plant NLRs is generally a leucine-rich repeat (LRR) domain, but they can be sorted into phylogenetic sub-groups with distinct N-terminal domains (16-18). In angiosperms, NLRs form several major phylogenetic sub-groups, including TIR-NLRs with an N-terminal Toll/interleukin-1 receptor (TIR) domain, CC-NLRs, with the Rx-type coiled-coil (CC) domain, CC_R_-NLRs with the RPW8-type CC (CC_R_) domain, and the CC_G10_-NLR subclade with a distinct type of CC (CC_A or_ CC_G10_) domain. Of these, CC-NLRs are the most common type, forming the largest group of NLRs in angiosperms (16, 18-20). The TIR- and CC-type N-terminal domains are involved in downstream immune signalling, oligomerization and cell death execution (21). The central NB-ARC domain exhibits ATP binding and hydrolysis activities and functions as a molecular switch that determines the NLR inactive/active status (22). Finally, the LRR domain can mediate effector perception and often engages in intra-molecular interactions (8, 22).

Plant genomes may encode anywhere between 50 and ∼1000 NLRs (17, 23). Some of these NLRs are functional singletons operating as single biochemical units, while others function in pairs or in more complex receptor networks (24). Paired and networked NLRs consist of functionally specialized sensor NLRs that detect pathogen effectors and helper NLRs that translate this effector recognition into hypersensitive cell death and resistance. In the Solanaceae, a major phylogenetic clade of NLRs forms a complex immunoreceptor network in which multiple helper CC-NLRs, known as NRCs (NLR required for cell death), are necessary for a large number of sensor NLRs (25). These sensor NLRs, encoded by R gene loci, confer resistance against diverse pathogens, such as viruses, bacteria, oomycetes, nematodes and insects (25) (Figure S1B). Together, NRCs and their NLR partners form the NRC superclade, a well-supported phylogenetic cluster divided into an NRC helper clade and two larger clades that include all known NRC-dependent sensor NLRs (25, 26). The NRC superclade emerged over 100 million years ago (Mya) from an NLR pair that expanded to make up a significant fraction of the NLRome of asterid plant species (25). The current model is that NRCs form redundant central nodes in this massively diversified bow-tie network, with different NRCs exhibiting some specificity towards their sensor NLR partners. For example, whereas the bacterial resistance protein Prf requires NRC2 and NRC3 in a genetically redundant manner, the potato blight resistance protein Rpi-blb2 is dependent only on NRC4 (25, 27) (Figure S1B).

Until very recently, the molecular mechanisms that underpin plant NLR activation and the subsequent execution of NTI pathways and hypersensitive cell death have remained unknown. The recent elucidation of the CC-NLR ZAR1 (HopZ-activated resistance 1) and the TIR-NLRs ROQ1 (Recognition of XopQ 1) and RPP1 (Recognition of *Peronospora parasitica* 1) structures have resulted in a new conceptual framework (28-31). Activated NLRs oligomerize into a wheel-like multimeric complex known as the resistosome. In the case of ZAR1, activation induces conformational changes in the NB-ARC domain resulting in ADP release followed by dATP/ATP binding and oligomerization of ZAR1 and its partner host proteins into the pentameric resistosome structure (29, 32). ZAR1 resistosomes expose a funnel-shaped structure formed by the N-terminal α1 helices, which was proposed to perturb plasma membrane integrity to trigger hypersensitive cell death (29, 30). This α1 helix is defined by a molecular signature, the MADA motif, that is present in ∼20% of angiosperm CC-NLRs, including NRCs (26). This implies that the biochemical ‘death switch’ mechanism of the ZAR1 resistosome probably applies to NRCs and other MADA motif containing NLRs across diverse plant taxa.

The NRC superclade is massively expanded in solanaceous plants, where it comprises up to half of the NLRome in some species (25). Given this prevalence, we hypothesized that pathogens convergently evolved to secrete effectors that target the NRC network. In this study, we screened collections of effector proteins from six major parasites of solanaceous plants, i.e. the bacterium *Pseudomonas syringae*, the oomycete *Phytophthora infestans*, the cyst nematodes *Globodera rostochiensis* and *Globodera pallida*, and the aphids *Myzus persicae* and *Acrythosiphon pisum*, for their capacity to suppress the hypersensitive cell death triggered by the NRC-dependent sensor NLRs Prf and Rpi-blb2. These screens revealed five effectors as suppressors of components of the NRC network. Among these, the oomycete protein PITG-16705 (PBHR_012, henceforth, AVRcap1b) (33) and the cyst nematode protein SPRYSEC15 (where the term SPRYSEC will be referred to as SS from here on in) stood out for being able to specifically suppress the response triggered by autoimmune mutants of NRC2 and NRC3, but not NRC4, indicating that they are able to counteract a subset of NRC helper proteins independently of their sensor NLR partners. Further studies revealed that AVRcap1b and SS15 suppress NRC2 and NRC3 through different mechanisms. While AVRcap1b suppression of NRC2 and NRC3 mediated immunity is dependent on the membrane trafficking-associated protein NbTOL9a (Target of Myb 1-like protein 9a), SS15 directly binds the NB-ARC domain of NRC2 and NRC3 to perturb their function. We conclude that evolutionarily divergent plant pathogens have convergently evolved distinct molecular strategies to counteract central nodes of the NRC immune receptor network. Coevolution with pathogen effectors may, therefore, underpin the diversification of NRCs as functionally redundant nodes in a massively expanded NLR network.

## RESULTS

### *In planta* screens reveal pathogen effectors that suppress NRC-mediated hypersensitive cell death

To determine the extent to which pathogens have evolved to target the NRC network, we screened candidate effectors from six diverse pathogen species for their ability to suppress the hypersensitive cell death triggered by the disease resistance proteins Prf (responds to Pto/AvrPto; NRC2/3 dependent) and Rpi-blb2 (responds to AVRblb2; NRC4 dependent) (25, 27). The effectors and empty vector (EV, control) were transiently expressed with Pto/AvrPto or Rpi-blb2/AVRblb2 in leaves of the model experimental plant *Nicotiana benthamiana* and assessed for the absence of cell death, which is indicative of a suppression phenotype (Figure 1A). Our screen included 167 effector candidates from oomycetes (*Phytophthora infestans*, 65), bacteria (*Pseudomonas syringae*, 26), cyst nematodes (*Globodera rostochiensis*, 23 and *Globodera pallida*, 3), and aphids (*Myzus persicae*, 47 and *Acrythosiphon pisum*, 3). Among the 167 effectors tested, two effectors (AVRcap1b, SS15) suppressed Prf-mediated cell death and three effectors (PITG-15278, SS10 and SS34) suppressed Rpi-blb2-mediated cell death (Figure 1B, Figure S2). AVRcap1b and PITG-15278 are RXLR-WY/LWY domain containing effectors from *P. infestans* (34, 35). SS10, SS15, SS34 are SPRY domain containing effectors from *G. rostochiensis* (36-38) (Figure 2A; Table S1). Interestingly, HopAB2 (also known as AvrPtoB), which is well known to suppress Prf-mediated cell death (39), was not a robust suppressor in our assays, and only suppressed Prf-mediated cell death in older leaves of *N. benthamiana* (Figure S3), leaf age as defined by (40).

**Figure 1.**
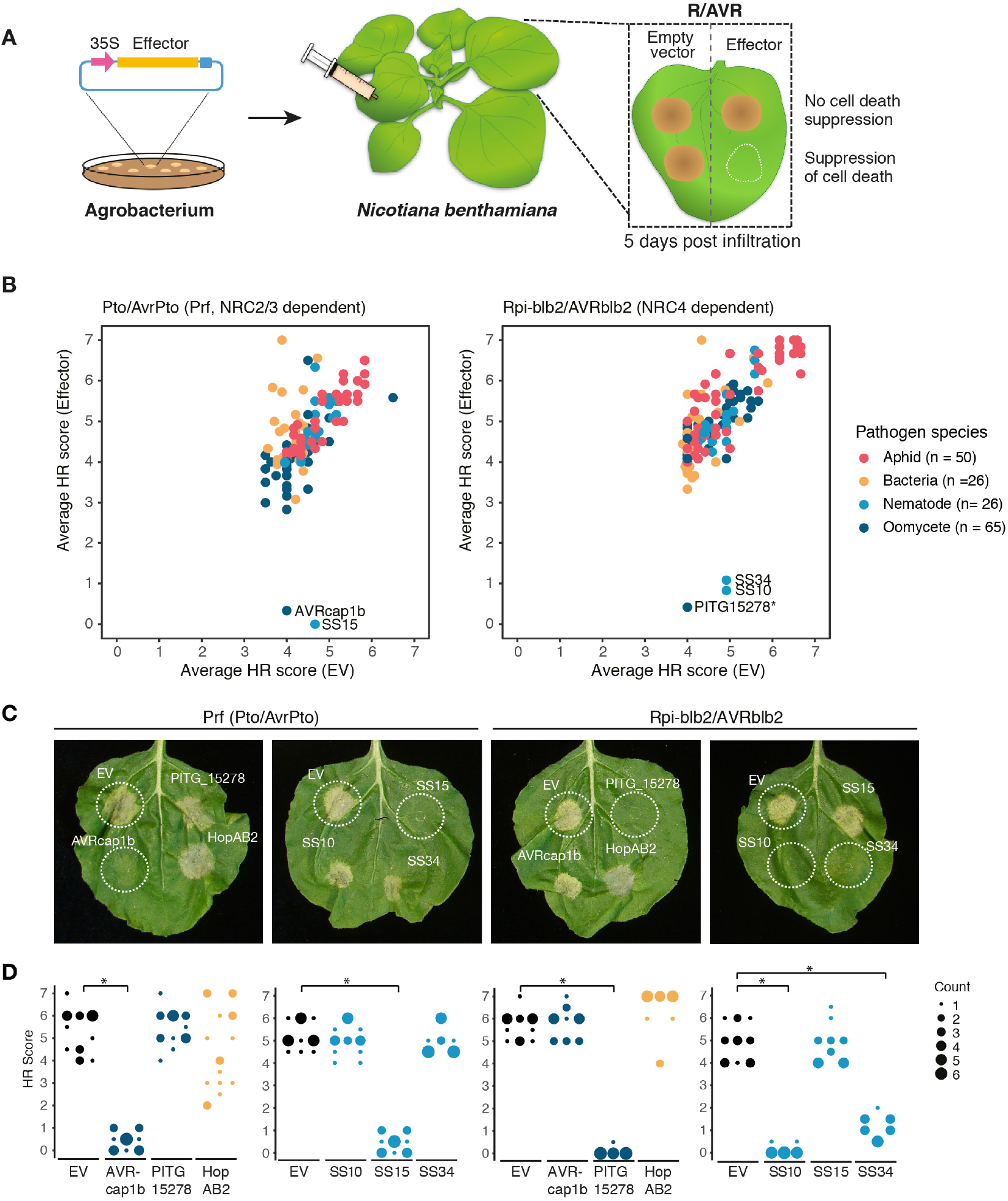
Five effectors suppress Prf-(NRC2/3 dependent) or Rpi-blb2-(NRC4 dependent) mediated cell death. (A) A schematic representation of cell death assay strategy used. Effectors and Empty Vector (EV) were transformed into *A. tumefaciens*, and transiently co-expressed in *N. benthamiana* plants with either Pto/AvrPto or Rpi-blb2/AVRblb2. EV was used as a negative control. Hypersensitive Response (HR) was scored based on a modified 0 - 7 scale (Segretin et al 2014) between 5 – 7 days post infiltration. (B) A scatterplot of the average HR score of EV versus the average HR score of each tested effector (n=167); Pto/AvrPto (left panel) Rpi-blb2/AVRblb2 (right panel). Effectors that have suppression activity are represented as outliers within the plot. Results are based on six technical replicates. (C) Representative *N. benthamiana* leaves infiltrated with appropriate constructs photographed 5 – 7 days after infiltration. The R/AVR pair tested, Prf (Pto/AvrPto) or Rpi-blb2/AVRblb2, are labeled above the leaf panel and effectors labeled on leaf image. (D) HR was scored 5 days post-agroinfiltration. The results are presented as a dot plot, where the size of each dot is proportional to the number of samples with the same score (count) within each biological replicate. The experiment was independently repeated three times with six technical replicates. The columns for either EV or for each individual effector correspond to results from different biological replicates. Dot colours represent pathogen species as indicated in (B). Significant differences between the conditions are indicated with an asterisk (*). The details of statistical analysis are presented in Figure S2.

**Figure 2.**
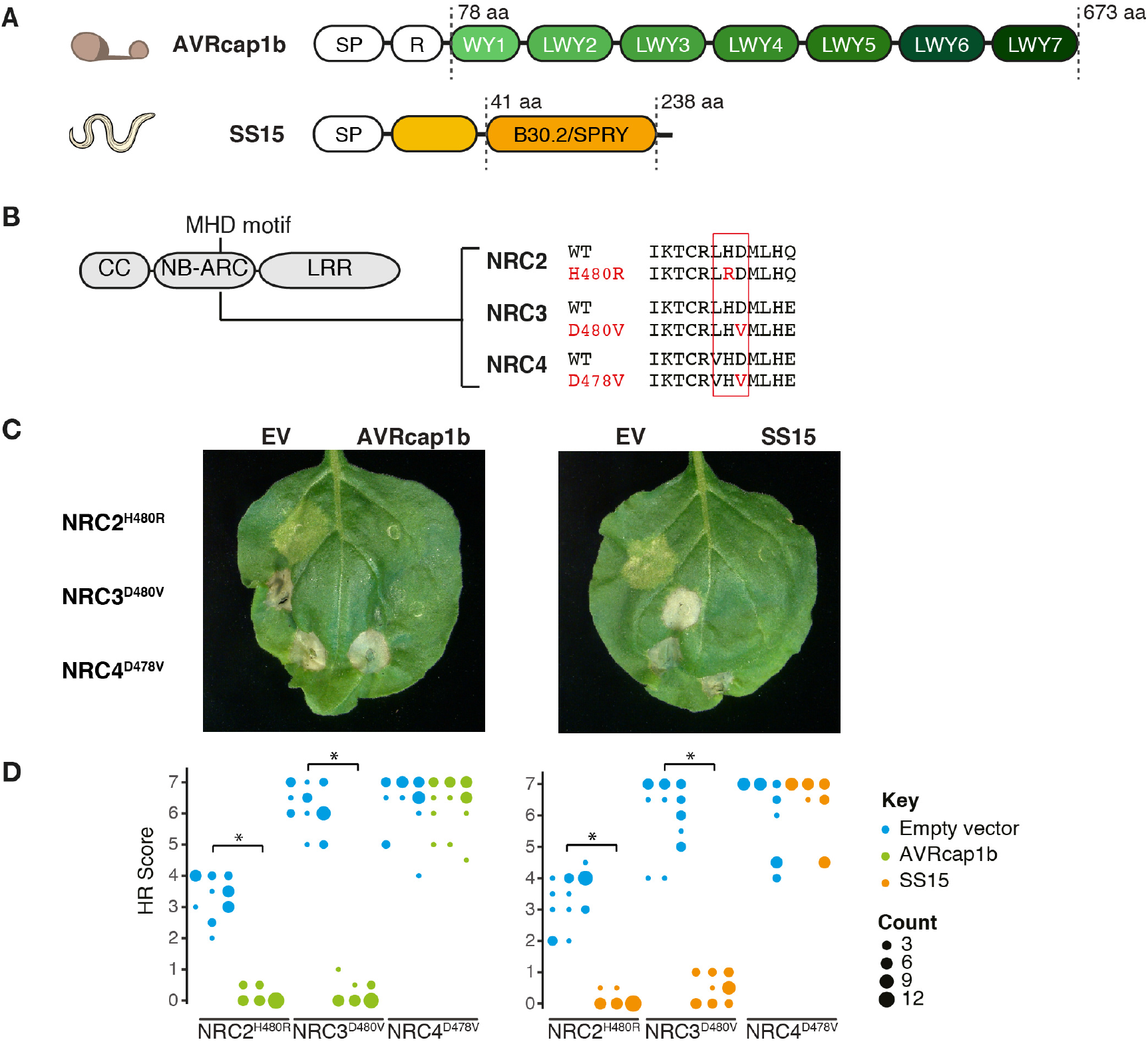
AVRcap1b and SS15 suppress cell death activity mediated by autoactive NRC2^H480R^ and NRC3^D480V^ but not NRC4^D478V^. (A) Domain organization of the RXLR-WY/LWY domain containing *P. infestans* effector AVRcap1b and the B30.2/SPRY (IPR001870) domain containing *G. rostochiensis* effector SS15. (B) Schematic representation of NRC2, NRC3 and NRC4 and the mutated sites in the MHD motif. Substituted residues are shown in red in the multiple sequence alignment. (C) Representative *N. benthamiana* leaves showing HR after co-expression of Empty vector, AVRcap1b or SS15, indicated above leaf panels, with autoimmune NRC2^H480R^, NRC3^D480V^, and NRC4^D478V^ mutants. Plants were photographed 5 days after agroinfiltration. (D) HR was scored 5 days post-agroinfiltration. The results are presented as a dot plot, where the size of each dot is proportional to the number of samples with the same score (count) within each biological replicate. The experiment was independently repeated three times each with six technical replicates. The columns of each tested condition (labelled on the bottom of the plot) correspond to results from different biological replicates. Significant differences between the conditions are indicated with an asterisk (*). The details of statistical analysis are presented in Figure S4.

### AVRcap1b and SS15 effectors suppress the cell death triggered by autoimmune mutants of NRC2 and NRC3 but not NRC4

To ascertain whether the identified effectors suppress the activity of the sensor NLRs (Prf or Rpi-blb2) or the underlying helper NLRs (NRC2, NRC3, NRC4), we generated constitutively active (autoimmune) NRCs by mutating the MHD motif resulting in the NRC autoactive variants NRC2^H480R^, NRC3^D480V^ and NRC4^D478V^ (41) (Figure 2B). When transiently expressed in *N. benthamiana*, all three NRC mutants caused cell death in the absence of a pathogen AVR effector (Figure 2C). Next, we co-expressed the suppressors AVRcap1b, SS15, PITG-15278, SS10 and SS34, along with the EV control, with NRC2^H480R^, NRC3^D480V^ and NRC4^D478V^ in *N. benthamiana* leaves (Figure S4). PITG-15278, SS10 and SS34 did not suppress any of the autoactive NRCs, indicating that they target the sensor NLR Rpi-blb2 or the interaction between Rpi-blb2 and NRC4 (Figure S4). Interestingly, both AVRcap1b and SS15 suppressed the cell death induced by autoimmune NRC2 and NRC3, therefore recapitulating their specific suppression of the sensor NLRs Prf (Figure 2C-D, Figure S5). In contrast, AVRcap1b and SS15 didn’t affect the autoactivity of NRC4 (Figure 2C-D, Figure S5). These results indicate that the suppression activity of AVRcap1b and SS15 is independent of the sensor NLR Prf and is specific to NRC2 and NRC3. We conclude that AVRcap1b and SS15 are likely acting on the NRC2 and NRC3 helper nodes of the NRC network.

### AVRcap1b and SS15 effectors suppress multiple disease resistance proteins (sensor NLRs) that signal through NRC2 and NRC3

To further challenge our finding that AVRcap1b and SS15 target NRC2 and NRC3 but not NRC4, we screened the two effectors for suppression of six disease resistance sensor NLRs (SW5b, Gpa2, R1, Rx, Bs2 and R8) that were previously assigned to the NRC network (25, 26). We transiently co-expressed AVRcap1b and SS15 with these NRC-dependent sensor NLRs in addition to the previously tested Prf and Rpi-blb2, in *N. benthamiana* leaves using agroinfiltration. Of these, five NLRs were co-expressed with their cognate AVR effectors, whereas we used a constitutively active version of SW5b (SW5b^D857V^). We also included Rpi-vnt1 as an NRC-independent NLR protein negative control (25). Interestingly, while SW5b was previously shown to signal through NRC2, NRC3 and NRC4 in a redundant manner (25), SW5b^D857V^ signalled through NRC2 and NRC3 only. These experiments revealed that AVRcap1b and SS15 suppress SW5b^D857V^ and Gpa2 in addition to Prf, but none of the other five NLRs (Rpi-blb2, R8, R1, Rx, Bs2). Additionally, neither AVRcap1b nor SS15 suppressed the cell death mediated by the NRC-independent NLR Rpi-vnt1 (Figure 3, Figure S6). These results are consistent with the model that AVRcap1b and SS15 suppress NRC2 and NRC3 but not NRC4 given that, similar to Prf, both SW5b^D857V^ and Gpa2 are dependent on NRC2 and NRC3, whereas the other tested NLRs signal through NRC4 specifically or redundantly with NRC2 and NRC3 (25, 26).

**Figure 3.**
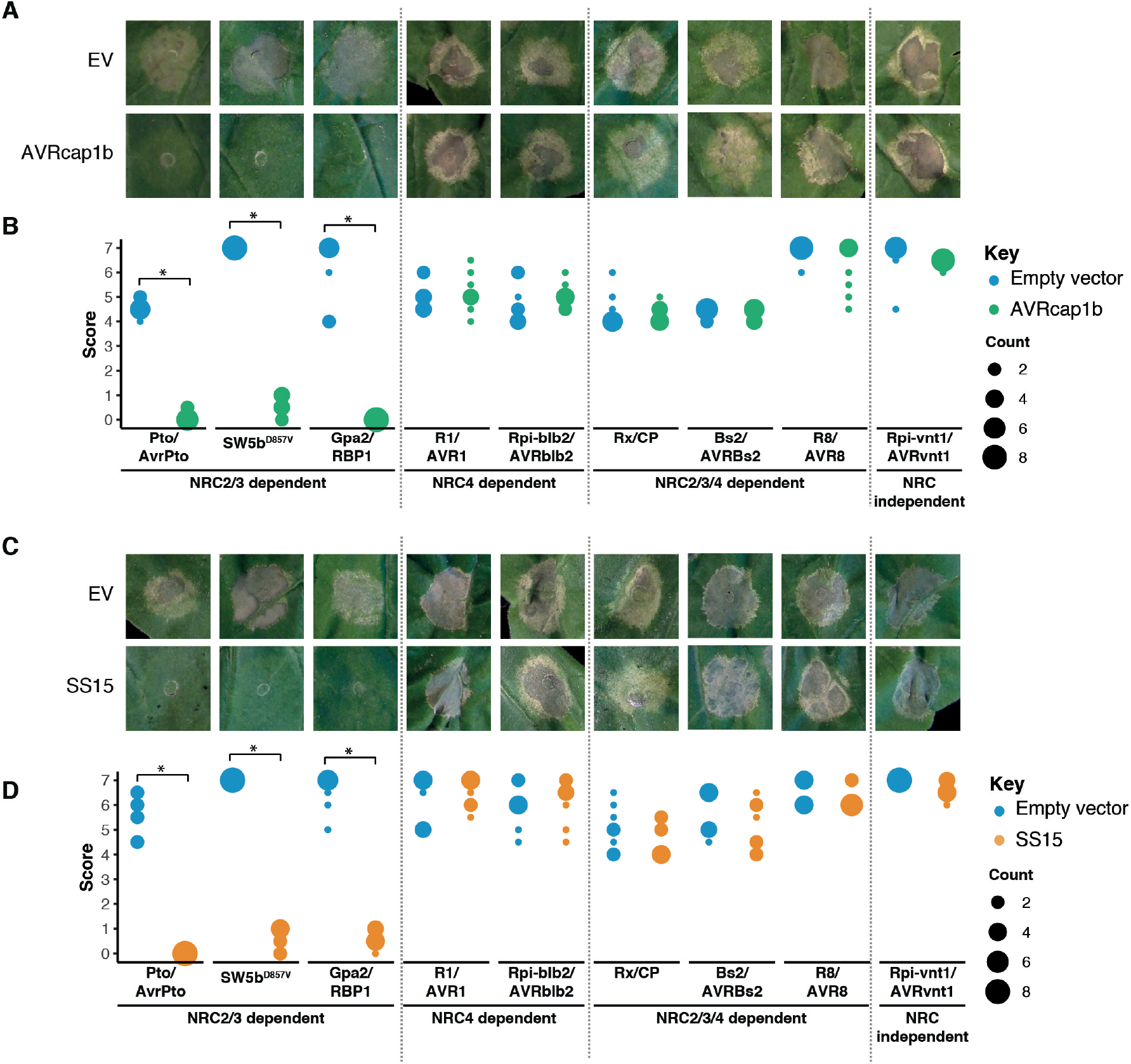
AVRcap1b and SS15 suppress cell death mediated by sensor NLRs that signal through NRC2 and NRC3 but not NRC4. Representative leaf panels showing HR phenotypes of (A) Empty vector (EV) and AVRcap1b or (C) EV and SS15 co-infiltrated with a range of sensor NLRs (labeled on the bottom of panels B and D). Photographs were taken 5 days post-agroinfiltration. (B, D) HR results are presented as dot plots, where the size of each dot is proportional to the number of samples with the same score (count). A total of eight technical replicates was completed for each treatment. Significant differences between the conditions are indicated with an asterisk (*). The details of statistical analysis are presented in Figure S6.

### AVRcap1b and SS15 suppress Rx-mediated cell death only in the absence of NRC4

The NRC dependent sensor NLR Rx, which confers extreme resistance to *Potato virus X* (PVX) by recognizing viral coat protein (CP), is dependent on NRC2, NRC3, and NRC4 in a genetically redundant manner (42-45). The genetic redundancy of NRCs may explain why the Rx-mediated cell death is not suppressed by AVRcap1b and SS15 in *N. benthamiana* (Figure 3). Since AVRcap1b and SS15 suppress the cell death activity of NRC2 and NRC3 but not NRC4, we reasoned that these two effectors should be able to suppress Rx-mediated cell death in the absence of NRC4. To challenge this hypothesis, we knocked-down NRCs in Rx-transgenic *N. benthamiana* plants using *Tobacco rattle virus* (TRV)-induced gene silencing, either individually (TRV:*NRC4*) or in combination (TRV:*NRC2/3*) (Figure 4A). Three weeks after inoculation with TRV, we co-expressed AVRcap1b or SS15 with CP in *NRC*-silenced leaves. These experiments showed that both AVRcap1b and SS15 suppress Rx-mediated cell death in *NRC4*-silenced leaves, but not in *NRC2/3*-silenced leaves nor in the TRV:EV control (Figure 4B-E, Figure S7). These results further validate our earlier finding that AVRcap1b and SS15 can specifically suppress the activities of NRC2 and NRC3 but not NRC4.

**Figure 4.**
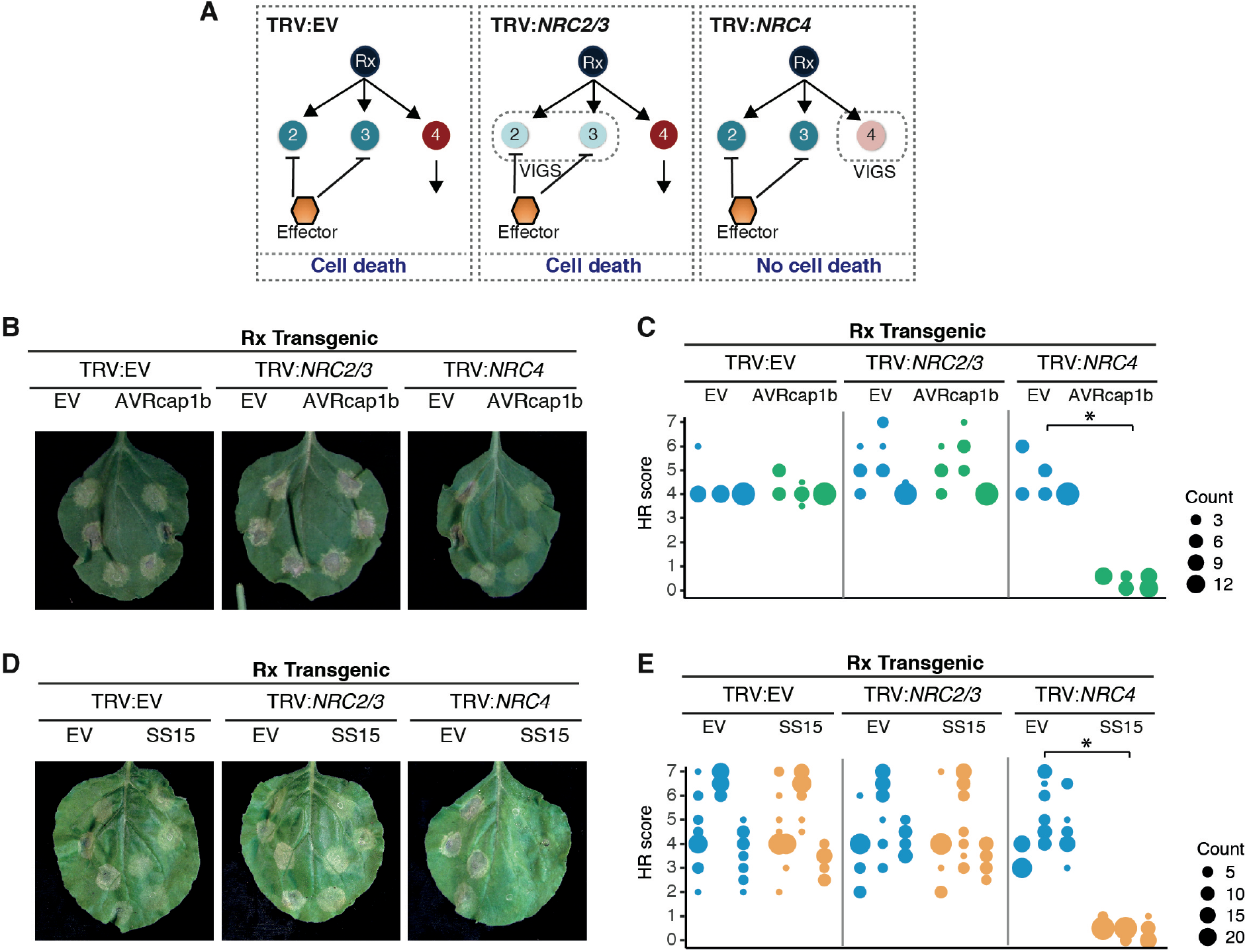
AVRcap1b and SS15 suppress Rx-mediated cell death in *NRC4*-silenced Rx transgenic *Nicotiana benthamiana* plants. (A) A schematic representation of the silencing and infiltration strategy undertaken. Photo of representative *N. benthamiana* leaves showing HR after co-expression of (B) Empty vector (EV) or AVRcap1b, and (D) EV or SS15 in TRV:EV, TRV:*NRC2/3* and TRV:*NRC4* silenced Rx-transgenic *N. benthamiana* plants, where EV was used as a negative control. Silencing treatments (TRV:EV, TRV:*NRC2/3*, TRV:*NRC4*) are indicated above leaf panels. Plants were photographed 5 days after agroinfiltration. Cell death response was scored 5 days after agroinfiltration (C) EV and AVRcap1b and (E) EV and SS15 and presented as dot plots, where dot colour represents either EV (blue), AVRcap1b (green) or SS15 (orange) and dot size is proportional to the number of samples with the same score (count). The experiment was independently repeated three times. Each replicate is represented by different columns within each silencing treatment for either EV, AVRcap1b or SS15. Significant differences between the conditions are indicated with an asterisk (*). The details of statistical analysis are presented in Figure S7.

### AVRcap1b and SS15 compromise Rx-mediated extreme resistance to *Potato virus X* in an NRC2 and NRC3 but not NRC4-dependent manner

We previously showed that co-silencing of *NRC2, NRC3* and *NRC4* not only compromised Rx-mediated hypersensitive cell death but also abolished extreme resistance to PVX, leading to trailing necrotic lesions indicative of virus spread (25). To determine the extent to which suppression mediated by AVRcap1b and SS15 translates into reduced viral infection, we tested the effectors ability to compromise Rx-mediated resistance to PVX. We transiently expressed AVRcap1b or SS15 in Rx transgenic *N. benthamiana* plants that were silenced for NRCs either individually (TRV:*NRC4*) or in combination (TRV:*NRC2/3*, TRV:*NRC2/3/4*). We then infected the leaves by agroinfection with *A. tumefaciens* carrying PVX (pGR106::PVX::GFP) using a toothpick inoculation method and documented the formation of trailing necrotic lesions at the inoculated spots (materials and methods) (25, 43, 46) (Figure 5A). Consistent with previous experiments (25), we observed trailing necrosis in the *NRC* triple silenced Rx leaves (TRV:*NRC2/3/4*) regardless of the presence or absence of AVRcap1b or SS15 (Figure 5B-C). We also observed trailing necrosis when AVRcap1b or SS15 was expressed in NRC4-silenced (TRV:*NRC4*) Rx leaves, indicating that both effectors compromised Rx-mediated resistance when NRC4 is depleted (Figures 5B-C). In contrast, Rx-mediated resistance to PVX was not compromised by AVRcap1b or SS15 in *NRC2/3*-silenced (TRV:*NRC2/3*) leaves. Virus accumulation in the different treatments was validated by western blot detection of GFP protein driven by the sub-genomic promoter of PVX::GFP (Figures 5D-E). Perturbation of PVX resistance was more markedly affected by SS15 compared to AVRcap1b based on lesion size and PVX accumulation (Figure 5C-E). These results indicate that the AVRcap1b and SS15 effectors not only suppress NLR-mediated cell death but also counteract the disease resistance phenotype mediated through NRC2 and NRC3.

**Figure 5.**
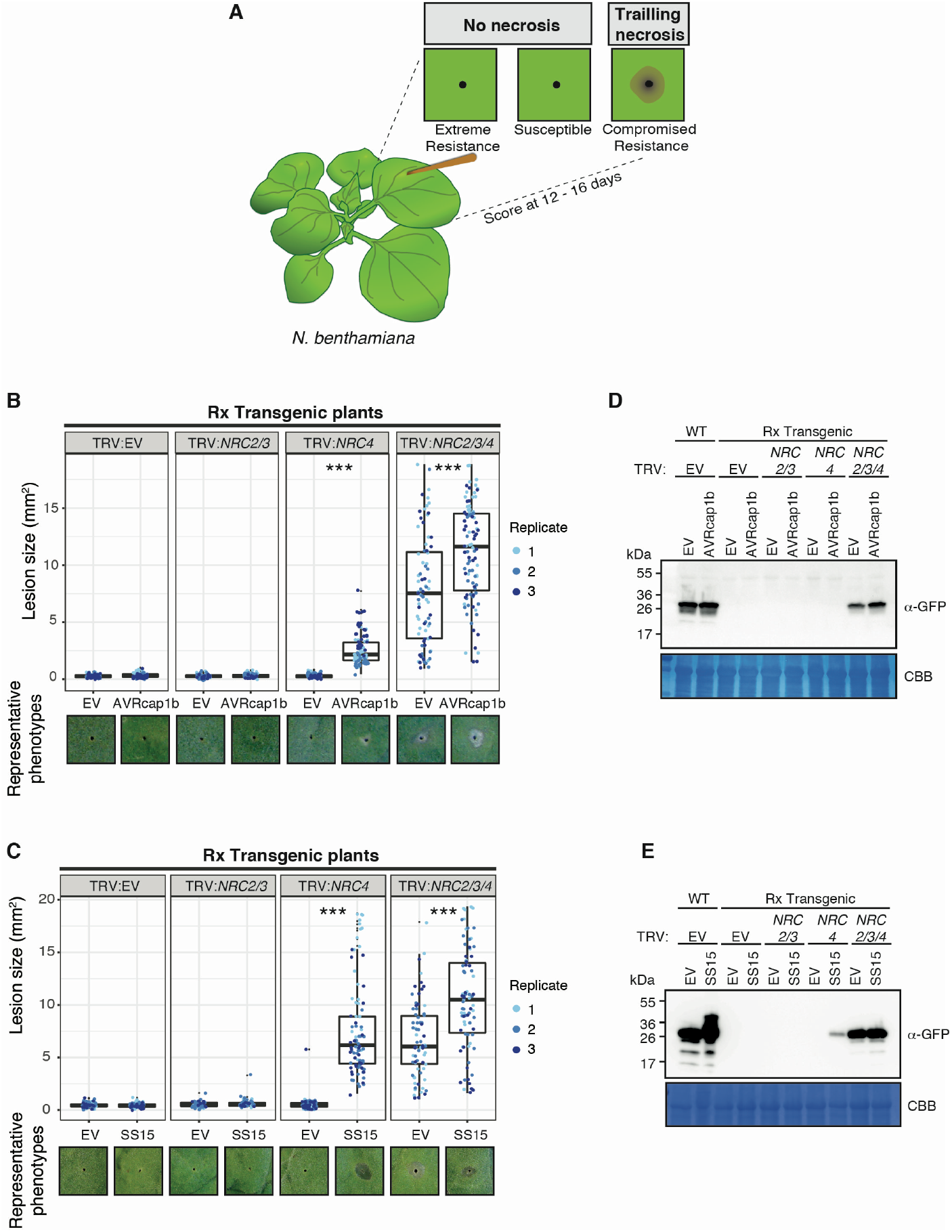
AVRcap1b and SS15 compromise Rx-mediated extreme resistance to PVX in NRC4 silenced plants. (A) Toothpick inoculation method previously described by Wu et al 2017, allowed examination of the spread of trailing necrotic lesions due to the partial resistance mediated by Rx. Empty vector (EV), NRC2, NRC3, or NRC4 were silenced individually or in combination in Rx transgenic *N. benthamiana* plants by TRV. (B) EV or AVRcap1b and (C) EV or SS15 were expressed, via agroinfiltration, in leaves of the NRC silenced plants one day before PVX inoculation. PVX-GFP (pGR106-GFP) was inoculated using the toothpick inoculation method, as per panel (A). Photographs were taken 12 – 16 days after PVX inoculation. The size of the necrotic lesions was measured using Fiji (previously Image J). Data acquired from different biological replications are presented in different colours. Statistical differences among the samples were analysed with mixed model ANOVA and a Tukey’s HSD test (*p-*value < 0.01), where the fixed effect is the silencing treatment (TRV:EV, TRV:*NRC2/3*, TRV:*NRC4*, TRV:*NRC2/3/4*) with either EV or AVRcap1b and the random effect is the experimental replicate.. Significant differences between the conditions are indicated with asterisks (***, *p-*value <0.0001). Representative phenotypes observed for each treatment are presented below the boxplots. Immunoblot analysis of GFP accumulation of pGR106-GFP toothpick inoculated sites in the presence of (D) AVRcap1b and (E) SS15 in Rx transgenic or Wild Type (WT) *N. benthamiana* plants.

### Yeast two-hybrid screens reveal candidate host interactors of the AVRcap1b and SS15 effectors

To investigate how AVRcap1b and SS15 suppress NRC activities, we set out to identify their host interactors. We used the effectors as baits in unbiased yeast two-hybrid (Y2H) screens against a *N. benthamiana* mixed tissue cDNA library (ULTImate Y2H, Hybrigenics Services, Paris, France). AVRcap1b was screened against a combined ∼140 million clones, which resulted in 13 candidate interacting proteins from a total of 35 positive clones (Table S2). SS15 was screened against ∼61 million clones resulting in 10 candidate proteins from a total of 202 positive clones (Table S3). Remarkably, NRC3 and NRC4 were among the recovered SS15 protein interactors, which is notable given that NLR proteins are rarely recovered from Y2H screens. The NRC3 and NRC4 fragments that were recovered from the Y2H screen matched the CC-NB-ARC domains indicating that SS15 may bind the N-terminal half of NRC proteins (Figure 6A; Table S3).

**Figure 6.**
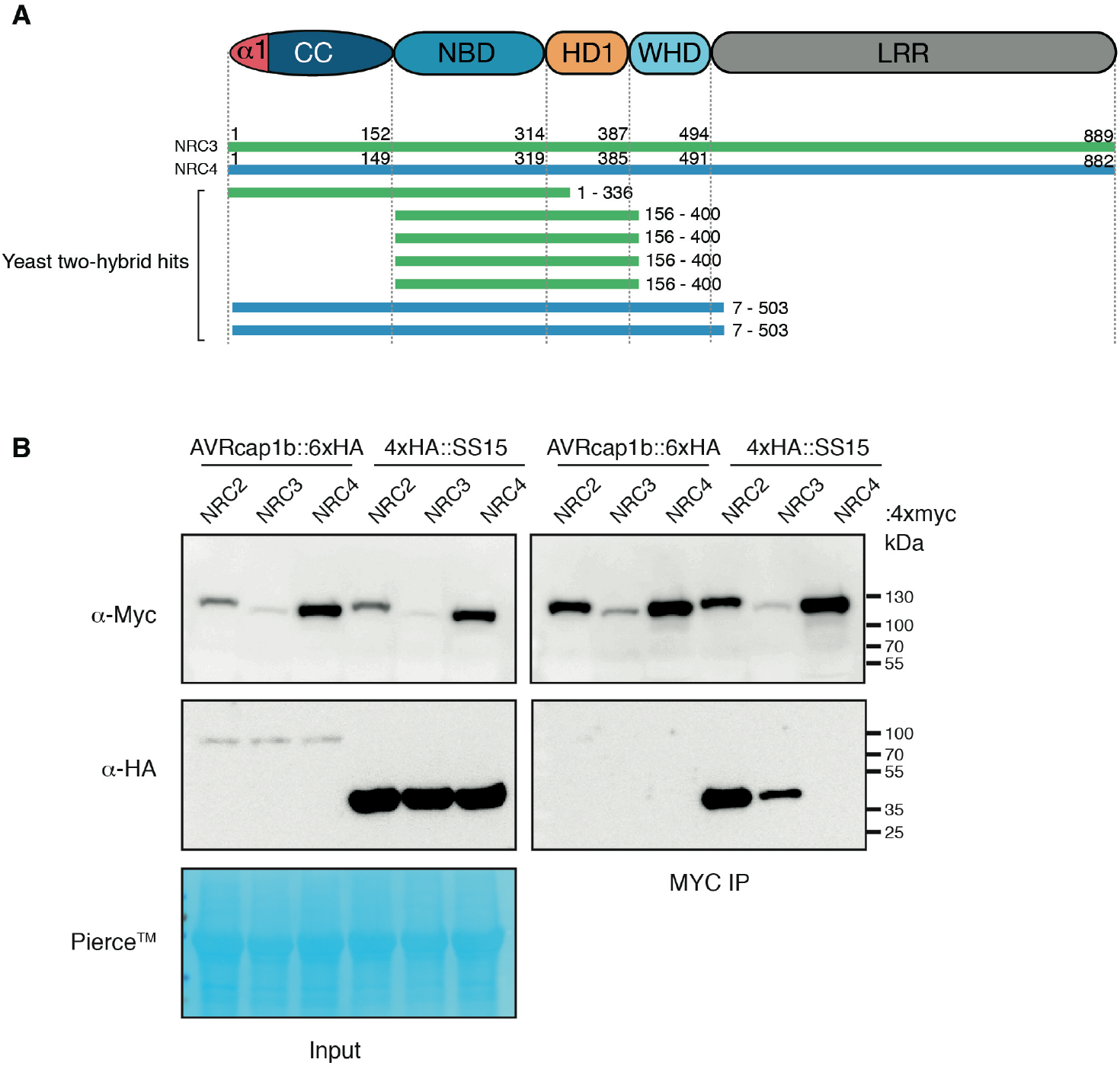
SS15, but not AVRcap1b associates with NRCs *in planta*. (A) Schematic diagram of domain organization of NRC proteins, showing yeast two-hybrid hits from the Hybrigenics services screen (Table S3). (B) Co-immunoprecipitation experiment of C-terminally 4xmyc tagged NRC2, NRC3 and NRC4 with C-terminally HA tagged AVRcap1b::6xHA and N-terminally tagged 4xHA:SS15 (labelled above). Proteins obtained by co-immunoprecipitation with MYC beads (MYC IP) and total protein extracts (input) were immunoblotted with the appropriate antisera labelled on the left. Approximate molecular weights (kDa) of the proteins are shown on the right. Rubisco loading control was carried out using Pierce™ staining. The experiment was performed more than three times under different pulldown conditions with similar results.

To further investigate the interactions between SS15 and NRCs, we sought out to delimit the binding domain within the NLRs. Since the prey fragment hits from the Y2H screen covered the CC-NB-ARC domain of NRC3 and NRC4 (Figure 6A), we generated CC and NB-ARC domain truncations and tested them in additional Y2H experiments for interaction with SS15. These assays revealed that SS15 binds to the NB-ARC but not the CC domains of NRC2, NRC3 and NRC4 (Figure S8). Consistent with earlier observations, AVRcap1b did not interact with either the CC or NB-ARC domains of the NRCs in these Y2H experiments (Figure S8).

### Unlike AVRcap1b, SS15 associates with NRC2 and NRC3 *in planta*

The Y2H results prompted us to investigate the association between AVRcap1b, SS15 and NRCs using co-immunoprecipitation (coIP) of proteins expressed in *N. benthamiana*, which should reveal the association between the effectors and full-length NRC proteins in a more physiologically relevant condition. To achieve this, we co-expressed each of AVRcap1b::6xHA and 4xHA::SS15 with NRC2::4xMyc, NRC3::4xMyc and NRC4::4xMyc in *N. benthamiana* leaves using agroinfiltration, and subjected the protein extracts to anti-Myc immunoprecipitation and western blot analyses. These experiments revealed that SS15 co-immunoprecipitated with NRC2 and NRC3, but not with NRC4 (Figure 6B). Occasionally, a weak NRC4 signal was detected in some biological replicates, suggesting that SS15 may also associate with NRC4 but with markedly lower affinity compared to NRC2 and NRC3 (Figure S9). Consistent with the findings of the Y2H screens, we did not detect an association between AVRcap1b and any of the three NRCs in these coIP experiments (Figure 6B).

### SS15 directly binds the NB-ARC domain of NRC3 *in vitro*

To further examine the association between SS15 and the NB-ARC domains of NRCs, we purified these proteins for *in vitro* assays using *Escherichia coli* as a heterologous expression system. We successfully obtained homogeneous SS15 and NRC3 NB-ARC domain (NRC3^NB-ARC^) proteins but were unable to obtain purified NRC2 NB-ARC or NRC4 NB-ARC domains due to solubility and stability issues. We subjected purified SS15 and NRC3^NB-ARC^ to gel filtration and found that these proteins elute at 239 ml and 227 ml, which correspond to 42.52 and 27.75 kDa respectively (Figure 7). To determine whether the two proteins form a complex *in vitro*, we mixed cells expressing individual proteins and co-purified SS15 and NRC3^NB-ARC^ for gel filtration assays (Materials and methods). The co-purified mixture of SS15 and NRC3^NB-ARC^ resulted in a peak shift with an elution volume at 211 ml, which corresponds to 84.02 kDa. Further validation by SDS-PAGE of the fractions under the new peak confirmed the presence of both proteins (Figure 7). In our gel filtration assays, the protein molecular weight calibration led to overestimates of the predicted molecular masses of the proteins, both alone and in complex (NRC3^NB-ARC^ is 41 kDa, SS15 is 24.5 kDa, and NRC3^NB-ARC^ - SS15 complex is 65.5 kDa). However, the results indicate that monomeric forms of each state exist in solution and that the two proteins probably enter in a 1:1 complex under these experimental conditions. Taken together, these results suggest that SS15 forms a complex with NRC^NB-ARC^ *in vitro*.

**Figure 7.**
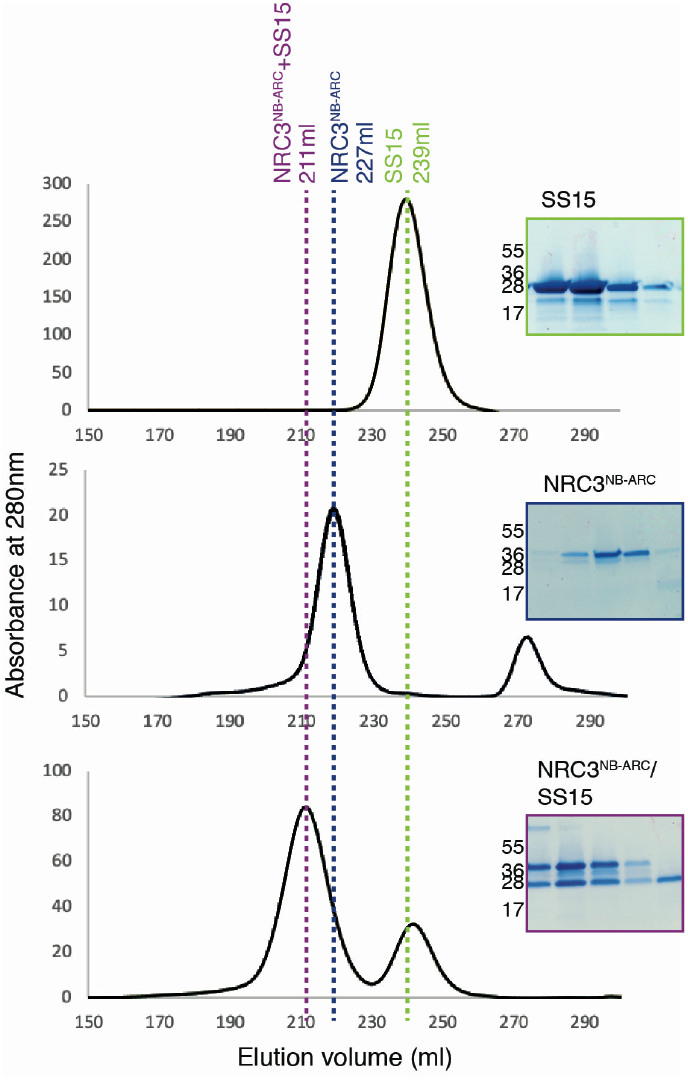
SS15 binds the NB-ARC domain of NRC proteins *in vitro*. SS15 binds the NRC3^NB-ARC^ domain *in vitro*. Gel filtration traces obtained for SS15 (top), NRC3^NB-ARC^ domain (middle) and a 1:1 Mixture of the complex (bottom). Insets show SDS-PAGE gels of the fractions collected across the elution peaks.

### SS15 associates with p-loop mutants of NRC2 and NRC3

The p-loop motif within the NB-ARC domain of NLR proteins is crucial for ATP binding and hydrolysis, a biochemical step that is essential for NLR oligomerisation and activation (47). To determine whether SS15 associates with p-loop mutants of the NRCs, we performed coIP experiments in *N. benthamiana* with the NRC p-loop mutants NRC2^K188R^, NRC3^K191R^ and NRC4^K190R^ (25). These experiments revealed that SS15 associates with the p-loop mutants of NRC2 (NRC2^K188R^) and NRC3 (NRC3^K191R^) (Figure S10). Since an intact p-loop is not required for SS15-NRC association, our findings suggest that SS15 can probably enter in complex with inactive forms of NRC2 and NRC3.

### SS15 associates with activated forms of NRC2

We next investigated the extent to which SS15 associates with activated NRC2 and NRC3. We co-expressed 4xHA::SS15 with C-terminally 4xMyc tagged autoimmune forms of NRC2^H480R^ and NRC3^D480V^ in *N. benthamiana* using agroinfiltration and subjected the protein extracts to anti-Myc immunoprecipitation and western blot analyses. We found that SS15 co-immunoprecipitated with NRC2^H480R^, indicating that this effector associates with activated forms of NRC2 (Figure 8). Our results for NRC3^D480V^, however, were inconclusive, since this protein displayed poor accumulation and couldn’t be detected in western blot analyses under these conditions. Despite this, our overall conclusion is that SS15 binds both inactive and activated forms of NRC2.

**Figure 8.**
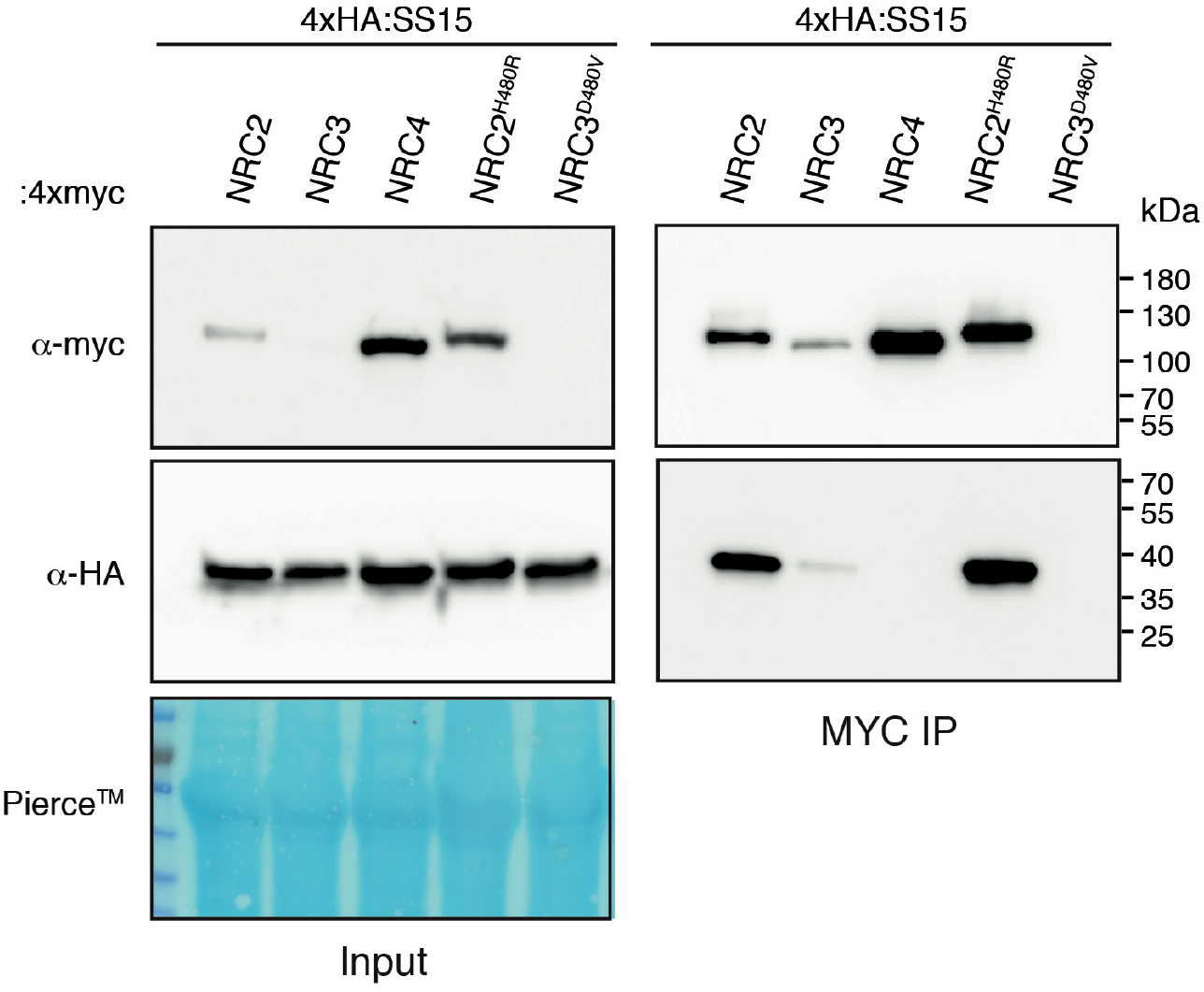
SS15 associates with autoactive NRC2. N-terminally 4xHA tagged SS15 was transiently co-expressed in *N. benthamiana* with C-terminally 4xMyc tagged NRC2, NRC3, NRC4, NRC2^H480R^, NRC3^D480V^ and NRC4^D478V^. Immunoprecipitation (IP) was performed with agarose beads conjugated to Myc (Myc IP) antibodies. Total protein extracts (Input) and proteins obtained by co-immunoprecipitation were immunoblotted with appropriate antisera labelled on the right. Approximate molecular weights (kDa) of the proteins are shown on the right. Rubisco loading controls were conducted using Pierce™ staining. This experiment is representative of two independent replicates.

### AVRcap1b associates with the *Nicotiana benthamiana* ENTH/VHS-GAT domain protein TOL9a *in planta*

Given that AVRcap1b did not interact with NRCs, we reasoned that this effector targets another host protein that is involved in NLR immunity. To narrow down the list of 13 candidate interactors obtained from the Y2H screen, we subjected AVRcap1b to *in planta* coIP coupled with tandem mass spectrometry (IP-MS) using methods that are well established in our laboratory (48-50). IP-MS experiments with GFP::AVRcap1b resulted in the identification of eight unique AVRcap1b interactors that were not recovered with the control GFP::PexRD54, another *P. infestans* effector of a similar fold and size (Figure S11; Table S4). Three of the eight interactors were different members of the Target of Myb 1-like protein (TOL) family of ENTH/VHS-GAT domain-containing proteins that function in membrane trafficking as part of the endosomal sorting complex required for transport (ESCRT) pathway (51, 52) (Figure 9A, Figure S10, Table S4). One of these candidate TOLs, Nbv6.1trP4361, was independently recovered in the Y2H screen (Table S2). Therefore, we decided to further investigate the corresponding protein (hereafter referred to as NbTOL9a) as a candidate host target of AVRcap1b.

**Figure 9.**
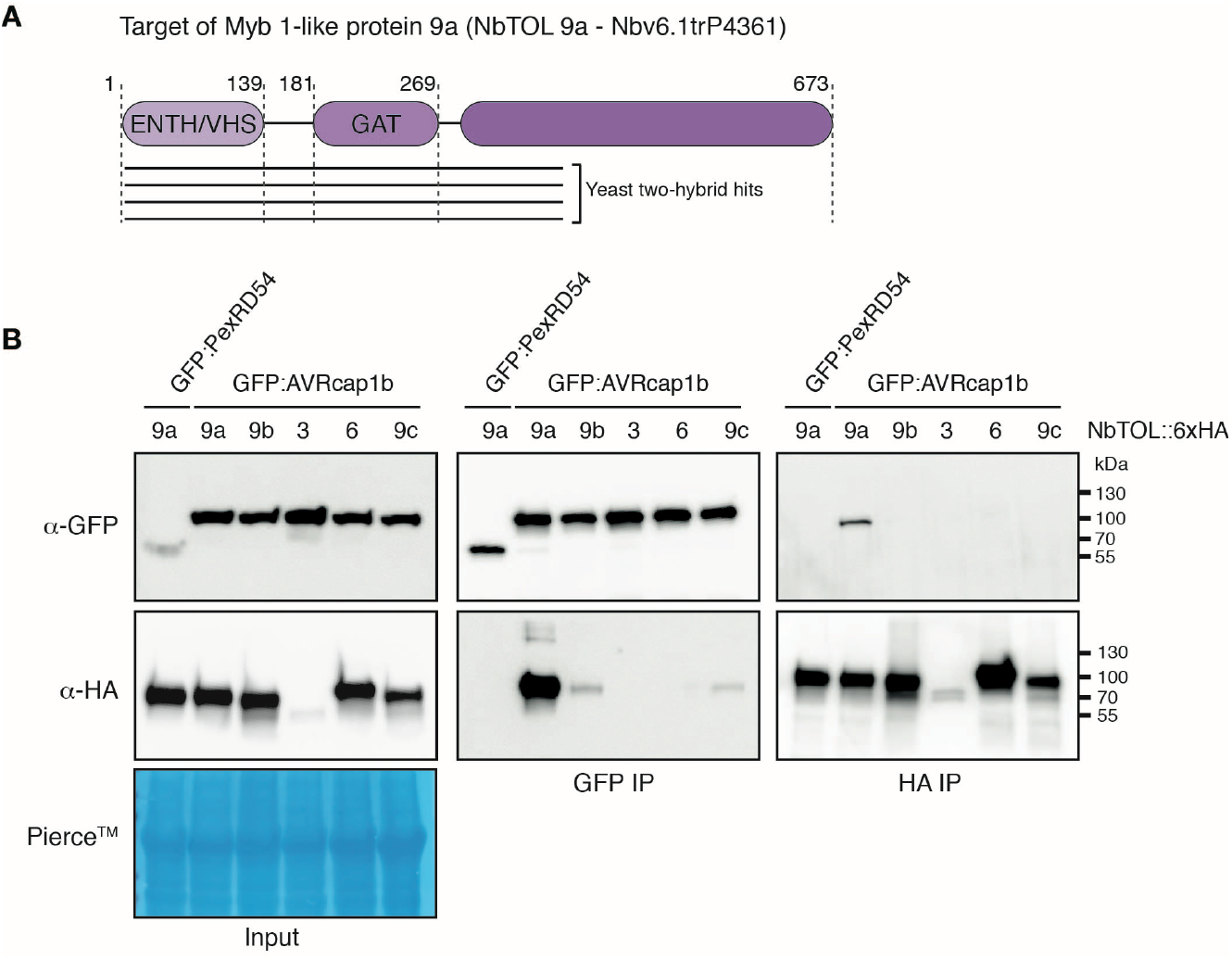
AVRcap1b associates with Target of Myb 1 protein-like 9a (NbTOL9a) *in planta*. (A) Schematic diagram of domain organization of *N. benthamina* Target of Myb 1-like protein (NbTOL) 9a, showing yeast two-hybrid hits from the screen (Table S2). (B) Co-immunoprecipitation experiment between AVRcap1b and five NbTOL family proteins (NbTOL9a, NbTOL9b, NbTOL3, NbTOL6, NbTOL9c). N-terminally GFP-tagged AVRcap1b was transiently co-expressed with all five NbTOL proteins fused to a C-terminal 6xHA tag. N-terminally GFP-tagged PexRD54 was used as a negative control. IP were performed with agarose beads conjugated to either GFP (GFP-IP) or HA (HA-IP) antibodies. Total protein extracts were immunoblotted with appropriate antisera labelled on the left. Approximate molecular weights (kDa) of the proteins are shown on the right. Rubisco loading controls were conducted using Pierce™ staining. This experiment is representative of three independent replicates.

Computational analyses of the *N. benthamiana* genome revealed four TOL paralogs in addition to NbTOL9a, which we termed NbTOL9b (Nbv6.1trP9166), NbTOL3 (Nbv6.1trA40123), NbTOL6 (Nbv6.1trP73492), and NbTOL9c (Nbv6.1trA64113) following previously published nomenclature (Table S5, Figure S12) (53). To validate the association between AVRcap1b and NbTOL proteins, we co-expressed GFP::AVRcap1b with C-terminally 6xHA tagged fusions of the five TOL paralogs, in *N. benthamiana* leaves, and performed anti GFP and anti HA immunoprecipitations. AVRcap1b associated with NbTOL9a, and to a lesser extent with NbTOL9b and NbTOL9c in the GFP pulldown (GFP IP). However, AVRcap1b only associated with NbTOL9a in the reciprocal HA pulldown (HA IP) (Figure 9B). In both experiments, NbTOL9a protein did not associate with the negative control GFP::PexRD54. These results indicate that AVRcap1b associates with members of the NbTOL family, exhibiting a stronger affinity for NbTOL9a. Based on this conclusion, we focused subsequent experiments on NbTOL9a.

### NbTOL9a negatively modulates the cell death triggered by NRC3 but not NRC4

To gain additional insights into the role of NbTOL9a in NRC mediated hypersensitive cell death, we altered NbTOL9a expression in *N. benthamiana*. First, we determined the effect of NbTOL9a overexpression by co-expressing it with the autoimmune NRC3^D480V^ in *N. benthamiana* leaves using agroinfiltration. NbTOL9a overexpression reduced the cell death response triggered by NRC3^D480V^ but did not affect NRC4^D478V^ or the constitutively active MEK2^DD^, a mitogen-activated protein kinase kinase (MAPKK) involved in plant immune signalling, that we included as an additional control (Figure 10, Figure S13).

**Figure 10.**
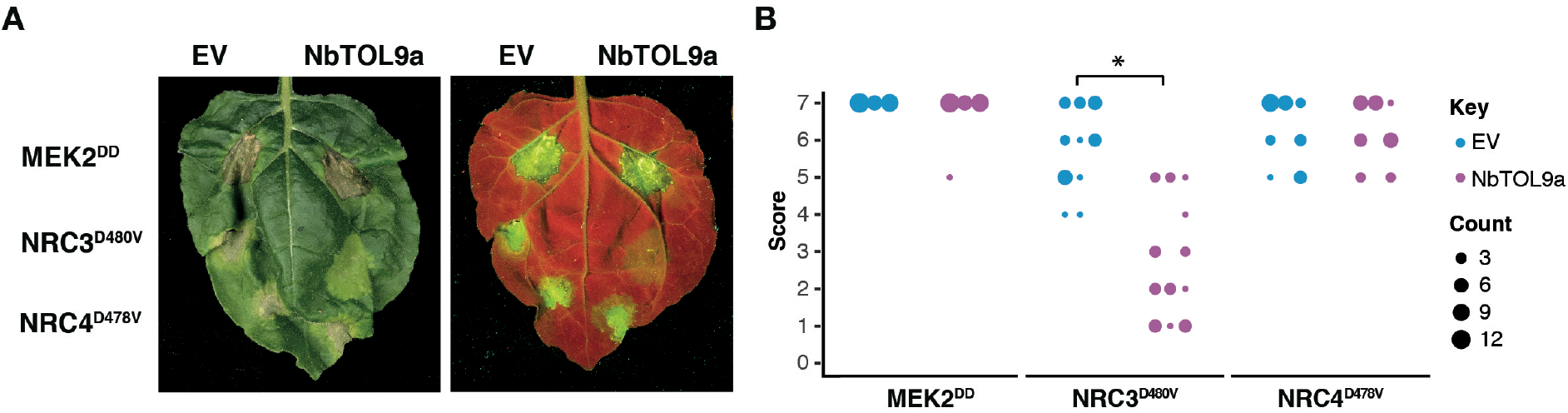
Overexpression of NbTOL9a suppresses autoactive NRC3^D480V^ but not MEK2^DD^ or NRC4^D478V^. (A) Photo of representative *N. benthamiana* leaves showing HR after co-expression of EV and NbTOL9a (labelled above leaf panels) with MEK2^DD^, NRC3^D480V^ and NRC4^D478V^. HR response was scored and photographed 5 days after agroinfiltration (left panel under white light, right panel autofluorescence under UV light). MEK2^DD^ was included as a positive control for cell death. (B) HR results are presented as dot plots, where the size of each dot is proportional to the number of samples with the same score (count). Three biological replicates were completed, indicated by columns for EV, NbTOL9a in each treatment (MEK2^DD^, NRC3^D480V^, NRC4^D478V^). Significant differences between the conditions are indicated with an asterisk (*).

Next, we investigated the effect of silencing *NbTOL9a* on NRC autoimmunity. We generated a hairpin-silencing construct (RNAi::*NbTOL9a*) that mediates silencing of *NbTOL9a* in transient expression assays in *N. benthamiana* leaves (Figure S14). We then co-expressed RNAi::*NbTOL9a* with NRC3^D480V^ using agroinfiltration of *N. benthamiana* leaves. As NRC3^D480V^ gives strong cell death, we used three different concentrations of *A. tumefaciens* expressing NRC3^D480V^ (OD_600_ = 0.1, 0.25 or 0.5) to test the degree to which silencing of *NbTOL9a* affects NRC3-mediated cell death. Silencing of *NbTOL9a* at all tested OD_600_ concentrations enhanced the cell death response triggered by NRC3^D480V^, but did not affect NRC4^D478V^, compared to the RNAi::*GUS* silencing control (Figure 11, Figure S15). Altogether, these two sets of experiments indicate that NbTOL9a modulates NRC3 activity in a manner consistent with a negative regulatory role in NRC3 mediated immunity. Thus, we conclude that AVRcap1b is potentially co-opting the immunomodulator NbTOL9a to suppress NRC mediated immunity.

**Figure 11.**
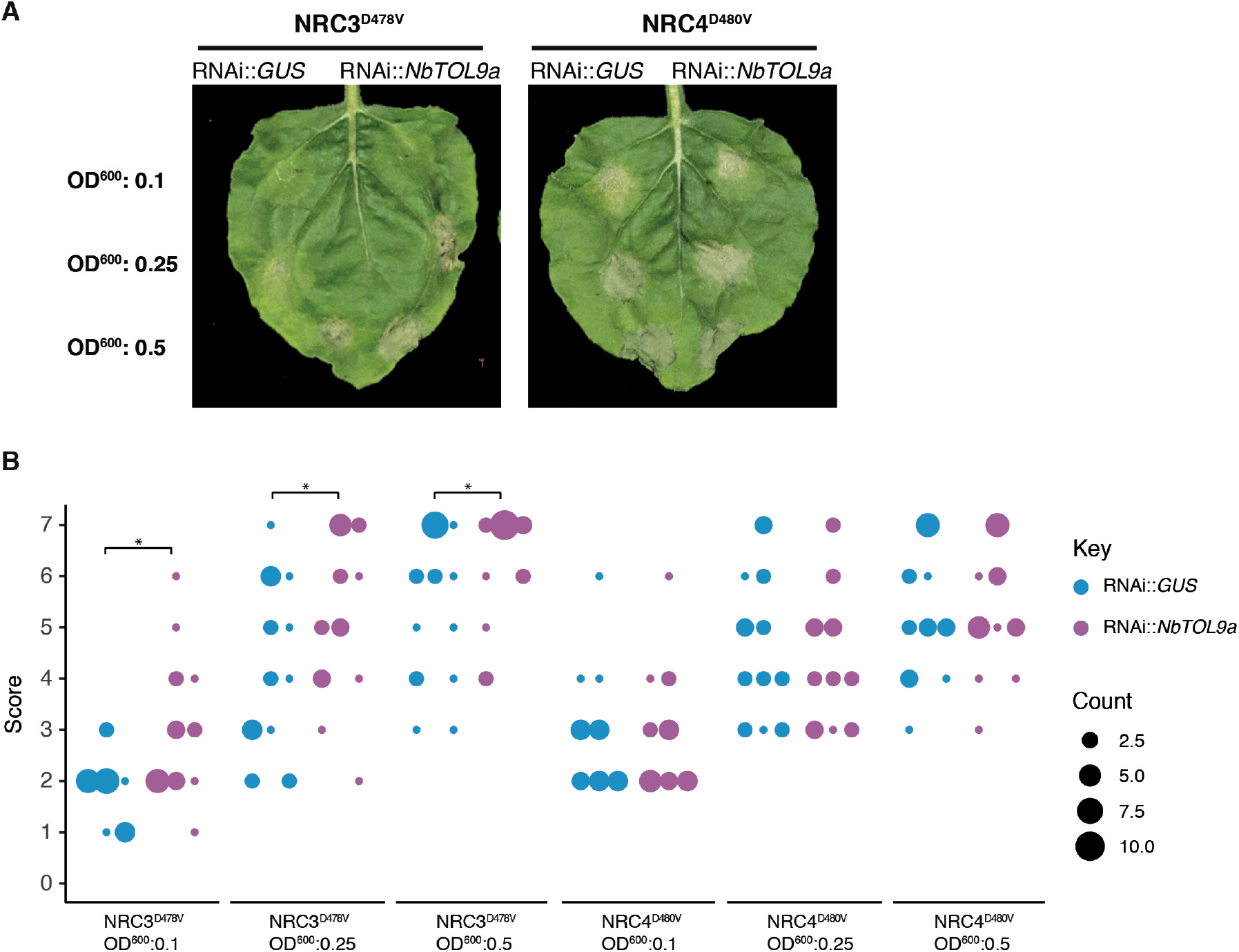
Silencing of NbTOL9a enhances cell death mediated by NRC3^D480V^ but not NRC4^D478V^. (A) Photo of representative *N. benthamiana* leaves showing HR after co-expression of NRC3^D480V^ and NRC4^D478V^, with RNAi::*GUS* (control) and RNAi:*NbTOL9a* (labelled above leaf panels). To improve the robustness of the assay we used increasing concentrations of *A. tumefaciens* expressing NRC3^D480V^ and NRC4^D478V^ (OD_600_ = 0.1, 0.25 or 0.5). HR response was scored and photographed 5 days after agroinfiltration. (B) HR results are presented as dot plots, where the size of each dot is proportional to the number of samples with the same score (count). Three biological replicates were completed, indicated by columns for RNAi::*GUS* and RNAi::*NbTOL9a*, for each treatment combination. Significant differences between the conditions are indicated with an asterisk (*). The details of statistical analysis are presented in (Figure S15).

### AVRcap1b suppression of NRC3 is compromised in the absence of NbTOL9a

To test the hypothesis that NbTOL9a is required for AVRcap1b suppression activity, we co-expressed AVRcap1b with the autoimmune mutants NRC3^D480V^ or NRC4^D478V^ in *N. benthamiana* leaves that are either expressing RNAi::*NbTOL9a* (*NbTOL9a*-silenced) or RNAi::*GUS* (negative control). Consistent with Figure 2, overexpression of AVRcap1b suppressed the cell death triggered by NRC3^D480V^ but not by NRC4^D478V^ (Figure 12). However, silencing of *NbTOL9a* compromised AVRcap1b suppression of NRC3^D480V^ autoimmunity and partially restored the cell death phenotype (Figure 12). These results indicate that AVRcap1b co-opts NbTOL9a to down-regulate NRC3 cell death activity.

**Figure 12.**
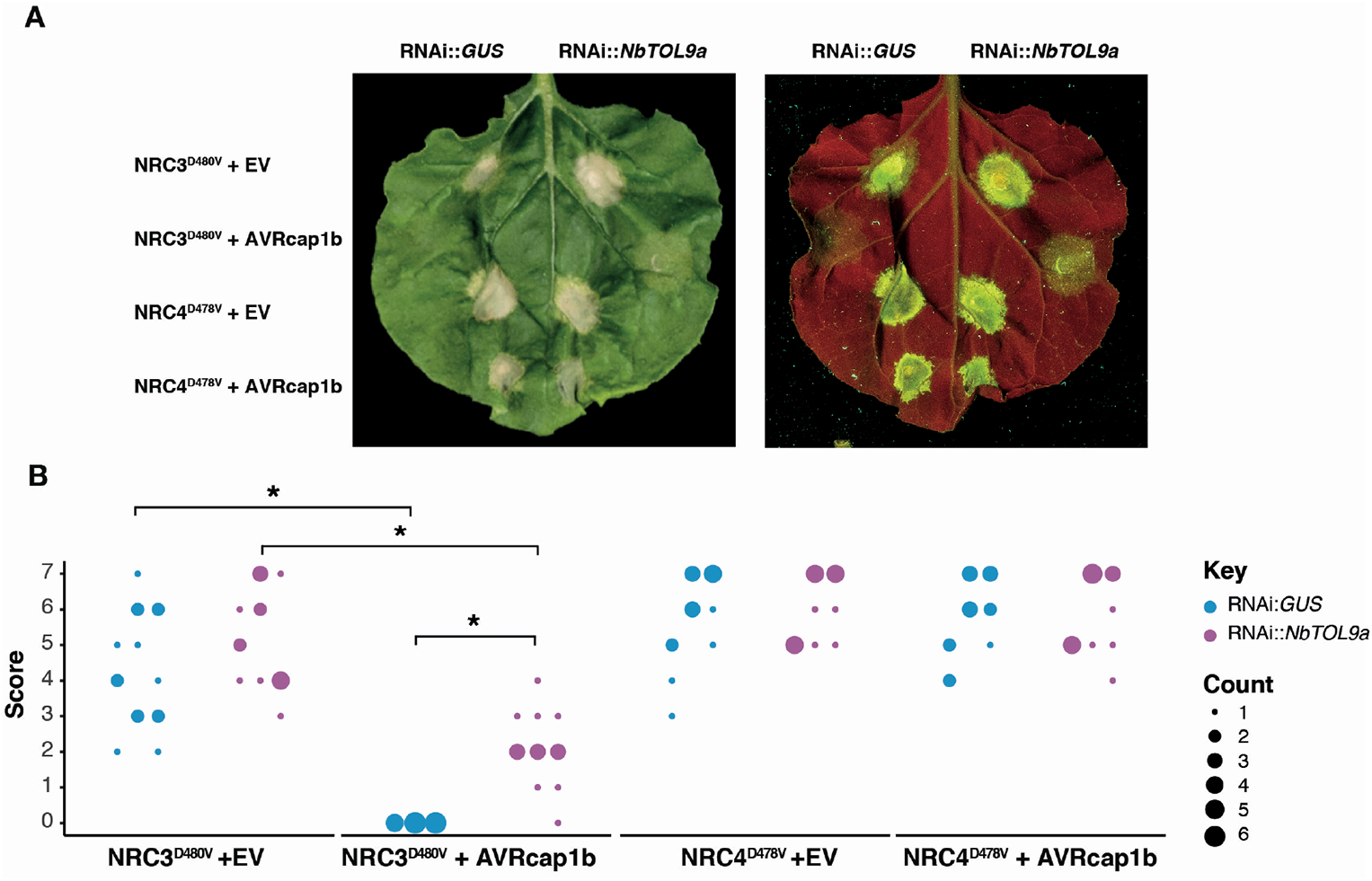
Silencing of NbTOL9a compromises AVRcap1b mediated suppression of NRC3. (A) Photo of representative *N. benthamiana* leaves showing HR after co-expression of RNAi::*GUS* and RNAi::*NbTOL9a* with NRC3^D480V^ + EV, NRC3^D480V^ + AVRcap1b, NRC4^D478V^ + EV and NRC4^D478V^ + AVRcap1b. HR response was scored and photographed 5 days after agroinfiltration (left panel under white light, right panel autofluorescence under UV light). (B) HR results are presented as dot plots, where the size of each dot is proportional to the number of samples with the same score (count). Results are based on three biological replicates. Significant differences between the conditions are indicated with an asterisk (*). The details of statistical analysis are presented in (Figure S16).

## DISCUSSION

The aim of this study was to address the hypothesis that solanaceous parasites have evolved effector proteins that target the NRC network of NLR immune receptors. We confirmed this hypothesis by carrying out an effectoromics screen which yielded five effectors that can compromise the NRC network: SS10, SS15 and SS34 from the cyst nematode *G. rostochiensis* and AVRcap1b and PITG-15278 from the potato late blight pathogen *P. infestans*. These five effectors can suppress the hypersensitive cell death induced in *N. benthamiana* by either Prf or Rpi-blb2, two NRC-dependent sensor NLRs that function as bona fide disease resistance proteins. Interestingly, these effectors appear to function at different points in the NRC network (Figure 13). While SS10, SS34 and PITG-15278 suppress cell death mediated by Rpi-blb2, they do not interfere with an autoimmune mutant of the downstream helper NRC4. SS15 and AVRcap1b, however, can robustly suppress autoimmune mutants of NRC2 and NRC3, indicating that they act at the level of the NRC helpers or their downstream pathways. We found that SS15 directly binds the NB-ARC domain of NRC2 and NRC3, while AVRcap1b associates with NbTOL9a and requires this host protein to fully suppress NRC2 and NRC3. We conclude that cyst nematodes and *P. infestans* convergently evolved effectors that target the same key nodes of the NRC network to suppress host immune signalling. Our paper also highlights the value of using effectors as probes to dissect key regulatory components of immunity and to study the complex interactions between NLR receptors and the networks they form.

**Figure 13.**
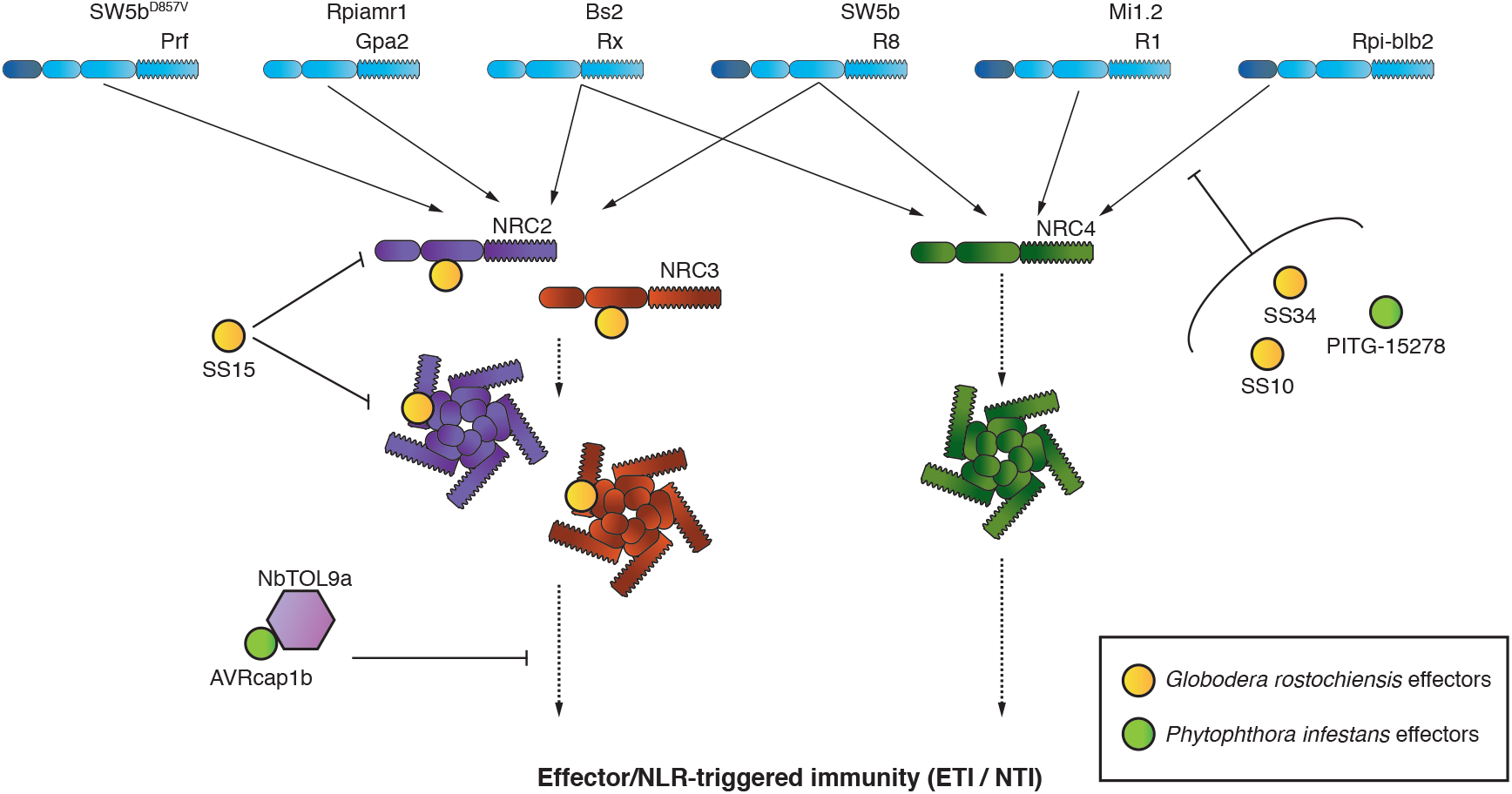
Evolutionary divergent pathogens have evolved to target multiple layers of the Solanaceae NLR network. *P. infestans* and *G. rostochiensis*, two evolutionary unrelated pathogens, have evolved effectors that suppress signaling mediated by the NRC network. Two effectors from the cyst nematode pathogen, *G. rostochiensis*, (SS10, SS34) and one effector from the blight pathogen, *P. infestans* (PITG-15278) suppress the function of the NRC4 – dependent sensor NLR, Rpi-blb2. The cyst nematode effector, SS15, can suppress both inactive and activated NRC2 and NRC3 by binding the NB-ARC domain. The *P. infestans* effector, AVRcap1b, can suppress the function of NRC2 and NRC3 by associating with the ESCRT-related protein NbTOL9a.

Our findings help to explain why plants have evolved NLR receptor networks with complex architectures. We previously postulated that NLR networks, such as the NRC network, help maintain the robustness of the immune system in light of external perturbations (54). A key feature of the NRC network is that NRCs act as central nodes that function downstream of multiple disease resistance proteins (NLR sensors) that form massively expanded phylogenetic clade in the Solanaceae. The NRC nodes have overlapping NLR sensor specificities and display varying degrees of redundancy (25). Given that NRCs are critical for immune signalling of multiple disease resistance proteins, they would be ideal targets for pathogen effectors. Our finding that two unrelated pathogens, an oomycete and a cyst nematode, evolved effectors to suppress NRC signalling through distinct mechanisms, supports this hypothesis. It is tempting to speculate that NRC redundancy has, therefore, emerged as a strategy to enhance the plant’s capacity to evade immune suppression. For example, although both SS15 and AVRcap1b are robust suppressors of NRC2 and NRC3, they are not able to suppress their paralog NRC4. This redundancy would allow the host to mount an effective immune response even in the presence of NRC2 and NRC3 suppressors. Interestingly, the majority of NRC-dependent potato blight R genes, e.g. R1, R8 and Rpi-blb2, signal through NRC4 (25). NRC4 may therefore have evolved as the helper NLR that predominantly functions with late blight R genes because it evades suppression by *P. infestans*. However, Rpi-amr1 was recently shown to require NRC2 and NRC3 for resistance to *P. infestans*, indicating that the interactions between sensor NLRs, helper NRCs and the pathogen effectors are far more complex (55). Nonetheless, this work supports the hypothesis that NLR networks make plant immune systems more resilient through redundant signalling architectures. The extent to which pathogen effectors target NLR networks and the molecular mechanisms by which they do so, however, are still not fully understood.

An emerging paradigm in plant immunity is that NTI and PTI signalling employ common modules and reinforce each other, blurring the division between these two classes of plant immunity (56-59). It is possible that NTI suppressing effectors, such as AVRcap1b and SS15, simultaneously compromise both NTI and PTI by targeting shared immune signalling nodes. Therefore, identification of NTI suppressors may translate into new insights regarding basal resistance and help us further decipher the intricate nature of the plant immune system.

While the precise molecular mechanism that the cyst nematode effector SS15 utilises to suppress NRC2 and NRC3 remains to be determined, our experiments provide some important insights. SS15 binds both inactive (Figure 6B, Figure S9, Figure S10) and constitutively active forms of NRCs (Figure 8), presumably through their NB-ARC domain (Figure 7). There are only a few examples of effectors that directly bind NLRs to suppress their activities. NleA, an effector from human enteropathogenic *Escherichia coli* was shown to suppress the NOD-like receptor (NLR) NLRP3 by directly binding to it, thereby, interfering with de-ubiquitination which is critical for inflammasome activation (60). NleA associates with both ubiquitinated and non-ubiquitinated NLRP3 by interacting with the PYD and LRR domains (61). While the exact mechanism utilised by NleA is currently unknown, the authors theorise that binding of NleA to the PYD and LRR domains may be responsible for inhibition of NLRP3 inflammasome formation by preventing NLRP3 interaction with other downstream signalling partners or blocking access of the de-ubiquitinating enzyme to the polyubiquitinated NLR (61). *P. infestans* IPI-O4 is a plant pathogen effector reported to bind NLRs to suppress host immunity. Chen et al. (62) and Karki et al. (63) showed that the *P. infestans* effector IPI-O4 compromises the hypersensitive response mediated by the NLR disease resistance protein RB (also known as Rpi-blb1) by directly binding to its N-terminal CC domain, possibly to compete with binding by the AVR effector AVRblb1, a homolog of IPI-O4. Therefore, IPI-O4 acts directly on the sensor NLR and is more reminiscent of the three Rpi-blb2 suppressors we report here than to the NRC suppressors AVRcap1b and SS15 (Figure 13).

The recent elucidation of the ZAR1, RPP1 and ROQ1 structures have revealed that structural remodelling of the NB-ARC domain, a region involved in NLR activation, is essential for resistosome formation (28-31). The mechanism utilised by SS15 to suppress NRC mediated immunity may involve tampering with NB-ARC structural rearrangements. In mammalian systems, for example, a compound known as MCC950 directly binds both the inactive and activated forms of NLRP3 to inhibit its function (64, 65). MCC950 binds the central NACHT domain, the mammalian equivalent of the NB-ARC domain of plant NLRs, to interfere with ATP hydrolysis and prevent conformational changes that are critical for NLRP3 activation and subsequent inflammasome assembly. This ultimately drives NLRP3 towards a closed and inactive conformation (64, 65). Based on our findings, we propose that SS15 could be acting as an NLR inhibitor that directly perturbs NRC activities by binding the NB-ARC domain and forcing NRC2 and NRC3 into an inactivated state, possibly through mechanisms analogous to MCC950. Further studies investigating the extent to which SS15 perturbs structural remodelling of the NB-ARC domain of NRCs will provide mechanistic insights into how this effector is able to suppress NRC2 and NRC3.

Unlike SS15, AVRcap1b from the potato late blight pathogen *P. infestans*, indirectly suppresses the function of autoimmune NRC2 and NRC3 and therefore likely targets host proteins downstream of these cell death executor NLRs. We identified NbTOL9a, a member of the TOL protein family, as a host target of AVRcap1b and showed that this host protein acts as a negative modulator of NRC3-mediated hypersensitive cell death. TOL proteins are key components of the ESCRT machinery and have well characterized roles in intracellular protein trafficking (52, 66, 67). They act as ubiquitin receptors in the early steps of the ESCRT trafficking pathway, by interacting with ubiquitinated cargo via their ENTH/VHS and GAT domains (51-53). Conlan et al. (68) identified a TOL family member that is proximal to the *P. syringae* effector AvrPto, when this effector protein was transiently expressed in *N. benthamiana* leaf tissue. In addition, silencing of this TOL protein resulted in decreased growth of *P. syringae* pv. *tabaci* on *N. benthamiana*, which is in line with our observation that TOL proteins can act as negative regulators of plant immune signalling. The fact that AvrPto is proximal to TOLs further links the NRC network to TOL proteins, since AvrPto is recognised by the NRC2 and NRC3-dependent sensor NLR Prf. Together these findings strengthen our hypothesis that TOLs can act as negative regulators of this complex immune signalling network. The precise mechanism TOLs utilize to modulate plant immunity, however, is still unknown. In mammalian systems, the ESCRT pathway is involved in negatively regulating several forms of programmed cell death, including necroptosis and pyroptosis, by repairing damaged sections of the plasma membrane (69-71). It is possible that AVRcap1b is co-opting NbTOL9a to hijack a similar immunomodulatory trafficking pathway in the plant cell to counteract NRC-mediated HR cell death and suppress immunity. This model would be consistent with the observation that vesicle trafficking is massively reprogrammed by *P. infestans* effectors during infection (49, 72-74).

Interestingly, even though AVRcap1b is a robust NTI suppressor, it can activate immunity on accessions of *Solanum capsicibaccatum* carrying the NLR disease resistance gene Rpi-cap1– hence the moniker AVR (33, 75). The fact that AVRcap1b can act as both a trigger and suppressor of NLR immunity further highlights the complex coevolutionary dynamics that exist between effectors and NLRs and the need for studies that take into account these intricate epistatic interactions. Understanding the interplay between effectors, TOLs and NRCs will significantly advance our knowledge of the regulatory mechanisms that govern plant immunity and determine the outcome of multipartite host-pathogen interactions mediated by a complex immune receptor network.

The field of NLR biology has seen significant progress over the past few years, and yet the genetic components and immune signalling pathways downstream of NLR activation remain obscure (76). Moreover, the precise molecular mechanisms that underpin the activation of paired and networked NLRs and subsequent cell death response are still unknown. Here, we have gained insights into the molecular strategies that plant pathogens utilise to counteract host immune function. We identified two effectors, SS15 and AVRcap1b, that have the unique potential to uncover valuable mechanistic details regarding the activation of MADA type CC-NLR resistosomes and their downstream signalling elements. The fact that these distantly related pathogens, an oomycete and a cyst-nematode, have independently evolved effectors to counteract NRC2 and NRC3 further highlights the critical role NRCs play in mediating immunity to solanaceous parasites. Recently, effectors from the oomycete pathogen *Phytophthora capsici* and the aphid pest *M. persicae* were shown to converge on host E3 SUMO ligase SIZ1 to suppress plant immunity (77). Along with our work, this indicates that microbial pathogens, herbivorous insects and parasitic nematodes may share more common virulence mechanisms than anticipated. Additionally, further work on these immunosuppressors will hopefully advance our understanding of the functional principles and evolutionary dynamics that underpin plant immune receptor networks. This knowledge can then be leveraged to guide new approaches for breeding disease resistance to maximize crop protection, for example by engineering NLRs that evade pathogen suppression.

## MATERIALS AND METHODS

### Plant growth conditions

Wild type and Rx transgenic (78) *Nicotiana benthamiana* lines were grown in a controlled growth chamber with temperature 22-25°C, humidity 45-65% and 16/8-h light/dark cycle.

### Gene synthesis and cloning

#### Effector library

All sequence information for pathogen effectors used in suppression assays can be found in Table S1.

*Phytophthora infestans* effectors, without signal peptide (34, 35) were synthesized by GENEWIZ (South Plainfield, NJ, USA) into pUC57-Amp. Effector sequences were amplified from the pUC57-Amp vector by Phusion High-Fidelity DNA Polymerase (Thermo Fisher), and the purified amplicon was used directly for Golden Gate assembly into the Level 0 Universal Acceptor pUAP1 (The Sainsbury Laboratory (TSL) SynBio, addgene no. 63674). Primers used for PCR amplification are listed in Table S6. Expression constructs were generated by Golden Gate assembly of the Level 0 module into binary vector pICSL86977OD [with Cauliflower Mosaic Virus (CaMV) 35S promoter and octopine synthase gene) terminator] (addgene no. 86180). PITG-05910 and PITG-22926 were synthesized by GENEWIZ (South Plainfield, NJ, USA) into pUC57-kan as Golden Gate L0 modules, then subcloned into binary vector pICH47742 [addgene no. 48001], together with pICH51266 [35S promoter+Ω promoter, addgene no. 50267], and pICH41432 [octopine synthase terminator, addgene no. 50343]. All constructs were verified by sequencing.

*Globodera rostochiensis* effectors; SS4, SS8, SS9, SS15, SS16 and SS19 cloned into pBINPLUS expression vector with N-terminal 4xHA tag were provided by Aska Goverse (Laboratory of Nematology, Wageningen University, Netherlands) (37, 79). *G. pallida* effector sequences 12N3, 33H17 (80) were synthesized by GENEWIZ (South Plainfield, NJ, USA) as Golden Gate Level 0 modules into pICH41155, Gpa-SS37 (81) was synthesized by GENEWIZ (South Plainfield, NJ, USA) as a Golden Gate Level 0 module into pUC57-kan. The remaining *G. rostochiensis* effector sequences were extracted from the https://parasite.wormbase.org website (38) and synthesized by GENEWIZ (South Plainfield, NJ, USA) as Golden Gate Level 0 modules into pUC57-kan. These effectors were subcloned into binary vector pICH47732 [addgene no. 50434], together with pICH51266 [35S promoter+Ω promoter, addgene no. 50267], and pICH41432 [octopine synthase terminator, addgene no. 50343] for cell death suppression assays.

The green peach aphid (*Myzus persicae*) and pea aphid (*Acrythosiphon pisum*) effectors cloned into pCB302-3 in *Agrobacterium tumefaciens* strain GV3101::pM90, were provided by Saskia Hogenhout (John Innes Centre, Norwich, United Kingdom). The tomato bacterial speck pathogen (*Pseudomonas syringae*) effectors cloned into pEarly Gate 100 in *A. tumefaciens* strain C58C1 were kindly provided by Wenbo Ma (previously University California, Riverside, United States of America, currently The Sainsbury Laboratory, Norwich, United Kingdom).

#### Synthesis of NRC constructs

Full length *N. benthamiana* NRC2a^syn^ (NCBI accession number KT936525) (25), NRC3^WT^ and NRC4^WT^ were synthesized by GENEWIZ (South Plainfield, NJ, USA) into pICH41155, as Golden Gate L0 modules (Table S7). These sequences were manually domesticated to remove BpiI and BsaI sites.

#### Site-directed mutagenesis of NRC2, NRC3 and SW5b

Autoactive mutants of NRC2, NRC3 and SW5b were generated by introducing a histidine (H) to arginine (R) mutation in the MHD motif of NRC2 and an aspartic acid (D) to valine (V) substitution in the MHD motif of NRC3 and SW5b independently. Primers listed in Table S8 were used for introducing mutations by inverse PCR with Phusion High-Fidelity DNA Polymerase (Thermo Fisher). pICH41155::NRC2, PCR8::NRC3^WT^ (27), and pICH441155::SW5b (25) were used as templates for site-directed mutagenesis of NRC2, NRC3, and SW5b, respectively. The mutated variants were verified by sequencing and then subcloned into pICSL86977OD [addgene no. 86180] (for NRC2^H480R^ and SW5b^D847V^) and pICH86988 [addgene no. 48076] (NRC3^D480V^). Autoactive NRC4 mutants used in this study, were described previously (26). Verified plasmids were then transformed into GV3101:pM90 for cell death assay screens.

To determine whether an intact p-loop is essential for SS15 *in planta* association to NRC2 and NRC3, a lysine (K) to arginine (R) mutation was introduced independently into the p-loops of both proteins by site-directed mutagenesis using Phusion High-Fidelity DNA Polymerase (Thermo Fisher). pCR8::NRC2^WT^-ns and PCR8::NRC3^WT^-ns (27) were used as a templates. Primers used to for introducing mutations are listed in Table S8. The mutated NRC variants were verified by sequencing and subcloned into pICH86988 [addgene no. 48076] together with pICSL50010 [C-terminal 4xMyc tag, addgene no. 50310]. Both constructs were verified by sequencing and transformed into *A. tumefaciens* GV3101::pM90.

#### Cloning into *E. coli* expression vectors for protein purification

SS15: DNA encoding SS15 residues Ser-25 to Ile-246 (lacking signal peptide) was amplified from pBINPLUS::4HA::SS15 plasmid (using primers shown in Table S15) and cloned into the pOPINS3C vector resulting in an N-terminal 6xHis-SUMO tag with SS15, linked by a 3C cleavage site (pOPINS3C:SS15) (82).

NB-ARC domains of NRC2, NRC3 and NRC4: For NRC2, two NB-ARC domain constructs with alternative domain boundaries were generated. DNA encoding NRC2 NB-ARC domain residues Val-148 to Tyr-508 (NRC2^148-508^) or Val-148 to Asn-496 (NRC2^148-496^) was amplified from pICH41155::NRC2a^syn^ (using primers shown in Table S15) and cloned into the vector pOPINS3C resulting in an N-terminal 6xHis-SUMO tag with NRC2 NB-ARC, linked by a 3C cleavage site (pOPINS3C:NRC2^148-508^ and pOPINS3C:NRC2^148-496^, respectively). Similarly, DNA encoding NRC3 NB-ARC residues Val-152 to Ser-507 (NRC3^NB-ARC^) was amplified from pICH41155::NRC3^WT^ (using primers shown in Table S15) and DNA encoding NRC4 NB-ARC domain Ala-150 to Lys-491 (NRC4^NB-ARC^) was amplified from pICH41155::NRC4^WT^ (using primers shown in Table S15) and cloned into the vector pOPINS3C as described above (pOPINS3C:NRC3^NB-ARC^ and pOPINS3C:NRC4^NB-ARC^, respectively).

#### Generating C-terminally 4xMyc tagged NRC2 and NRC3 autoactive mutants

To generate NRC2^H480R^::4xMyc and NRC3^D480V^::4xMyc, the stop codon was removed from pICSL86977OD::NRC2^H480R^ and pICH86988::NRC3^D480V^ by Phusion High-Fidelity DNA polymerase (Thermo Fischer), using primers in Table S8. The purified amplicon was used directly for Golden Gate assembly into binary vector pICH86988 [addgene no. 48076], together with pICSL50010 [C-terminal 4xMyc tag, addgene no. 50310]. Both constructs were verified by DNA sequencing and then transformed into *A. bacterium* strain GV3101::pM90.

### Generating N-terminally GFP tagged and C-terminally 6xHA tagged AVRcap1b

AVRcap1b constructs used in coIP experiments were generated by Golden Gate assembly. To generate GFP::AVRcap1b, level 0 pUAP1::AVRcap1b was assembled into binary vector pICH86966 [addgene no. 48075] driven by pICSL13008 [CaMV 35S promoter+5’ untranslated leader tobacco mosaic virus, TSL synbio], pICSL30006 [GFP, addgene no. 50303], pICH41432 [octopine synthase terminator, addgene no. 50343]. To generate AVRcap1b::6xHA, the stop codon was removed from pUAP1::AVRcap1b by Phusion High-Fidelity DNA Polymerase (Thermo Fisher), using primers listed in Table S6. The purified amplicon was used directly for Golden Gate assembly into binary vector pICH47742 [addgene no. 48001] driven by pICH85281 [mannopine synthase promoter+Ω (MasΩpro), Addgene no. 50272], pICSL50009 [6xHA, addgene no. 50309], pICSL60008 [Arabidopsis heat shock protein terminator (HSPter), TSL SynBio]. All constructs were verified by DNA sequencing and then transformed into *A. bacterium* strain GV3101::pM90.

#### Cloning of NbTOL paralogs

We used reciprocal BLAST searches of Nbv6.1trP4561, which was identified in both IP-MS and yeast two-hybrid assays, and mined the V6.1 transcriptome (https://benthgenome.qut.edu.au/) for additional TOL sequences. We cross validated the extracted TOLs with the genome-based predicted proteomes (83) and removed likely duplicates representing chimeric transcripts due to assembly of short read sequences. We identified a total of five TOL paralogs, which we termed NbTOL9a (Nbv6.1trP4361), NbTOL9b (Nbv6.1trP9166), NbTOL3 (Nbv6.1trA40123), NbTOL6 (Nbv6.1trP73492), and NbTOL9c (Nbv6.1trA64113) (Table S5). NbTOL paralogs were synthesized using GENEWIZ (South Plainfield, NJ, USA) into pUC57-Kan as L0 modules. Individual TOL paralogs were subjected to PCR by Phusion High-Fidelity DNA Polymerase (Thermo Fisher) to remove the stop codon (Primer information listed in Table S12). The purified amplicons were used directly for Golden Gate assembly into binary vector pICH47732 [addgene no. 50434], together with pICH51266 [35S promoter+Ω promoter, addgene no. 50267], pICSL50009 [6xHA, addgene no. 50309], and pICH41432 [octopine synthase terminator, addgene no. 50343]. All constructs were verified by DNA sequencing and transformed into *A. tumefaciens* GV3101::pM90.

### Cell death and suppression assays by agroinfiltration

Information of constructs used for cell death assays are summarized in Table S9. Transient expression of NLR immune receptors and cognate effectors, and autoactive immune receptors with empty vector (EV) or effector constructs were performed according methods previously described (13). Briefly, four-to-five-week-old *N. benthamiana* plants were infiltrated with *A. tumefaciens* GV3101::pM90 strains carrying the expression vectors of different indicated proteins within the text. *A. tumefaciens* suspensions were adjusted in infiltration buffer (10 mM MES, 10 mM MgCl_2_, and 150 μM acetosyringone, pH5.6) to final OD_600_ indicated in Table S9. OD_600_ for all effectors were adjusted to 0.2. For NbTOL9a overexpression assays, NbTOL9a::6xHA construct was co-infiltrated at a final OD_600_ of 0.2. The cell death, hypersensitive response (HR), phenotype was scored 5 – 7 days after agroinfiltration, unless otherwise stated, using a previously described scale (84), modified from 0 (no necrosis observed) – 7 (confluent necrosis).

### Virus induced gene silencing (VIGS) of NRC homologs

VIGS was performed in *N. benthamiana* as previously described (85). Suspensions of *A. tumefaciens* strain GV3101::pM90 harbouring TRV RNA1 (pYL155) and TRV RNA2 (pYL279), with corresponding fragments from *NRC2, NRC3* and *NRC4*, were mixed in a 2:1 ratio in infiltration buffer (10 mM 2-[N-morpholine]-ethanesulfonic acid [MES]; 10 mM MgCl_2_; and 150 μM acetosyringone, pH 5.6) to a final OD_600_ of 0.3. Two-week-old *N. benthamiana* plants were infiltrated with *A. tumefaciens* for VIGS assays, upper leaves were used two to three weeks later for further agroinfiltrations. The *NRC2/3* double silencing, *NRC4* and *NRC2/3/4* triple silencing constructs were described previously (25, 27).

### PVX infection assays (agroinfection)

Plants were used three weeks after TRV infection. Plants were infected with *Potato virus X* (PVX, pGR106) using the toothpick inoculation method, expressed via *A. tumefaciens* (25, 46), and examined for the spread of trailing necrotic lesions from the inoculated spots (43). pGR106::PVX::GFP construct generation was described previously (25). One day before PVX toothpick inoculation, EV and pICH86977::AVRcap1b or pBINPLUS::4HA::SS15 constructs were expressed by agroinfiltration into leaves of *Rx* plants that were subjected to *NRC2/3* double, *NRC4* and *NRC2/3/4* triple silencing. The infiltrated area was then circled with a marker pen. *NRC* homologs were silenced by VIGS as described above in *Rx* transgenic *N. benthamiana*. Toothpicks were dipped into culture of *A. tumefaciens* harboring the PVX-GFP vector and then used to pierce small holes in the leaves of *N. benthamiana*. Photos were taken at three weeks after PVX inoculation, and the size of the lesions were measured in Fiji (formerly ImageJ). Scatterplot of the lesion size was generated in R using the ggplot2 package, as described previously (86). A core borer (0.8 cm^2^) was used to collect leaf discs from the inoculation sites three weeks after PVX inoculation for immunoblot with anti-GFP (B-2, sc9996 HRP, Santa Cruz Biotechnology). GFP accumulation was visualised with Pierce™ ECL Western (32106, Thermo Scientific) and where necessary up to 50% SuperSignal™ West Femto Maximum Sensitivity Substrate (34095, Thermo Scientific).

### Phylogenetic analyses of *Nicotiana benthamiana* and *Arabidopsis thaliana* TOL proteins

Amino acid sequences of the NbTOL paralogs identified in *N. benthamiana* and previously published *Arabidopsis thaliana* AtTOL proteins (53, 87) were aligned using Clustal Omega (88). The alignment was then manually edited in MEGAX (89). The gaps in the alignment were manually removed and only the ENTH and GAT domains were used to generate the phylogenetic tree. A maximum-likelihood tree of the *N. benthamiana* and *A. thaliana* TOLs was generated in MEGAX using the JTT model and with bootstrap values based on 1000 iterations (Figure S12). The resulting tree was then visualized using iTOL (90). The alignment used to make the tree is provided as Supplementary File 1.

### Hairpin RNA-mediated gene silencing

The silencing fragment was amplified out of *N. benthamiana* cDNA by Phusion High-Fidelity DNA Polymerase (Thermo Fisher) using primers listed in Table S10. The specificity of the silencing fragment was analysed using the *N. benthamiana* genome sequence and associated gene silencing target prediction tool (SGN VIGS tool: https://vigs.solgenomics.net). The purified amplicon was cloned into pRNAi-GG vector according to Yan et al (91). Construct was verified by DNA sequencing and then transformed into *A. tumefaciens* strain GV3101::pM90. Silencing evading NbTOL9a (NbTOL9a^syn^) was synthesized by GENEWIZ (South Plainfield, NJ, USA) in pUC57-kan and generated according to Wu et al (25). Leaves were co-infiltrated with either pRNAi-GG::*NbTOL9a* or pRNAi-GG::*GUS*, at a final OD_600_ of 0.5, together with different proteins indicated in the text with final OD_600_ indicated in Table S10. The HR cell death on the leaves was scored at 5-7 days as described above.

### Protein-Protein interaction studies

#### Yeast two-hybrid screens

Unbiased Yeast two-hybrid screens were performed by Hybrigenics Services (http://www.hybrigenics.com, Paris, France). For AVRcap1b (residues Ala62 – Pro678, lacking the signal peptide) was cloned into both the pB27 and pB66 bait plasmids, as a C-terminal fusions to LexA (LexA-AVRcap1b) and Gal4 (Gal4-AVRcap1b), respectively. The two screens were performed against a randomly-primed *N. benthamiana* mixed tissue cDNA library. For LexA-AVRcap1b a total of 52.8 million interactions were screened (∼5-fold library coverage) and nine positive clones were fully analysed. For Gal4-AVRcap1b a total of 87.6 million interactions were screened (∼8-fold library coverage) and 26 positive clones were fully analysed. These clones correspond to 14 different annotated proteins, one of which was unknown (Table S2). For SS15 (residues Ser25 – stop 246, lacking the signal peptide) was cloned into pB27 as a C-terminal LexA bait (LexA-SS15) and screened against a randomly-primed *N. benthamiana* mixed tissue cDNA library. A total of 61.4 million interactions were screened (∼6-fold library coverage) and 202 positive clones were fully processed, corresponding to 10 different annotated proteins, three of which were unknown (Table S3). Interactions were categorised based on their Predicted Biological Score, a measure to assess the interaction reliability, which is calculated based on a statistical model of the competition for bait-binding between fragments (92, 93).

To narrow down the region mediating protein–protein interactions between SS15 and NRCs, we generated NRC2, NRC3 and NRC4 CC and NB-ARC domain truncations and ran a yeast two-hybrid experiment using the Matchmaker Gold system (Takara Bio USA). Plasmid DNA encoding AVRcap1b (negative control) and SS15 in pGBKT7 (bait), generated in this study, (using primers shown in Table S11), were co-transformed into chemically competent Y2HGold cells (Takara Bio, USA) with individual NRC truncates in pGADT7 (prey) (using primers shown in Table S11), as described previously (94, 95), in the AH109 yeast strain. Single colonies grown on selection plates were inoculated in 5 ml of SD^-Leu-Trp^ overnight at 28 °C (ST0047, Takara Bio, USA). Saturated culture was then used to make serial dilutions of OD_600_ 1, 10^−1^ and 10^−2^, respectively. 3 μl of each dilution was then spotted on a SD^-Leu-Trp^ plate (ST0048, Takara Bio, USA) as a growth control, and on a SD^-Leu-Trp-Ade-His^ plate (ST0054, Takara Bio, USA) containing X-α-gal and supplemented with 0.2% Adenine. Plates were incubated for 3 – 6 days at 28 °C and then imaged. Each experiment was repeated a minimum of three times, with similar results. The commercial yeast constructs were used as positive (pGBKT7-53/pGADT7-T) and negative (pGBKT7-Lam/pGADT7-T) controls (clontech).

#### *In planta* CoIPs

Protein samples were extracted from two *N. benthamiana* leaves three days post agroinfiltration and homogenised in GTEN extraction buffer [10% glycerol, 25 mM Tris-HCl, pH 7.5, 1 mM EDTA, 150 mM NaCl, 2% (w/v) polyvinylpolypyrrolidone, 10 mM dithiothreitol, 1x protease inhibitor cocktail (SIGMA), 0.2% IGEPAL (SIGMA)] (96). The final concentration OD_600_ used for each protein is indicated in Table S13. After centrifugation at 5000 x*g* for 20 minutes, the supernatant was passed through a Minisart 0.45μM filter (sartorius stedium) and used for SDS-PAGE. For coIP, 1.4ml of filtered total protein extract was mixed with 30μl of GFP-Trap®-A agarose beads (chromatek, Munich, Germany), anti-c-myc agarose beads (A7470, SIGMA) or anti-HA affinity matrix beads (Roche) and incubated end-over-end for 1 hr at 4°C. Beads were washed five times with immunoprecipitation wash buffer [GTEN extraction buffer with 0.3% (v/v) IGEPAL (SIGMA)] and re-suspended in 70μl SDS loading dye. Proteins were eluted from beads by heating at 10 minutes at 70°C (for GFP) or 95°C (for myc or HA). Immunoprecipitated samples were separated by SDS-PAGE and transferred onto a polyvinylidene difluoride membrane using Trans-Blot turbo Transfer system (Bio-Rad, Munich), according to the manufacturer’s instructions. Blots were pre-blocked with 5% skim milk powder in Tris-buffered saline plus Tween 20 (TBS-T) overnight in 4°C, or for a minimum of 1hr at room temperature. Epitope tags were detected with HA-probe (F-7) horse radish peroxidase (HRP)-conjugated antibody (Santa Cruz Biotech), c-Myc (9E10) HRP (Santa Cruz Biotech), or anti-GFP (B-2) HRP (Santa Cruz Biotech) antibody in a 1:5,000 dilution in 5% skim milk powder in TBS-T. Proteins were visualised with Pierce™ ECL Western (32106, Thermo Scientific) and where necessary up to 50% SuperSignal™ West Femto Maximum Sensitivity Substrate (34095, Thermo Scientific). Membrane imaging was carried out with an ImageQuant LAS 4000 luminescent imager (GE Healthcare Life Sciences, Piscataway, NJ). Rubisco loading control was stained using Pierce™ (24580, Thermo Scientific) or Instant Blue (Expedeon, Cambridge).

#### Protein purification from *E. coli* and *in vitro* protein-protein interaction studies

Recombinant SS15 protein (lacking signal peptide) was produced using *E. coli* SHuffle cells (97) transformed with pOPINS3C:SS15 (see gene cloning and synthesis section above). Cell culture was grown in autoinduction media (98) at 30°C to an A_600_ 0.6–0.8 followed by overnight incubation at 18°C and harvested by centrifugation. Pelleted cells were resuspended in 50 mM Tris HCl pH 8, 500 mM NaCl, 50 mM Glycine, 5% (vol/vol) glycerol and 20 mM imidazole (buffer A) supplemented with cOmplete EDTA-free protease inhibitor tablets (Roche) and lysed by sonication. The clarified cell lysate was applied to a Ni^2+^-NTA column connected to an AKTA pure system. 6xHis+SUMO-SS15 was step-eluted with elution buffer (buffer A containing 500 mM imidazole) and directly injected onto a Superdex 200 26/600 gel filtration column pre-equilibrated in buffer B (20 mM HEPES pH 7.5, 150 mM NaCl and 1mM TCEP). The fractions containing 6xHis+SUMO-SS15 were pooled and concentrated to 2–3 mg/ml. The 6xHis+SUMO tag was cleaved by addition of 3C protease (10 µg/mg fusion protein) and incubation overnight at 4°C. Cleaved SS15 was further purified using a Ni^2+^-NTA column (collecting the eluate) followed by gel filtration as above. The concentration of protein was judged by absorbance at 280 nm (using a calculated molar extinction coefficient of SS15, 35920 M^−1^cm^−1^).

Recombinant NRC3^NB-ARC^ was produced using *E. coli* Lemo21 (DE3) cells transformed with pOPINS3C:NRC3^NB-ARC^ (see gene cloning and synthesis section for details). Cell culture was grown in autoinduction media at 30°C to an A_600_ 0.6–0.8 followed by overnight incubation at 18°C and harvested by centrifugation. Pelleted cells were resuspended in buffer A supplemented with EDTA free protease inhibitor tablets and lysed by sonication. The clarified cell lysate was applied to a Ni^2+^-NTA column connected to an AKTA pure system. 6xHis+SUMO-NRC3^NB-ARC^ was step-eluted with elution buffer (buffer A containing 500 mM imidazole) and directly injected onto a Superdex 200 26/600 gel filtration column pre-equilibrated in buffer B (20 mM HEPES pH 7.5, 150 mM NaCl and 1mM TCEP). The fractions containing 6xHis+SUMO-NRC3^NB-ARC^ were pooled and concentrated to 1 mg/ml. The 6xHis+SUMO tag was cleaved by addition of 3C protease (10 µg/mg fusion protein) and incubation overnight at 4°C. Cleaved NRC3^NB-ARC^ was further purified using a Ni^2+^-NTA column (collecting the eluate) followed by gel filtration as above. The concentration of protein was judged by absorbance at 280 nm (using a calculated molar extinction coefficient of NRC3^NB-ARC^, 48020 M^−1^cm^−1^).

To purify SS15 in complex with NRC3^NB-ARC^, *E. coli* cultures were grown individually for SS15 and NRC3^NB-ARC^ and harvested by centrifugation as explained above. Pelleted cells expressing SS15 and NRC3^NB-ARC^ were resuspended separately in buffer A supplemented with EDTA free protease inhibitor tablets. After resuspension, SS15 and NRC3^NB-ARC^ expressing cells were mixed and lysed together by sonication. The clarified cell lysate (containing 6xHis+SUMO-NRC3^NB-ARC^ and 6xHis-SUMO-SS15) was applied to a Ni^2+^-NTA column connected to an AKTA pure system. 6xHis+SUMO-tagged protein (SS15 and NRC3^NB-ARC^) was step-eluted with elution buffer (buffer A containing 500 mM imidazole) and directly injected onto a Superdex 200 26/600 gel filtration column pre-equilibrated in buffer B (20 mM HEPES pH 7.5, 150 mM NaCl and 1mM TCEP). The fractions containing 6xHis+SUMO-SS15 and 6xHis+SUMO-NRC3^NB-ARC^ were pooled and concentrated to 1-2 mg/ml. The 6xHis+SUMO tag was cleaved by addition of 3C protease (10 µg/mg fusion protein) and incubation overnight at 4°C. Cleaved NRC3^NB-ARC^ and SS15 complex was further purified using a Ni^2+^-NTA column (collecting the eluate) followed by gel filtration as above. The fractions containing complex of NRC3^NB-ARC^ and SS15 were pooled together and concentrated. The concentration of complex was judged by absorbance at 280 nm (using a calculated molar extinction coefficient of SS15 and NRC3^NB-ARC^, 83940 M^−1^cm^−1^).

### IP-MS

Total proteins were extracted from *N. benthamiana* leaves three days after agroinfiltration of GFP::AVRcap1b or GFP::PexRD54 (73) and subjected to immunoprecipitation using GFP_Trap_A beads (Chromotek, Munich, Germany), as described previously (50, 96, 99). PexRD54 was included as a control, as it is also a large *P. infestans* RxLR effector and extensive studies suggests that its role is likely independent of the NRC network (73) (74). Final OD_600_ for each protein is indicated in Table S13. Immunoprecipitated samples were separated by SDS-PAGE (4%–20% gradient gel, Biorad) and stained with Coomassie brilliant Blue G-250 (SimplyBlue Safe stain, Invitrogen). Enriched protein samples were cut out of the gel (∼5 × 10 mm) and digested with trypsin. Extracted peptides were analysed by liquid chromatography-tandem mass spectrometry (LC-MS/MS) with the Orbitrap Fusion mass spectrometer and nanoflow-HPLC system U3000 (Thermo Fisher Scientific, UK). (48) (50). A total of three biological replicates for each sample type was submitted.

#### Mass spectrometry data processing

As previously described (49, 50, 96, 99, 100), peak lists were extracted from raw data using MS Convert (101) and peptides identified on Mascot server 2.4.1 using Mascot Daemon (Matrix Science, Ltd.). The peak lists were searched against the *N. benthamiana* genome database called Nicotiana_Benthamiana_Nbv6trPAplusSGNUniq_20170808 (398,682 sequences; 137,880,484 residues), supplemented with common contaminants. The search settings allowed tryptic peptides with up to 2 possible mis-cleavages and charge states +2, +3, +4, carbamidomethylated Cysteine (static) and oxidized Methionine (variable) modifications, and monoisotopic precursor and fragment ion mass tolerances 10ppm and 0.6 Da respectively. A decoy database was used to validate peptide sequence matches. The Mascot results were combined in Scaffold 4.4.0 (Proteome Software Inc.), where thresholds for peptide sequence match and inferred protein were set to exceeded 95.0% and 99% respectively with at least 2 unique peptides identified for each protein. The aforementioned protein list with spectral counts as quantitative values was exported and further analysed in Excel (Microsoft Co.). More information on peptides identified is available in Table S14.

The MS proteomics data have been deposited to the ProteomeXchange Consortium via the PRIDE (102) partner repository with the dataset identifier PXD023178 and 10.6019/PXD023178.

## ACKNOWLEDGEMENTS

We thank Aleksandra ‘Ola’ Bialas (The Sainsbury Laboratory, Norwich, UK) for valuable comments on figures for this paper and Adeline Harant for technical assistance. This work was funded by the Gatsby Charitable Foundation and Biotechnology and Biological Sciences Research Council (BBSRC, UK). S.K. also receives funding from the European Research Council (ERC BLASTOFF projects). L.D. was funded by a Marie Sklodowska-Curie actions fellowship (BoostR), H.A. was funded by the Japan Society for the Promotion of Sciences Postdoctoral fellowship, A.V.C. was funded by a British Society for Plant Pathology summer bursary, J.U. was funded by the Gatsby Charitable Foundation PhD studentship.

## AUTHOR CONTRIBUTIONS

**Conceptualization:** Lida Derevnina, Mauricio P. Contreras, Chih-hang Wu, Sophien Kamoun

**Formal analysis:** Lida Derevnina, Dan MacLean, Jan Sklenar

**Funding acquisition:** Lida Derevnina, Hiroaki Adachi, Jessica Upson, Angel Vergara Cruces, Sophien Kamoun

**Investigation:** Lida Derevnina, Mauricio P. Contreras, Hiroaki Adachi, Rongrong Xie, Abbas Maqbool, Angel Vergara Cruces, Jessica Upson, Chih-hang Wu

**Methodology:** Lida Derevnina, Mauricio P. Contreras, Hiroaki Adachi, Abbas Maqbool, Jan Sklenar, Wenbo Ma, Saskia Hogenhout, Aska Goverse, Chih-hang Wu

**Project administration:** Lida Derevnina, Mauricio P. Contreras, Hiroaki Adachi, Chih-hang Wu, Sophien Kamoun

**Resources:** Lida Derevnina, Hiroaki Adachi, Abbas Maqbool, Angel Vergara Cruces, Jessica Upson, Jan Sklenar, Wenbo Ma, Saskia Hogenhout, Aska Goverse, Chih-hang Wu, Frank L.

H. Menke, Sophien Kamoun

**Supervision:** Lida Derevnina, Abbas Maqbool, Chih-hang Wu, Sophien Kamoun

**Validation:** Lida Derevnina, Mauricio P. Contreras, Abbas Maqbool, Jessica Upson, Sam Mugford

**Visualisation:** Lida Derevnina, Jan Sklenar

**Writing:** Lida Derevnina, Mauricio P. Contreras, Hiroaki Adachi, Abbas Maqbool, Sophien Kamoun

## SUPPLEMENTAL DATA

**Table S1: List of effectors used in this study**.

**Table S2: List of interactors of AVRcap1b identified in Y2H screen**.

**Table S3: List of interactors of SS15 identified in Y2H screen**.

**Table S4: List of interactors of AVRcap1b identified by IP-MS**.

**Table S5: Sequences of *N. benthamiana* TOLs used in this study**.

**Table S6: List of primers used for *P. infestans* effector cloning**.

**Table S7: Sequences of NRC2, NRC3 and NRC4 used in this study**.

**Table S8: List of primers used for NRC2, NRC3, NRC4 and SW5b cloning**.

**Table S9: List of NLRs and corresponding AVRs used in cell death assays**.

**Table S10: List of primers used to generate NbTOL9a hairpin silencing construct**.

**Table S11: List of primers used for cloning NRC Y2H constructs**.

**Table S12: List of primers used to generate NbTOL Golden Gate modules**.

**Table S13: List of final OD**_**600**_ **used in CoIP experiments**.

**Table S14: Spectrum reports for AVRcap1b IP-MS experiments**.

**Table S15: Primers used for cloning NRCs into *E. coli* heterologous expression vectors**.

**Supplementary File 1: Alignment used to generate phylogenetic tree of *A. thaliana* and *N. benthamiana* TOL proteins**.

**Figure S1.**
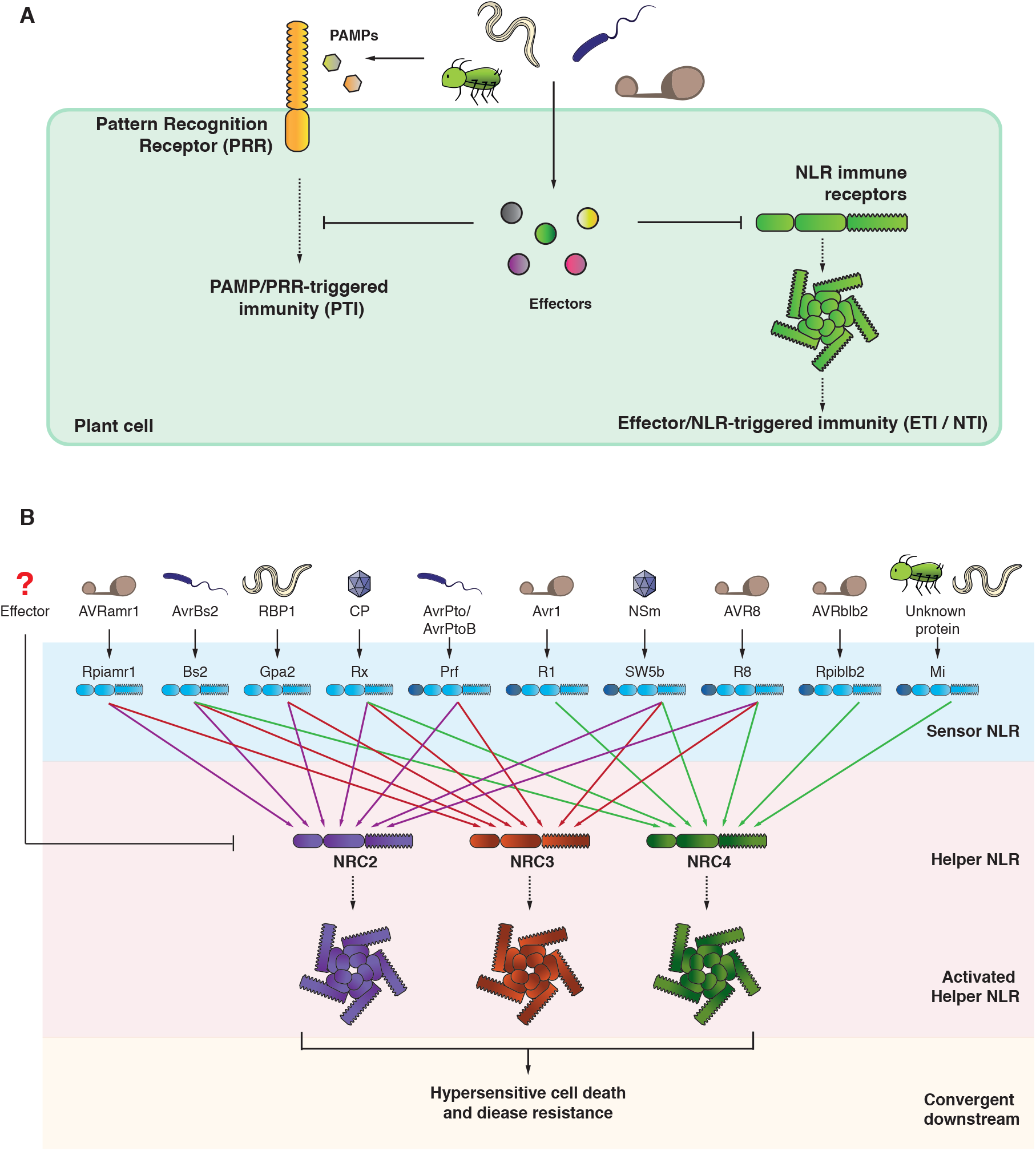
Effectors can suppress Effector- or NLR-Triggered Immunity (ETI / NTI). (A) ETI/NTI suppressing effectors counteract the activation of NLR immunity elicited by other effectors with an avirulence activity (AVR effectors). The majority of effectors studied to date are known to suppress PAMP-Triggered Immunity (PTI) by acting on various host targets involved in PTI signalling. However, we know little about the mechanisms of NTI suppressing effectors and how these effectors counteract NLR receptor functions and/or resistosome formation (B) The NRC network of Solanaceae. Helper NLRs (NRC2, NRC3, NRC4) function redundantly with a series of R genes that confer resistance against multiple pathogens. These R genes encode NLR sensors that have specialized in detecting effectors from pathogens as diverse as oomycetes, bacteria, nematodes, viruses and aphids. Many of the NRC-dependent R genes are agronomically important. Sensor NLRs (blue) with N-terminal extensions possess an additional domain at the N-terminus, represented in dark blue. The extent to which pathogen effectors have evolved to target the NRC helpers are currently unknown. Figure is adapted from Wu et al. PNAS 2017 [24, 94].

**Figure S2.**
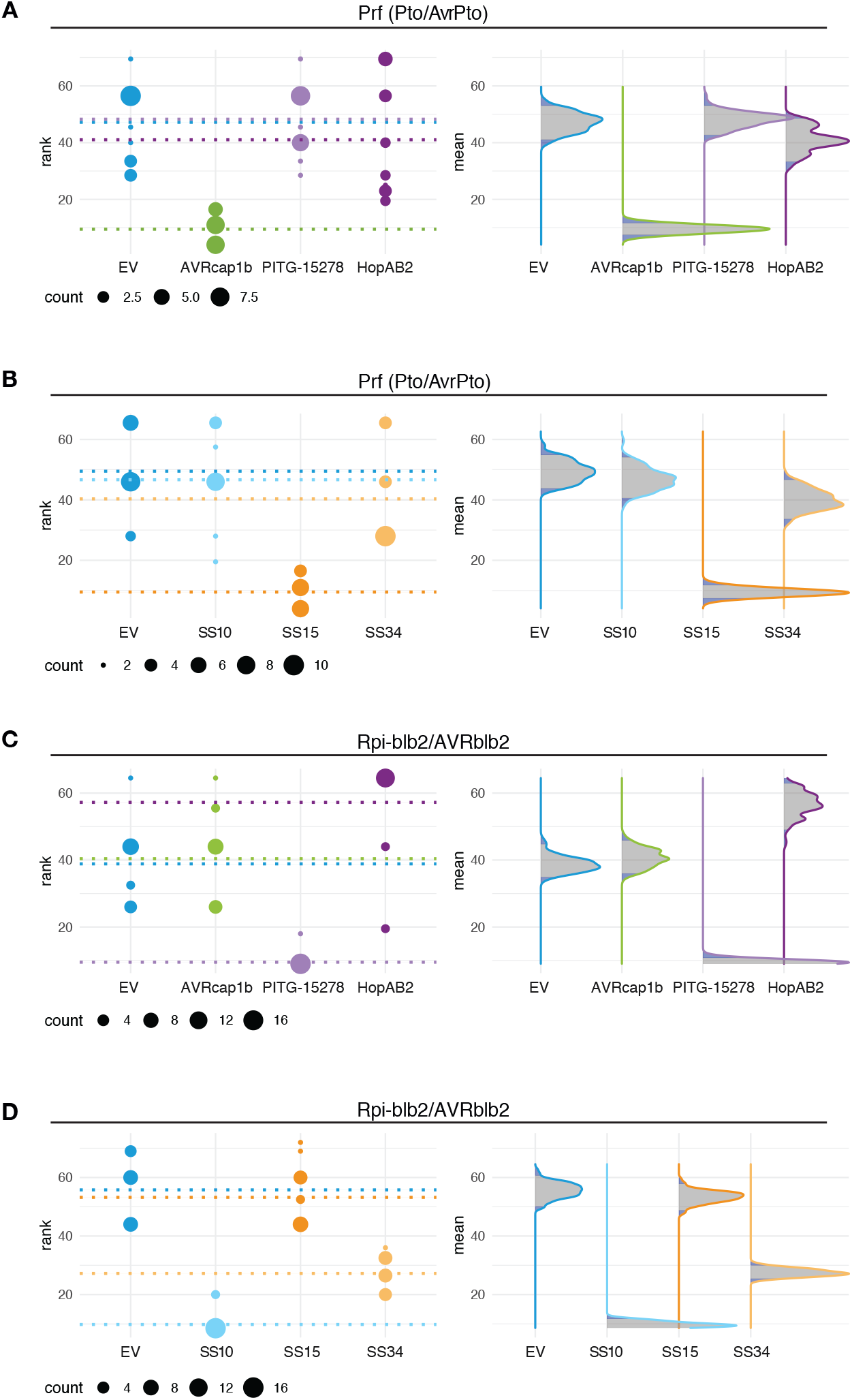
Statistical analysis of cell death suppression assay for five candidate effectors co-infiltrated with Pto/AvrPto or Rpi-blb2/AVRblb2. Statistical analysis was conducted using besthr R library (MacLean, 2019). (A–D) Each panel corresponds to results from Pto/AvrPto or Rpi-blb2/AVRblb2 (labelled above), co-expressed with either empty vector (EV, dark blue), AVRcap1b (green), HopAB2 (dark purple), PITG-15278 (light purple), SS10 (light blue), SS15 (dark orange), or SS34 (light orange), where EV was used as a negative control, labelled below plots. The left panel represents the ranked data (dots) and their corresponding means (dashed lines), with the size of each dot proportional to the number of observations for each specific value (count key below each panel). The panels on the right shows the distribution of 1000 bootstrap sample rank means, where the blue areas under the curve illustrates the 0.025 and 0.975 percentiles of the distribution. A difference is considered significant if the ranked mean for a given condition falls within or beyond the blue percentile of the mean distribution of the EV control.

**Figure S3.**
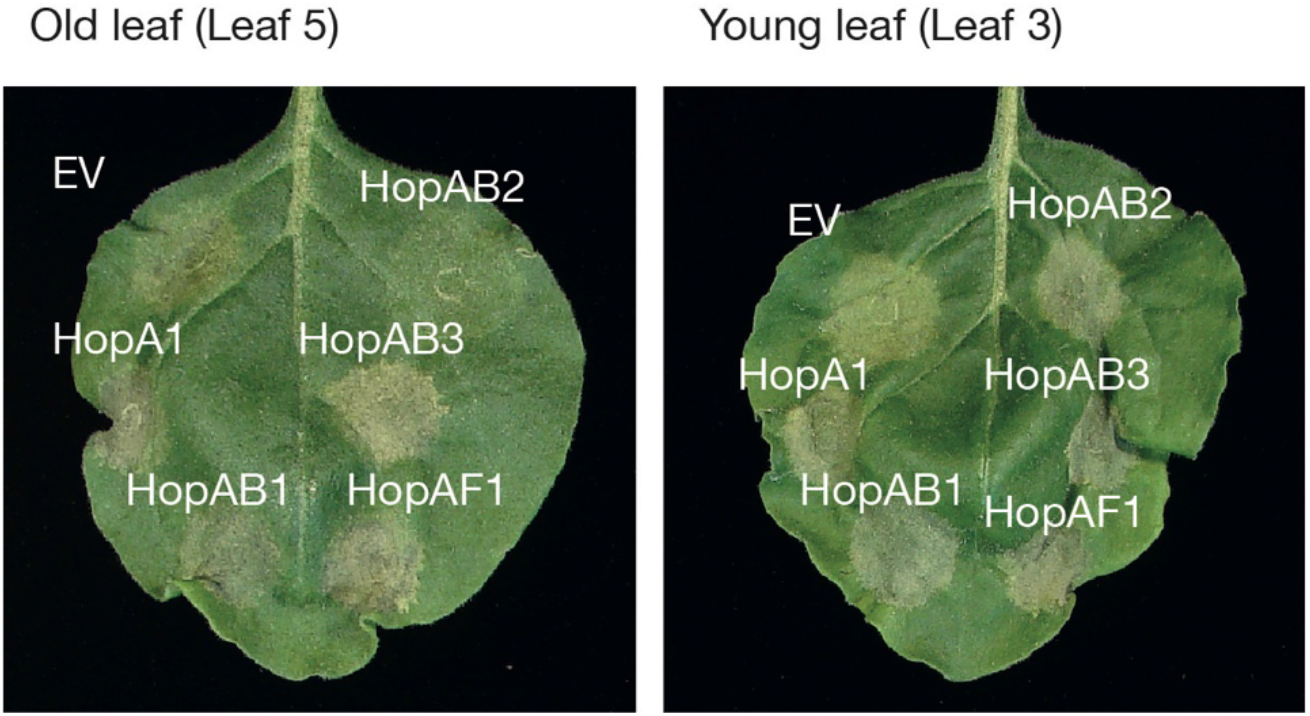
HopAB2 suppression of Prf-mediated cell death in *N. benthamiana* is not evident in young leaves. *Psuedomonas syringae* effectors HopA1, HopAB1, HopAB2, HopAB3, HopAF1, were transiently co-expressed with Pto/AvrPto in *N. benthamiana* leaves. The leaves were photographed 5 days after infiltration. HopAB2 suppression of Pto/AvrPto in *N. benthamiana* were evident on leaf 5 (old leaf, left), but not on leaf 3 (young leaf, right) as defined by [39]. Photographs are representative of 3 technical repeats.

**Figure S4.**
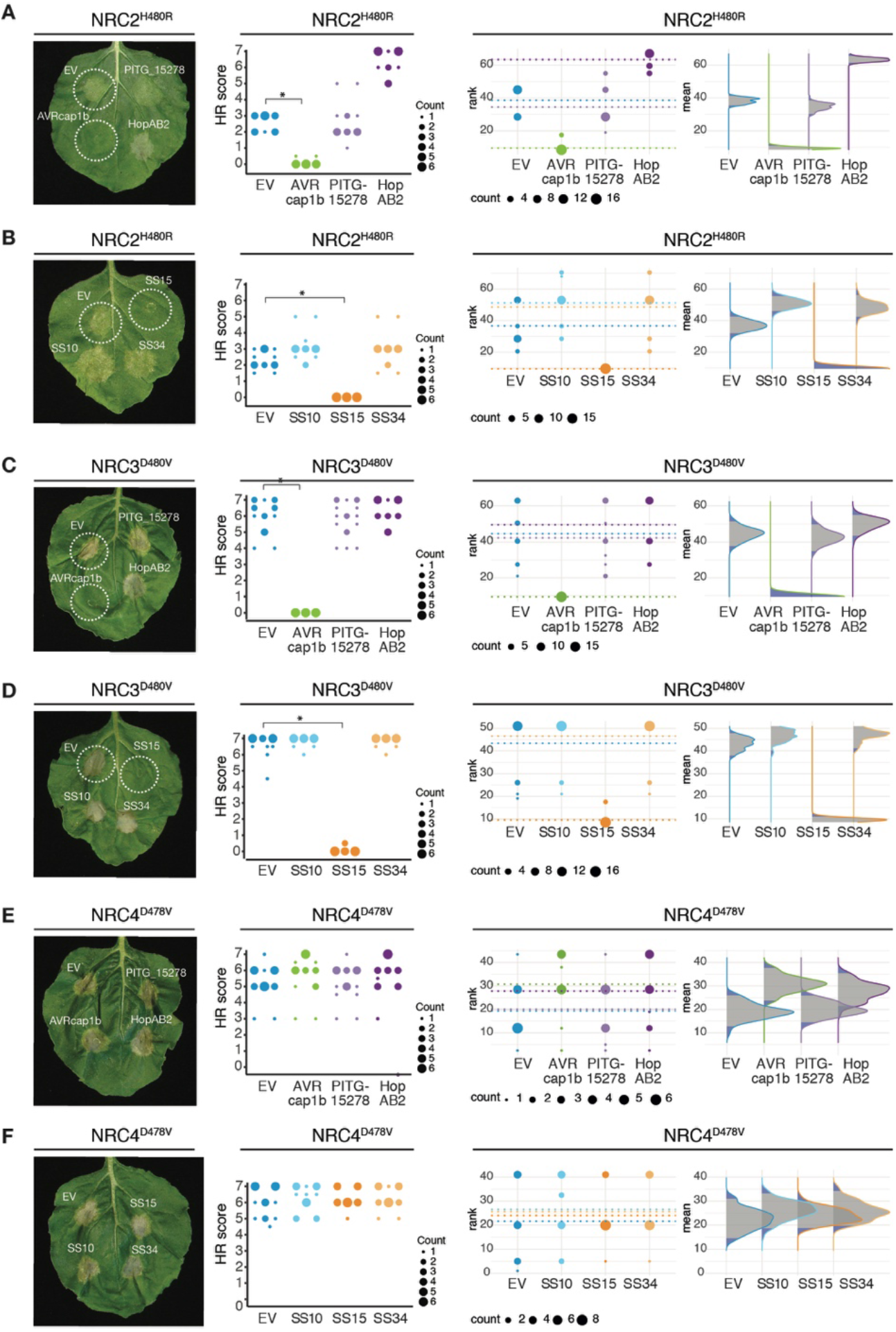
AVRcap1b and SS15 suppress cell death activity mediated by autoactive NRC2^H480R^ and NRC3^D480V^, but not of NRC4^D478V^ mutants. (A – F) Leaf panels: Photo of representative *Nicotiana benthamiana* leaves showing HR after co-expression of autoimmune NRC2^H480R^, NRC3^D480V^, and NRC4^D478V^ mutants, indicated above leaf panels, with either Empty vector, AVRcap1b, PITG_15278, HopAB2, SS10, SS15, or SS34 (labelled on leaf). Middle panels: Hypersensitive Response (HR) was scored 5 days post-agroinfiltration, using a modified 0 - 7 scale (Segretin et al 2014) and photographed at 5 days after agroinfiltration. HR results are presented as dot plots, where the size of each dot is proportional to the number of samples with the same score (count) within each biological replicate. Each effector is represented by a different dot colour (see key right side of plot). The experiment was independently repeated three times each with six technical replicates. The columns of each tested effector correspond to results from different biological replicates. Right panels: Statistical analysis was conducted using besthr R library (MacLean, 2019). The dots represent the ranked data and their corresponding means (dashed lines), with the size of each dot proportional to the number of observations for each specific value (count key below each panel). The panels on the right show the distribution of 1000 bootstrap sample rank means, where the blue areas under the curve illustrates the 0.025 and 0.975 percentiles of the distribution. A difference is considered significant if the ranked mean for a given condition falls within or beyond the blue percentile of the mean distribution of the EV control. Significant differences between the conditions are indicated with an asterisk (*).

**Figure S5.**
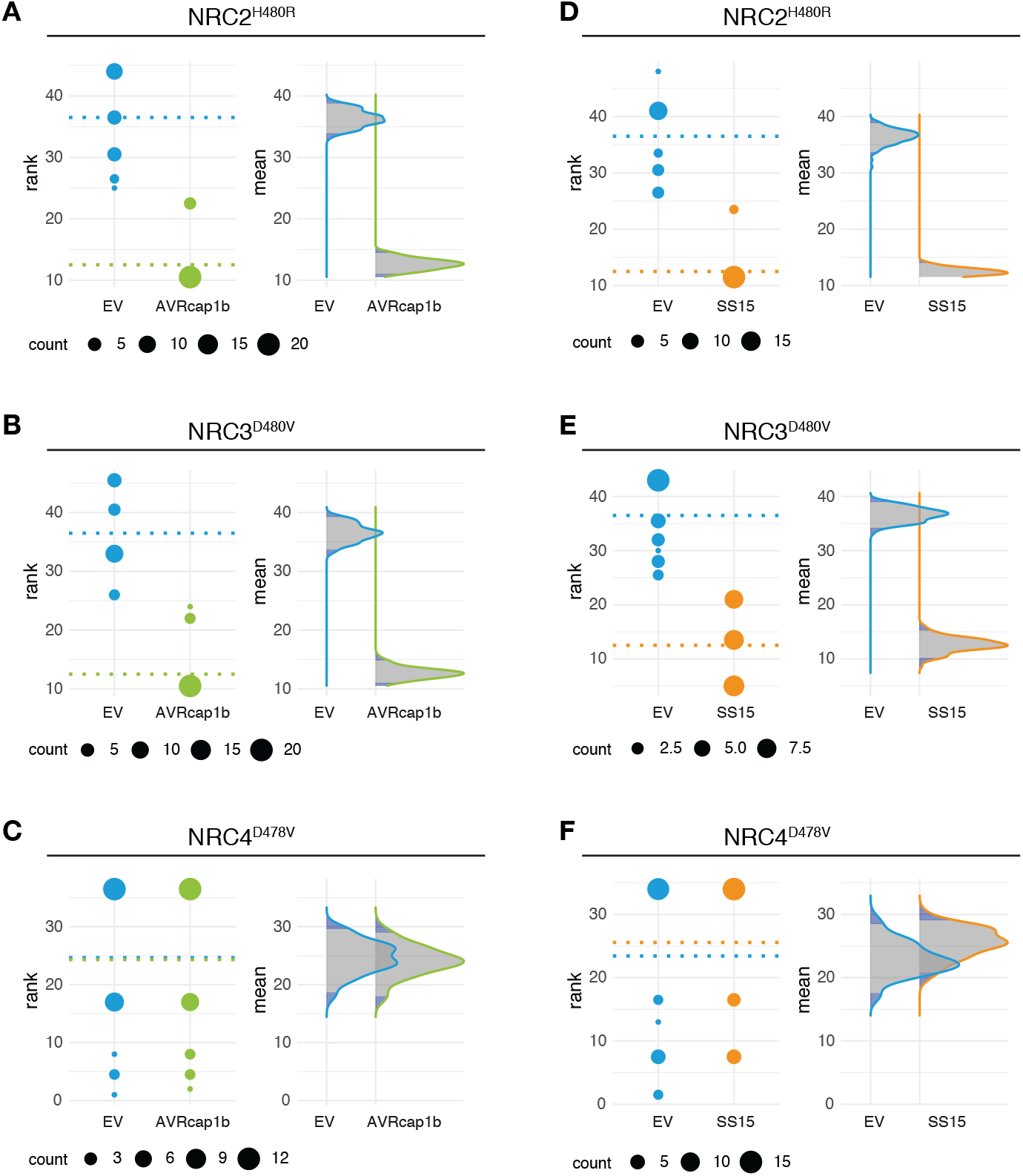
Statistical analysis of cell death suppression assay for AVRcap1b and SS15 co-expressed with autoactive NRC2^H480R^, NRC3^D480V^ and NRC4^D478V^. Statistical analysis was conducted using besthr R library (MacLean, 2019). Each panel corresponds to results from (A, D) NRC2^H480R^, (B, E) NRC3^D480V^ or (C, F) NRC4^D478V^ (labelled above), co-expressed with either empty vector (EV, blue), AVRcap1b (green) or SS15 (orange), where EV was used as a negative control (labelled under plots). The left panel represents the ranked data (dots) and their corresponding means (dashed lines), with the size of each dot proportional to the number of observations for each specific value (count key below each panel). The panels on the right show the distribution of 1000 bootstrap sample rank means, where the blue areas under the curve illustrates the 0.025 and 0.975 percentiles of the distribution. A difference is considered significant if the ranked mean for a given condition falls within or beyond the blue percentile of the mean distribution of the EV control.

**Figure S6.**
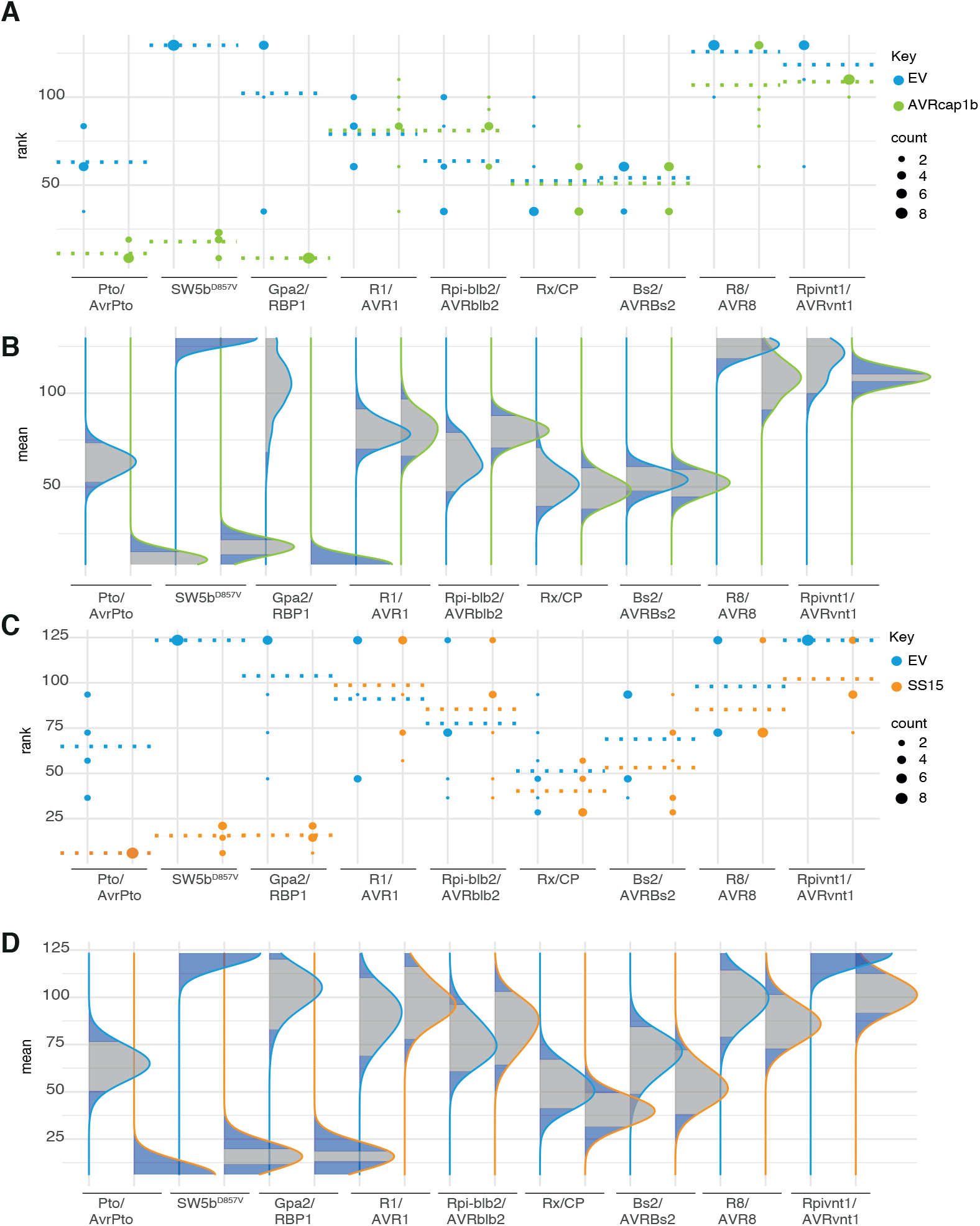
Statistical analysis of suppression of NRC2/3 dependent sensor NLRs by AVRcap1b or SS15. Statistical analysis was conducted using besthr R library (MacLean, 2019). (A–D) Within each panel results from a range of different sensor NLRs (labeled below) were co-expressed with either empty vector (EV, blue), AVRcap1b (green), or SS15 (orange), where EV was used as a negative control, see key on right hand side. Panels (A) and (C) represent the ranked data (dots) and their corresponding means (dashed lines), with the size of each dot proportional to the number of observations for each specific value (refer to count key, on left hand side). Panels (B) and (D) show the distribution of 1000 bootstrap sample rank means, where the blue areas under the curve illustrates the 0.025 and 0.975 percentiles of the distribution. A difference is considered significant if the ranked mean for a given condition falls within or beyond the blue percentile of the mean distribution of EV control for each treatment.

**Fig S7.**
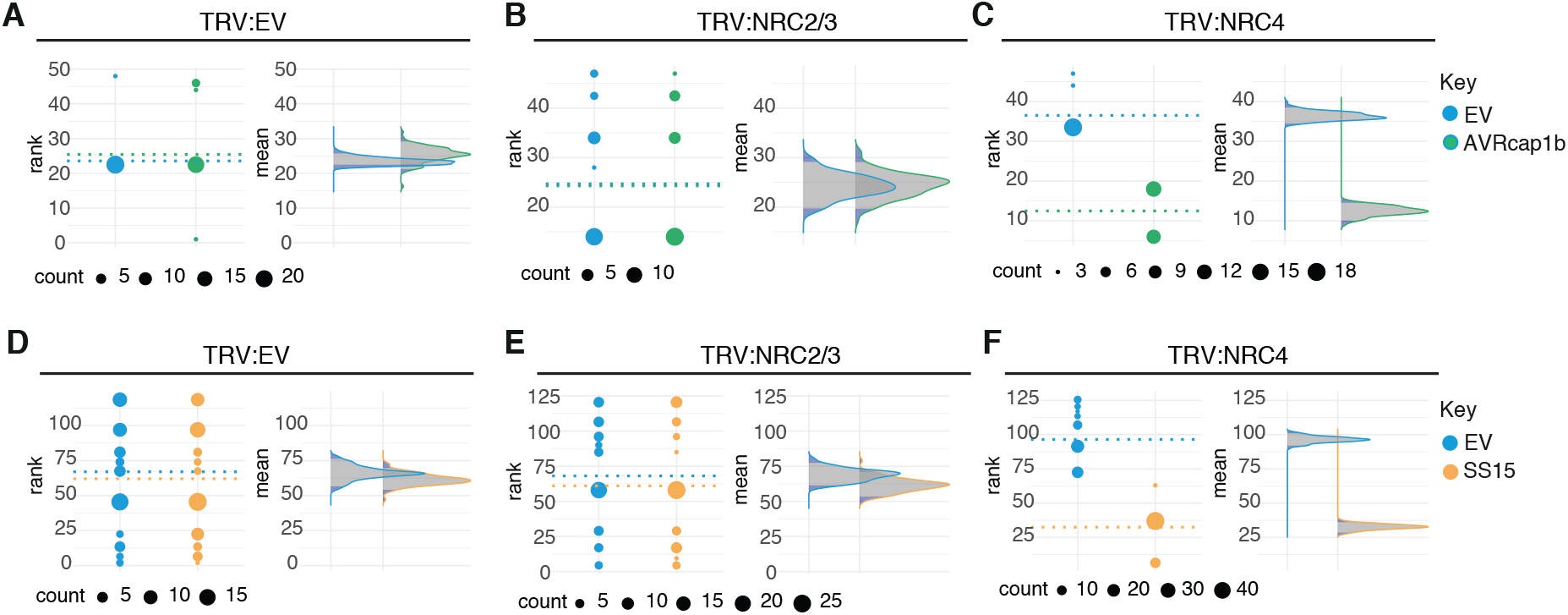
Statistical analysis shows that AVRcap1b and SS15 suppress Rx-mediated cell death in NRC4 silenced Rx transgenic *Nicotiana benthamiana* plants. Statistical analysis was conducted using besthr R library (MacLean, 2019). (A – F) Each panel represents a different silencing treatment (labelled above). Left plot represents the ranked data (dots) and their corresponding means (dashed lines), with the size of each dot proportional to the number of observations for each specific value (refer to count key below each individual panel). Right plot shows the distribution of 1000 bootstrap sample rank means, where the blue areas under the curve illustrates the 0.025 and 0.975 percentiles of the distribution. A difference is considered significant if the ranked mean for a given condition falls within or beyond the blue percentile of the mean distribution of EV control.

**Figure S8.**
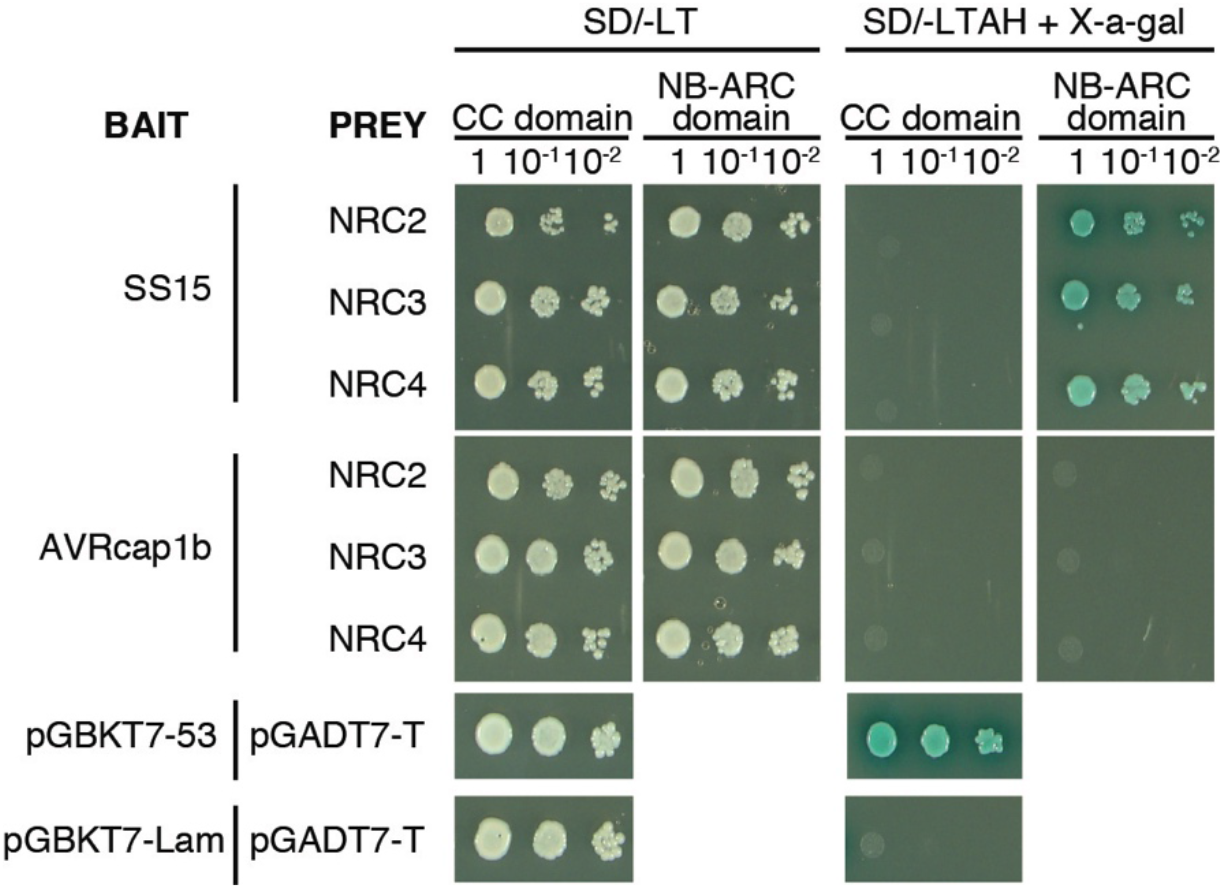
SS15 binds the NB-ARC domain of NRC proteins in yeast. Yeast two-hybrid assay of SS15 and AVRcap1b with CC and NB-ARC domains of NRC2, NRC3 and NRC4. Control plates for yeast growth are on the left (SD-Leu-Trp, ST0048, Takara Bio, USA), with quadruple dropout media supplemented with X-a-gal on the right (SD-Leu-Trp-Ade-His, ST0054, Takara Bio, USA). The commercial yeast constructs were used as positive (pGBKT7-53/pGADT7-T) and negative (pGBKT7-Lam/pGADT7-T) controls. Growth and development of blue colouration in the selection plate are both indicative of protein-protein interaction. SS15 and AVRcap1b were fused to the GAL4 DNA binding domain (Bait) and the truncated NRCs were fused to the GAL4-activator domain (Prey). Each experiment was repeated three times with similar results.

**Figure S9.**
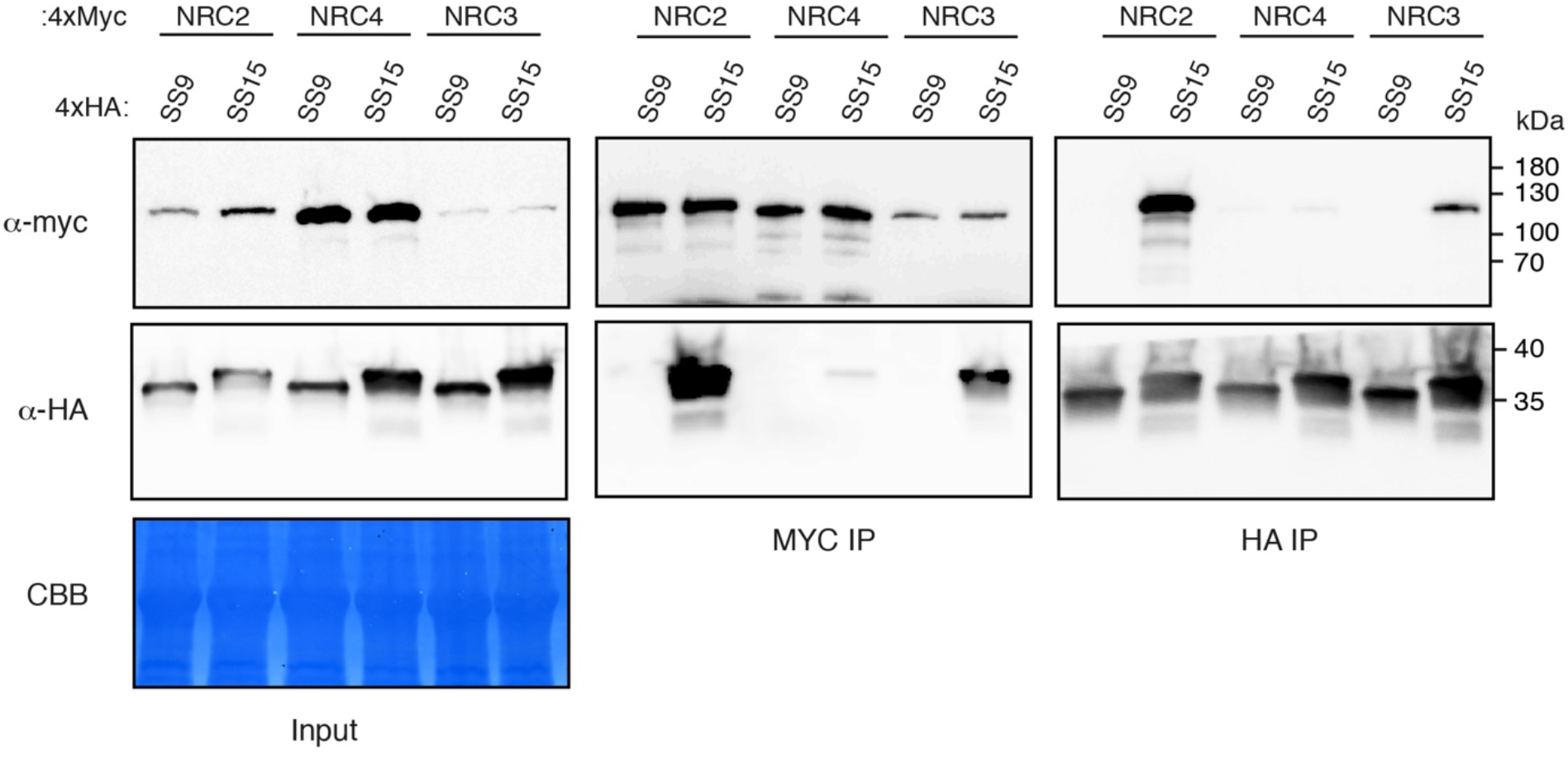
Co-immunoprecipitation experiment between SS15 and NRC2, NRC3 and NRC4. N-terminally 4xHA-tagged SS15 (SS15) and SS9 (SS9) were individually transiently co-expressed with NRC proteins fused to a C-terminal 4xMyc tag. SS9 was used as a negative control. Immunoprecipitation (IP) were performed with agarose beads conjugated to either anti-Myc (MYC IP) or anti-HA (HA IP) antibodies. Total protein extracts were immunoblotted with appropriate antisera labelled on the left. Approximate molecular weights (kDa) of the proteins are shown on the right. Rubisco loading controls were conducted using Coomassie Brilliant Blue (CBB) staining. This experiment is representative of three independent replicates.

**Figure S10.**
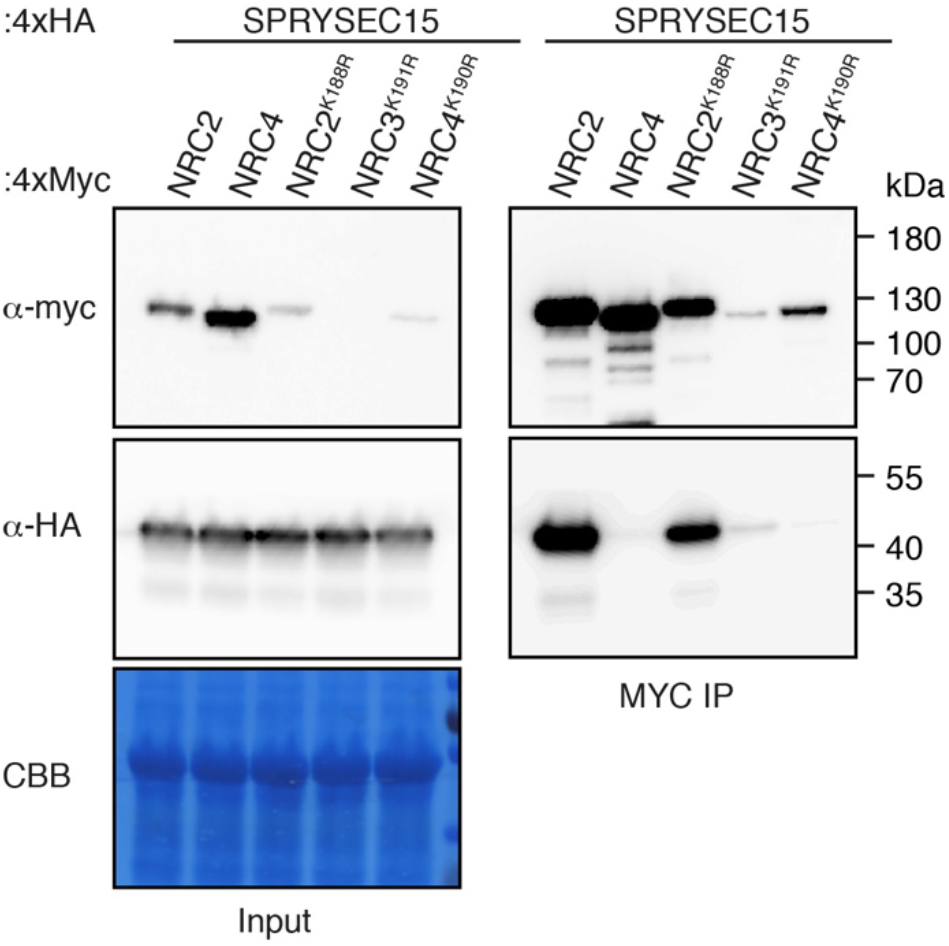
SS15 associates with NRC2, NRC3 and NRC4 p-loop mutants in planta. (A) N-terminally 4xHA tagged SS15 was transiently co-expressed in *N. benthamiana* with C-terminally 4xMyc tagged NRC2, NRC4, NRC2^K188R^, NRC3^K191R^, NRC4^K190R^. IP was performed with agarose beads conjugated to anti-Myc (Myc IP) antibodies. Total protein extracts (Input) and proteins obtained by co-immunoprecipitation were immunoblotted with appropriate antisera labelled on the right. Approximate molecular weights (kDa) of the proteins are shown on the right. Rubisco loading controls were conducted using Coomassie Brilliant Blue (CBB) staining. This experiment is representative of three independent replicates.

**Figure S11:**
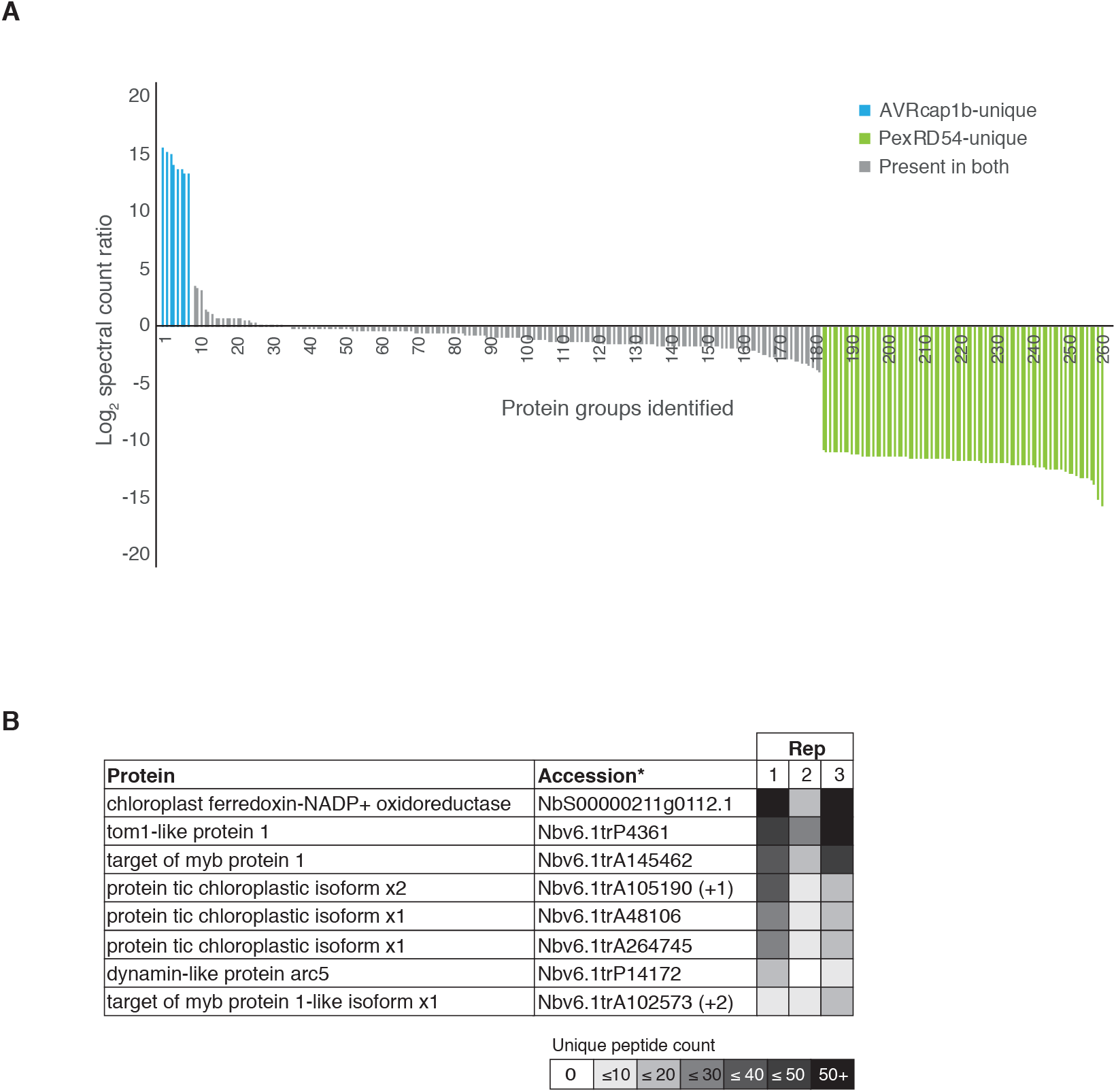
Target of AVRcap1b identified via immunoprecipitation-mass spectrometry. (A) Immune-affinity enriched protein abundance plot generated based on proteins groups identified for AVRcap1b and PexRD54. Protein abundances were calculated as the total mean spectral counts identified with each effector across three replicates. The Log_2_ of the ratio of mean counts observed for AVRcap1b divided by the mean counts observed for PexRD54 was calculated. All 261 proteins identified were then sorted from highest to lowest ratio and plotted in this order (see Table S4 for details). Proteins with the highest ratio correspond to the most likely AVRcap1b (blue) specific interactors. Proteins most likely to correspond to PexRD54 are labelled in green. Proteins in grey were identified in both effectors. (B) Eight putative unique AVRcap1b proteins (based on previously mentioned ratio of mean spectral counts) identified in immunoprecipitation mass spectrometry experiment. Unique spectral counts obtained in each of the three independent replicates with AVRcap1b are included on the right-hand side.

**Figure S12:**
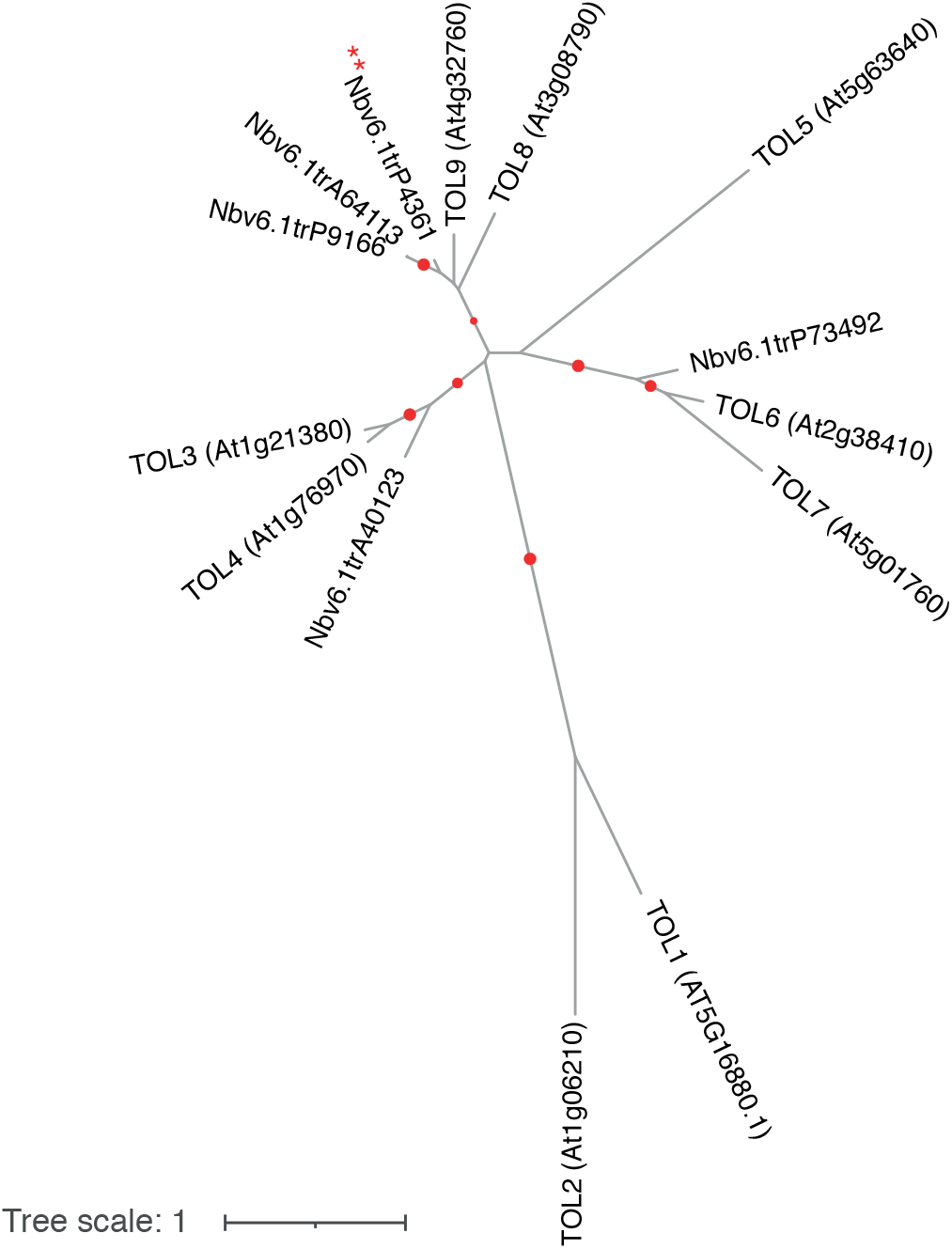
Maximum-likelihood phylogenetic tree of TOL proteins from *N. benthamiana* and *A. thaliana*. Protein sequences were aligned using Clustal Omega. The ENTH and GAT domains were used for further analysis. The phylogenetic tree was constructed in MEGAX using the Jones-Taylor-Thornton (JTT) substitution model and 1000 bootstrap iterations. Branches with bootstrap support higher than 80 are indicated with red dots. NbTOL9a is indicated by two red asterisks. The scale bar indicates the evolutionary distance in amino acid substitutions per site.

**Figure S13.**
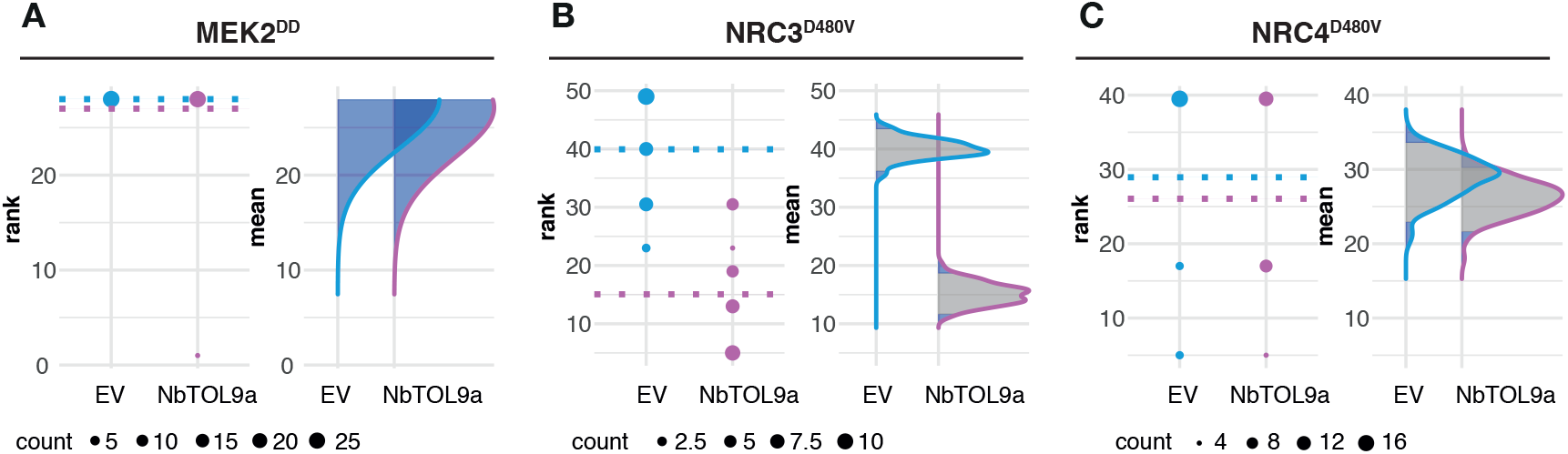
Statistical analysis of HR mediated by autoactive NRC3^D480V^, but not MEK2^DD^ or NRC4^D478V^ is reduced when NbTOL9a is overexpressed. Statistical analysis was conducted using besthr R library (MacLean, 2019). For (A) MEK2^DD^, (B) NRC3^D480V^ and (C) NRC4^D478V^ the left most panel represents the ranked data (dots) and their corresponding means (dashed lines), with the size of each dot proportional to the number of observations for each specific value (refer to count legend bottom of figure). The right panel shows the distribution of 1000 bootstrap sample rank means, where the blue areas under the curve illustrates the 0.025 and 0.975 percentiles of the distribution. A difference is considered significant if the ranked mean for AVRcap1b (purple) falls within or beyond the blue percentile of the mean distribution of EV control (blue) for each treatment (MEK2^DD^, NRC3^D480V^, NRC4^D478V^), in the presence of either EV or NbTOL9a.

**Figure S14.**
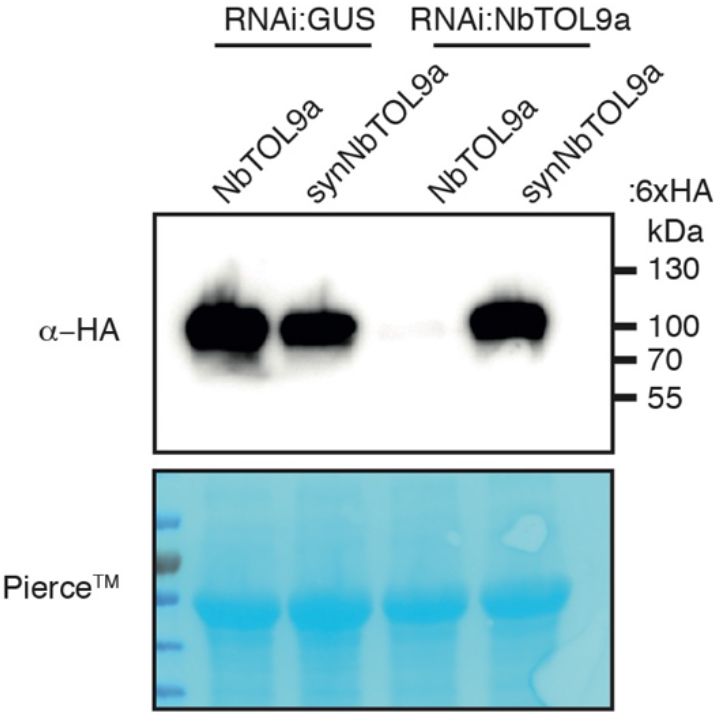
RNAi::*NbTOL9a* silencing construct effectively reduces protein accumulation levels of NbTOL9a. Wild-type NbTOL9a (NbTOL9a::6HA) and the synthetic version (synNbTOL9a::6xHA) were transiently co-expressed with RNAi::*GUS* or RNAi:*NbTOL9a*. synNbTOL9a::6xHA was used as a control that is not targeted for knockdown by the RNAi::*NbTOL9a*. Total protein extracts were immunoblotted with HA antiserum. Approximate molecular weights (kDa) of the proteins are shown on the right. Rubisco loading controls were conducted using Pierce™ staining.

**Figure S15.**
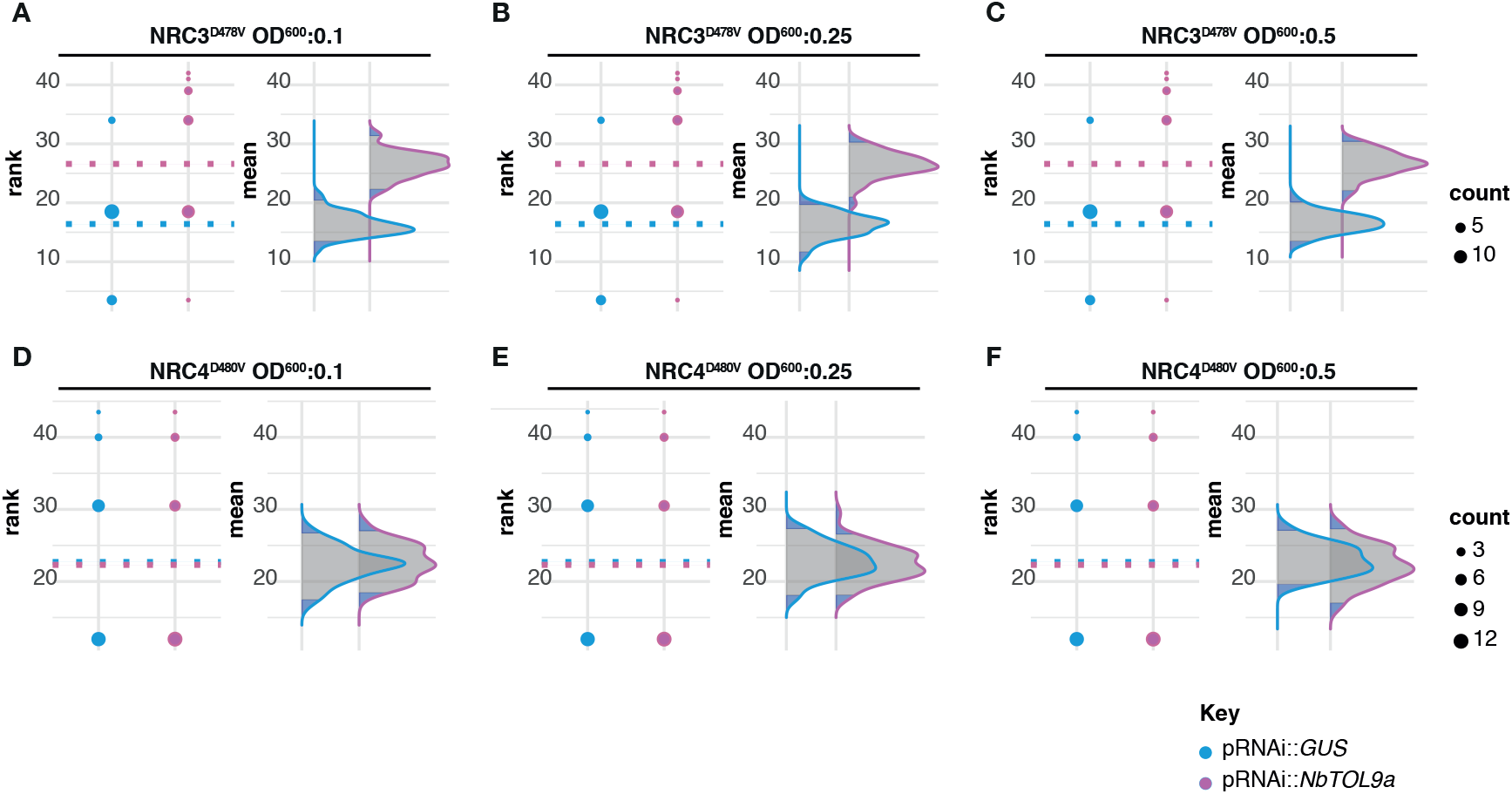
Statistical analysis showing silencing of NbTOL9a enhanced cell death mediated by NRC3^D480V^ but not NRC4^D478V^. Statistical analysis was conducted using besthr R library (MacLean, 2019). For (A - C) NRC3^D480V^ and (D - F) NRC4^D478V^, each panel represents increasing concertation of *A. tumefaciens* expressing NRC3^D480V^ and NRC4^D478V^ (OD_600_ = 0.1, 0.25 or 0.5). The left most panel represents the ranked data (dots) and their corresponding means (dashed lines), with the size of each dot proportional to the number of observations for each specific value (refer to count legend bottom of figure). The right panel shows the distribution of 1000 bootstrap sample rank means, where the blue areas under the curve illustrates the 0.025 and 0.975 percentiles of the distribution. A difference is considered significant if the ranked mean for RNAi::NbTOL9a (purple) falls within or beyond the blue percentile of the mean distribution of RNAi::GUS (blue) for each treatment combination.

**Figure S16.**
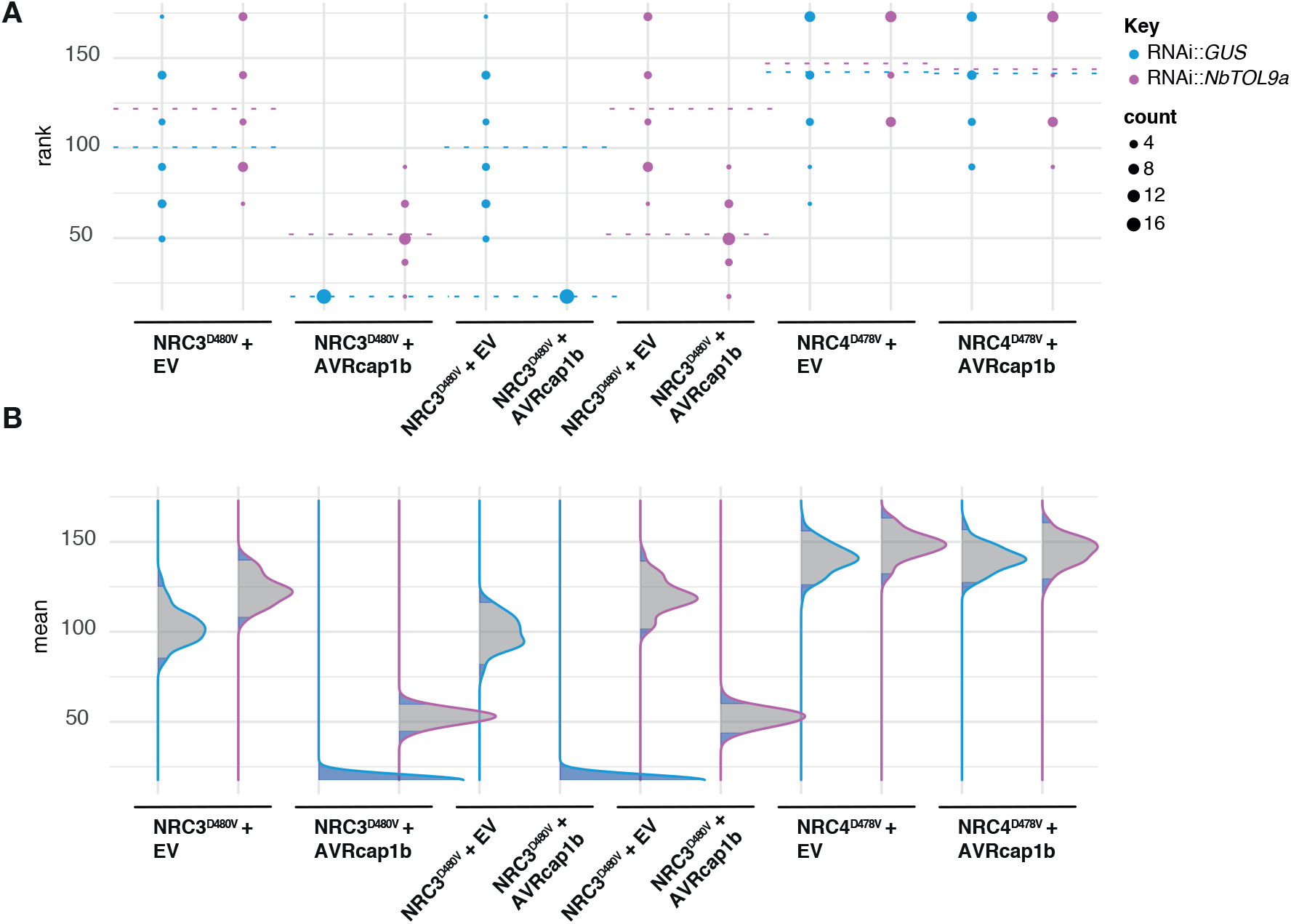
Statistical analysis of NRC3^D480V^ cell death suppression mediated by AVRcap1b in NbTOL9a silencing. Statistical analysis was conducted using besthr R library (MacLean, 2019). Autoactive NRC3 (NRC3^D480V^) was co-expressed with either empty vector (EV, negative control) or AVRcap1b (labelled on bottom of plot), with RNAi::GUS (blue) or RNAi::NbTOL9a (purple) silencing treatments. (A) represents the ranked data (dots) and their corresponding means (dashed lines), with the size of each dot proportional to the number of observations for each specific value (refer to count key, on right hand side). (B) shows the distribution of 1000 bootstrap sample rank means, where the blue areas under the curve illustrates the 0.025 and 0.975 percentiles of the distribution. A difference is considered significant if the ranked mean for a given condition falls within or beyond the blue percentile of the mean distribution of the corresponding RNAi::GUS controls within each given treatment (NRC3^D480V^ + EV, NRC3^D480V^ + AVRcap1b, NRC4^D478V^ + EV, NRC4^D478V^ + AVRcap1b).

## REFERENCES

1. Win J, Chaparro-Garcia A, Belhaj K, Saunders DG, Yoshida K, Dong S, et al. Effector biology of plant-associated organisms: concepts and perspectives. Cold Spring Harb Symp Quant Biol. 2012. 77:235–47. doi: 10.1101/sqb.2012.77.015933.

2. Toruno TY, Stergiopoulos I, Coaker G. Plant-pathogen effectors: cellular probes interfering with plant defenses in spatial and temporal manners. Annual Reviews of Phytopathology. 2016. 54:419 – 41. doi: https://doi.org/10.1146/annurev-phyto-080615-100204.

3. Macho AP, Zipfel C. Targeting of plant pattern recognition receptor-triggered immunity by bacterial type-III secretion system effectors. Curr Opin Microbiol. 2015. 23:14–22. doi: 10.1016/j.mib.2014.10.009.

4. Zheng X, Wagener N, McLellan H, Boevink PC, Hua C, Birch PRJ, et al. Phytophthora infestans RXLR effector SFI5 requires association with calmodulin for PTI/MTI suppressing activity. New Phytol. 2018. 219:1433–46. doi: 10.1111/nph.15250.

5. Cui H, Tsuda K, Parker JE. Effector-Triggered Immunity: from pathogen perception to robust defense. Annual Reviews of Plant Biology. 2015. 66:487 – 511. doi: https://doi.org/10.1146/annurev-arplant-050213-040012.

6. Jones JD, Vance RE, Dangl JL. Intracellular innate immune surveillance devices in plants and animals. Science. 2016. 354. doi: 10.1126/science.aaf6395.

7. Zhang X, Dodds PN, Bernoux M. What do we know about NOD-like receptors in plant immunity? Annual Review of Phytopathology. 2017. 55:205–29. doi: 10.1146/annurev-phyto-080516-035250 %M 28637398 %U.

8. Cesari S. Multiple strategies for pathogen perception by plant immune receptors. New Phytol. 2018. 219:17–24. doi: 10.1111/nph.14877.

9. Kourelis J, van der Hoorn RAL. Defended to the nines: 25 years of resistance gene cloning identifies nine mechanisms for R protein function. The Plant Cell. 2018. 30:285. doi: 10.1105/tpc.17.00579.

10. Dodds PN, Rathjen JP. Plant immunity: towards an integrated view of plant-pathogen interactions. Nat Rev Genet. 2010. 11:539–48. doi: 10.1038/nrg2812.

11. Jones JD, Dangl JL. The plant immune system. Nature. 2006. 444:323–9. doi: 10.1038/nature05286.

12. Thordal-Christensen H. A holistic view on plant effector-triggered immunity presented as an iceberg model. Cellular and Molecular Life Sciences. 2020. 77:3963–76. doi: 10.1007/s00018-020-03515-w.

13. Bos JIB, Kanneganti T-D, Young C, Cakir C, Huitema E, Win J, et al. The C-terminal half of Phytophthora infestans RXLR effector AVR3a is sufficient to trigger R3a-mediated hypersensitivity and suppress INF1-induced cell death in Nicotiana benthamiana. The Plant Journal. 2006. 48:165–76. doi: https://doi.org/10.1111/j.1365-313X.2006.02866.x.

14. D’Ambrosio JM, Couto D, Fabro G, Scuffi D, Lamattina L, Munnik T, et al. Phospholipase C2 affects MAMP-triggered immunity by modulating ROS production. Plant physiology. 2017. 175:970–81. doi: 10.1104/pp.17.00173.

15. Harris JM, Balint-Kurti P, Bede JC, Day B, Gold S, Goss EM, et al. What are the top 10 unanswered questions in molecular plant-microbe interactions? Molecular Plant-Microbe Interactions. 2020. 33:1354–65. doi: 10.1094/MPMI-08-20-0229-CR.

16. Kourelis J, Kamoun S. RefPlantNLR: a comprehensive collection of experimentally validated plant NLRs. bioRxiv. 2020. doi: https://doi.org/10.1101/2020.07.08.193961.

17. Shao ZQ, Xue JY, Wu P, Zhang YM, Wu Y, Hang YY, et al. Large-scale analyses of angiosperm nucleotide-binding site-leucine-rich repeat genes reveal three anciently diverged classes with distinct evolutionary patterns. Plant Physiol. 2016. 170:2095–109. doi: 10.1104/pp.15.01487.

18. Tamborski J, Krasileva KV. Evolution of plant NLRs: from natural history to precise modifications. Annual Reviews of Plant Biology. 2020. 71:355 – 78.

19. Lee H-Y, Mang H, Choi E-H, Seo Y-E, Kim M-S, Oh S, et al. Genome-wide functional analysis of hot pepper immune receptors reveals an autonomous NLR cluster in seed plants. bioRxiv. 2020. doi: https://doi.org/10.1101/2019.12.16.878959.

20. Lee H-Y, Mang H, Choi E, Seo Y-E, Kim M-S, Oh S, et al. Genome-wide functional analysis of hot pepper immune receptors reveals an autonomous NLR clade in seed plants. New Phytologist. 2020. 229: 532–547. doi: https://doi.org/10.1111/nph.16878.

21. Bentham AR, Zdrzalek R, De la Concepcion JC, Banfield MJ. Uncoiling CNLs: Structure/function approaches to understanding CC domain function in plant NLRs. Plant Cell Physiol. 2018. 59:2398–408. doi: 10.1093/pcp/pcy185.

22. Bentham A, Burdett H, Anderson PA, Williams SJ, Kobe B. Animal NLRs provide structural insights into plant NLR function. Ann Bot. 2017. 119:827–702. doi: 10.1093/aob/mcw171.

23. Sarris PF, Cevik V, Dagdas G, Jones JD, Krasileva KV. Comparative analysis of plant immune receptor architectures uncovers host proteins likely targeted by pathogens. BMC Biol. 2016. 14:8. doi: 10.1186/s12915-016-0228-7.

24. Adachi H, Derevnina L, Kamoun S. NLR singletons, pairs, and networks: evolution, assembly, and regulation of the intracellular immunoreceptor circuitry of plants. Curr Opin Plant Biol. 2019. 50:121–31. doi: 10.1016/j.pbi.2019.04.007.

25. Wu CH, Abd-El-Haliem A, Bozkurt TO, Belhaj K, Terauchi R, Vossen JH, et al. NLR network mediates immunity to diverse plant pathogens. Proc Natl Acad Sci U S A. 2017. 114:8113–8. doi: 10.1073/pnas.1702041114.

26. Adachi H, Contreras MP, Harant A, Wu CH, Derevnina L, Sakai T, et al. An N-terminal motif in NLR immune receptors is functionally conserved across distantly related plant species. eLife. 2019. 8. doi: 10.7554/eLife.49956.

27. Wu CH, Belhaj K, Bozkurt TO, Birk MS, Kamoun S. Helper NLR proteins NRC2a/b and NRC3 but not NRC1 are required for Pto-mediated cell death and resistance in Nicotiana benthamiana. New Phytologist. 2016. 209:1344–52.

28. Martin R, Qi T, Zhang H, Liu F, King M, Toth C, et al. Structure of the activated ROQ1 resistosome directly recognizing the pathogen effector XopQ. Science. 2020. 370(6521):eabd9993. doi: 10.1126/science.abd9993.

29. Wang J, Hu M, Wang J, Qi J, Han Z, Wang G, et al. Reconstitution and structure of a plant NLR resistosome conferring immunity. Science. 2019. 364:aav5870. doi: 10.1126/science.aav5870.

30. Wang J, Wang J, Hu M, Wu S, Qi J, Wang G, et al. Ligand-triggered allosteric ADP release primes a plant NLR complex. Science. 2019. 364: aav5868. doi: 10.1126/science.aav5868.

31. Ma S, Lapin D, Liu L, Sun Y, Song W, Zhang X, et al. Direct pathogen-induced assembly of an NLR immune receptor complex to form a holoenzyme. Science. 2020. 370:eabe3069. doi: 10.1126/science.abe3069.

32. He J, Ye W, Choi DS, Wu B, Zhai Y, Guo B, et al. Structural analysis of Phytophthora suppressor of RNA silencing 2 (PSR2) reveals a conserved modular fold contributing to virulence. Proc Natl Acad Sci U S A. 2019. 116:8054–9. doi: 10.1073/pnas.1819481116.

33. Rietman H. Putting the Phytophthora infestans genome sequence at work: multiple novel avirulence and potato resistance gene candidates revealed. https://library.wur.nl/WebQuery/wurpubs/406778: Wageningen University. 2011.

34. Yoshida K, Schuenemann VJ, Cano LM, Pais M, Mishra B, Sharma R, et al. The rise and fall of the Phytophthora infestans lineage that triggered the Irish potato famine. eLife. 2013. 2:e00731. doi: 10.7554/eLife.00731.

35. Haas BJ, Kamoun S, Zody MC, Jiang RH, Handsaker RE, Cano LM, et al. Genome sequence and analysis of the Irish potato famine pathogen Phytophthora infestans. Nature. 2009. 461:393–8. doi: 10.1038/nature08358.

36. Ali S, Magne M, Chen S, Obradovic N, Jamshaid L, Wang X, et al. Analysis of Globodera rostochiensis effectors reveals conserved functions of SPRYSEC proteins in suppressing and eliciting plant immune responses. Front Plant Sci. 2015. 6:623. doi: 10.3389/fpls.2015.00623.

37. Postma WJ, Slootweg EJ, Rehman S, Finkers-Tomczak A, Tytgat TO, van Gelderen K, et al. The effector SPRYSEC-19 of Globodera rostochiensis suppresses CC-NB-LRR-mediated disease resistance in plants. Plant Physiol. 2012. 160:944–54. doi: 10.1104/pp.112.200188.

38. Eves-van den Akker S, Laetsch DR, Thorpe P, Lilley CJ, Danchin EG, Da Rocha M, et al. The genome of the yellow potato cyst nematode, Globodera rostochiensis, reveals insights into the basis of parasitism and virulence. Genome Biol. 2016. 17:124. doi: 10.1186/s13059-016-0985-1.

39. Abramovitch RB, Kim Y-J, Chen S, Dickman MB, Martin GB. Pseudomonas type III effector AvrPtoB induces plant disease susceptibility by inhibition of host programmed cell death. The EMBO journal. 2003. 22:60–9. doi: 10.1093/emboj/cdg006.

40. Ma L, Lukasik E, Gawehns F, Takken FLW. /the use of agroinfiltration for transient expression of plant resistance and fungal effector proteins in nicotiana benthamiana leaves. In: Bolton MD, Thomma BPHJ, editors. Plant Fungal Pathogens: Methods and Protocols. Totowa, NJ: Humana Press. 2012. p. 61–74.

41. van Ooijen G, Mayr G, Kasiem MM, Albrecht M, Cornelissen BJ, Takken FL. Structure-function analysis of the NB-ARC domain of plant disease resistance proteins. J Exp Bot. 2008. 59:1383–97. doi: 10.1093/jxb/ern045.

42. Bendahmane A, Kanyuka K, Baulcombe DC. The Rx gene from potato controls separate virus resistance and cell death responses. The Plant Cell. 1999. 11:781–91. doi: https://doi.org/10.1105/tpc.11.5.781

43. Tameling WIL, Baulcombe DC. Physical association of the NB-LRR resistance protein Rx with a Ran GTPase–Activating protein is required for extreme resistance to Potato virus X. The Plant Cell. 2007. 19:1682–94. doi: https://doi.org/10.1105/tpc.107.050880

44. Tameling WIL, Nooijen C, Ludwig N, Boter M, Slootweg E, Goverse A, et al. RanGAP2 mediates nucleocytoplasmic partitioning of the NB-LRR immune receptor Rx in the solanaceae, thereby dictating Rx function. The Plant Cell. 2010. 22:4176–94. doi: https://doi.org/10.1105/tpc.110.077461

45. Sacco MA, Mansoor S, Moffett P. A RanGAP protein physically interacts with the NB-LRR protein Rx, and is required for Rx-mediated viral resistance. The Plant Journal. 2007. 52:82–93. doi: https://doi.org/10.1111/j.1365-313X.2007.03213.x.

46. Du J, Rietman H, Vleeshouwers VGAA. Agroinfiltration and PVX agroinfection in potato and Nicotiana benthamiana. JoVE. 2014. (83):e50971. doi: doi:10.3791/50971.

47. Tameling WIL, Elzinga SDJ, Darmin PS, Vossen JH, Takken FLW, Haring MA, et al. The tomato R gene products I-2 and MI-1 are functional ATP binding proteins with ATPase activity. The Plant cell. 2002. 14:2929–39. doi: 10.1105/tpc.005793.

48. Petre B, Saunders DG, Sklenar J, Lorrain C, Win J, Duplessis S, et al. Candidate effector proteins of the rust pathogen Melampsora larici-populina target diverse plant cell compartments. Mol Plant Microbe Interact. 2015. 28:689–700. doi: 10.1094/MPMI-01-15-0003-R.

49. Petre B, Contreras M, Bozkurt T, Schattat M, Sklenar J, Schornack S, et al. Host-interactor screens of Phytophthora infestans RXLR proteins reveal vesicle trafficking as a major effector-targeted process. bioRxiv. 2020. doi: https://doi.org/10.1101/2020.09.24.308585.

50. Zess EK, Jensen C, Cruz-Mireles N, De la Concepcion JC, Sklenar J, Stephani M, et al. Nterminal beta-strand underpins biochemical specialization of an ATG8 isoform. PLoS Biol. 2019. 17:e3000373. doi: 10.1371/journal.pbio.3000373.

51. Isono E. TOL Keepers for ubiquitin-mediated trafficking routes in plant cells. Molecular Plant. 2020. 13:685–687. doi: 10.1016/j.molp.2020.04.005.

52. Mosesso N, Nagel M-K, Isono E. Ubiquitin recognition in endocytic trafficking–with or without ESCRT-0. Journal of cell science. 2019. 132:jcs232868. doi: 10.1242/jcs.232868

53. Moulinier-Anzola J, Schwihla M, De-Araújo L, Artner C, Jörg L, Konstantinova N, et al. TOLs function as ubiquitin receptors in the early steps of the ESCRT pathway in higher plants. Molecular Plant. 2020. 13:717–731. doi: 10.1016/j.molp.2020.02.012.

54. Wu C-H, Derevnina L, Kamoun S. Receptor networks underpin plant immunity. Science. 2018. 360:1300. doi: 10.1126/science.aat2623.

55. Witek K, Lin X, Karki HS, Jupe F, Witek AI, Steuernagel B, et al. A complex resistance locus in Solanum americanum recognizes a conserved Phytophthora effector. BioRxiv. 2020. doi: https://doi.org/10.1101/2020.05.15.095497.

56. Ding P, Sakai T, Shrestha RK, Perez NM, Guo W, Ngou BPM, et al. Chromatin accessibility landscapes activated by cell surface and intracellular immune receptors. bioRxiv. 2020. doi: 10.1101/2020.06.17.157040.

57. Ngou BPM, Ahn H-K, Ding P, Jones JDG. Mutual potentiation of plant immunity by cell-surface and intracellular receptors. bioRxiv. 2020. doi: 10.1101/2020.04.10.034173.

58. Pruitt RN, Zhang L, Saile SC, Karelina D, Fröhlich K, Wan W-L, et al. Arabidopsis cell surface LRR immune receptor signaling through the EDS1-PAD4-ADR1 node. bioRxiv. 2020. doi: 10.1101/2020.11.23.391516.

59. Yuan M, Jiang Z, Bi G, Nomura K, Liu M, He SY, et al. Pattern-recognition receptors are required for NLR-mediated plant immunity. bioRxiv. 2020. doi: 10.1101/2020.04.10.031294.

60. Juliana C, Fernandes-Alnemri T, Kang S, Farias A, Qin F, Alnemri ES. Non-transcriptional priming and deubiquitination regulate NLRP3 inflammasome activation. The Journal of biological chemistry. 2012. 287:36617–22. doi: 10.1074/jbc.M112.407130.

61. Yen H, Sugimoto N, Tobe T. Enteropathogenic Escherichia coli uses NleA to inhibit NLRP3 inflammasome activation. PLOS Pathogens. 2015. 11:e1005121. doi: 10.1371/journal.ppat.1005121.

62. Chen Y, Liu Z, Halterman DA. Molecular determinants of resistance activation and suppression by Phytophthora infestans effector IPI-O. PLOS Pathogens. 2012. 8:e1002595. doi: 10.1371/journal.ppat.1002595.

63. Karki HS, Abdullah S, Chen Y, Halterman DA. Natural genetic diversity in the potato resistance gene RB confers suppression avoidance from Phytophthora effector IPI-O4. bioRxiv. 2020. doi: 10.1101/2020.11.13.381905.

64. Coll RC, Hill JR, Day CJ, Zamoshnikova A, Boucher D, Massey NL, et al. MCC950 directly targets the NLRP3 ATP-hydrolysis motif for inflammasome inhibition. Nat Chem Biol. 2019. 15:556–9. doi: 10.1038/s41589-019-0277-7.

65. Tapia-Abellan A, Angosto-Bazarra D, Martinez-Banaclocha H, de Torre-Minguela C, Ceron-Carrasco JP, Perez-Sanchez H, et al. MCC950 closes the active conformation of NLRP3 to an inactive state. Nat Chem Biol. 2019. 15:560–4. doi: 10.1038/s41589-019-0278-6.

66. Winter V, Hauser M-T. Exploring the ESCRTing machinery in eukaryotes. Trends in Plant Science. 2006. 11:115–23. doi: https://doi.org/10.1016/j.tplants.2006.01.008.

67. De Craene J-O, Ripp R, Lecompte O, Thompson JD, Poch O, Friant S. Evolutionary analysis of the ENTH/ANTH/VHS protein superfamily reveals a coevolution between membrane trafficking and metabolism. BMC Genomics. 2012. 13:297. doi: 10.1186/1471-2164-13-297.

68. Conlan B, Stoll T, Gorman JJ, Saur I, Rathjen JP. Development of a rapid in planta BioID system as a probe for plasma membrane-associated immunity proteins. Front Plant Sci. 2018. 9:1882. doi: 10.3389/fpls.2018.01882.

69. Rühl S, Shkarina K, Demarco B, Heilig R, Santos JC, Broz P. ESCRT-dependent membrane repair negatively regulates pyroptosis downstream of GSDMD activation. Science. 2018. 362:956–60. doi: 10.1126/science.aar7607.

70. Vietri M, Radulovic M, Stenmark H. The many functions of ESCRTs. Nature reviews Molecular cell biology. 2020. 21:25–42. doi: https://doi.org/10.1038/s41580-019-0177-4.

71. Tsuchiya K. Inflammasome-associated cell death: Pyroptosis, apoptosis, and physiological implications. Microbiology and immunology. 2020. 64:252–69. doi: https://doi.org/10.1111/1348-0421.12771.

72. Bozkurt TO, Belhaj K, Dagdas YF, Chaparro-Garcia A, Wu C-H, Cano LM, et al. Rerouting of plant late endocytic trafficking toward a pathogen interface. Traffic. 2015. 16:204–26. doi: https://doi.org/10.1111/tra.12245.

73. Dagdas YF, Belhaj K, Maqbool A, Chaparro-Garcia A, Pandey P, Petre B, et al. An effector of the Irish potato famine pathogen antagonizes a host autophagy cargo receptor. eLife. 2016. 5. doi: 10.7554/eLife.10856.

74. Pandey P, Leary AY, Tümtas Y, Savage Z, Dagvadorj B, Tan E, et al. The Irish potato famine pathogen subverts host vesicle trafficking to channel starvation-induced autophagy to the pathogen interface. bioRxiv. 2020. doi: https://doi.org/10.1101/2020.03.20.000117.

75. Verzaux E, van Arkel G, Vleeshouwers VGAA, van der Vossen EAG, Niks RE, Jacobsen E, et al. High-resolution mapping of two broad-spectrum late blight resistance genes from two wild species of the Solanum circaeifolium group. Potato Research. 2012. 55:109–23. doi: 10.1007/s11540-012-9213-x.

76. Zhou J-M, Zhang Y. Plant immunity: danger perception and signaling. Cell. 2020. 181:978–89. doi: 10.1016/j.cell.2020.04.028.

77. Liu S, Lenoir CJG, Amaro TMMM, Rodriguez PA, Huitema E, Bos JIB. Virulence strategies of an insect herbivore and oomycete plant pathogen converge on host E3 SUMO ligase SIZ1 to suppress plant immunity. bioRxiv. 2020. doi: 10.1101/2020.06.18.159178.

78. Lu R, Malcuit I, Moffett P, Ruiz MT, Peart J, Wu A-J, et al. High throughput virus-induced gene silencing implicates heat shock protein 90 in plant disease resistance. The EMBO journal. 2003. 22:5690–9. doi: 10.1093/emboj/cdg546.

79. Rehman S, Postma WJ, Tytgat TO, Prins P, Qin L, Overmars H, et al. A secreted SPRY domain-containing protein (SPRYSEC) from the plant-parasitic nematode Globodera rostochiensis interacts with a CC-NB-LRR protein from a susceptible tomato. Mol Plant Microbe Interact. 2009. 22:330 – 40. doi: 10.1094/MPMI-22-3-0330.

80. Mei Y, Wright KM, Haegeman A, Bauters L, Diaz-Granados A, Goverse A, et al. The Globodera pallida SPRYSEC effector GpSPRY-414-2 that suppresses plant defenses targets a regulatory component of the dynamic microtubule network. Front Plant Sci. 2018. 9:1019. doi: 10.3389/fpls.2018.01019.

81. Cotton JA, Lilley CJ, Jones LM, Kikuchi T, Reid AJ, Thorpe P, et al. The genome and life-stage specific transcriptomes of Globodera pallida elucidate key aspects of plant parasitism by a cyst nematode. Genome Biology. 2014. 15:R43. doi: 10.1186/gb-2014-15-3-r43.

82. Berrow NS, Alderton D, Sainsbury S, Nettleship J, Assenberg R, Rahman N, et al. A versatile ligation-independent cloning method suitable for high-throughput expression screening applications. Nucleic Acids Research. 2007. 35:e45–e. doi: 10.1093/nar/gkm047.

83. Kourelis J, Kaschani F, Grosse-Holz FM, Homma F, Kaiser M, van der Hoorn RAL. A homology-guided, genome-based proteome for improved proteomics in the alloploid Nicotiana benthamiana. BMC Genomics. 2019. 20:722. doi: 10.1186/s12864-019-6058-6.

84. Segretin ME, Pais M, Franceschetti M, Chaparro-Garcia A, Bos JI, Banfield MJ, et al. Single amino acid mutations in the potato immune receptor R3a expand response to Phytophthora effectors. Mol Plant Microbe Interact. 2014. 27:624–37. doi: 10.1094/MPMI-02-14-0040-R.

85. Liu Y, Schiff M, Dinesh-Kumar SP. Virus-induced gene silencing in tomato. The Plant Journal. 2002. 31:777–86. doi: 10.1046/j.1365-313x.2002.01394.x.

86. Petre B, Saunders DG, Sklenar J, Lorrain C, Krasileva KV, Win J, et al. Heterologous expression screens in Nicotiana benthamiana identify a candidate effector of the wheat yellow rust pathogen that associates with processing bodies. PLoS One. 2016. 11:e0149035. doi: 10.1371/journal.pone.0149035.

87. Korbei B, Moulinier-Anzola J, De-Araujo L, Lucyshyn D, Retzer K, Khan Muhammad A, et al. Arabidopsis TOL proteins act as gatekeepers for vacuolar sorting of PIN2 plasma membrane protein. Current Biology. 2013. 23:2500–5. doi: https://doi.org/10.1016/j.cub.2013.10.036.

88. Madeira F, Park YM, Lee J, Buso N, Gur T, Madhusoodanan N, et al. The EMBL-EBI search and sequence analysis tools APIs in 2019. Nucleic acids research. 2019. 47:W636–W41. doi: 10.1093/nar/gkz268.

89. Stecher G, Tamura K, Kumar S. Molecular Evolutionary Genetics Analysis (MEGA) for macOS. Molecular Biology and Evolution. 2020. 37:1237–1239. doi: 10.1093/molbev/msz312.

90. Letunic I, Bork P. Interactive tree of life (iTOL) v3: an online tool for the display and annotation of phylogenetic and other trees. Nucleic acids research. 2016. 44:W242–W5. doi: 10.1093/nar/gkw290.

91. Yan P, Shen W, Gao X, Li X, Zhou P, Duan J. High-throughput construction of intron-containing hairpin RNA vectors for RNAi in plants. PLOS one. 2012. 7:e38186. doi: https://doi.org/10.1371/journal.pone.0038186.

92. Rain JC, Selig L, De Reuse H, Battaglia V, Reverdy C, Simon S, et al. The protein-protein interaction map of Helicobacter pylori. Nature. 2001. 409:211–5. doi: 10.1038/35051615.

93. Formstecher E, Aresta S, Collura V, Hamburger A, Meil A, Trehin A, et al. Protein interaction mapping: a Drosophila case study. Genome research. 2005. 15 3:376–84. doi: 10.1101/gr.2659105.

94. De la Concepcion JC, Franceschetti M, Maqbool A, Saitoh H, Terauchi R, Kamoun S, et al. Polymorphic residues in rice NLRs expand binding and response to effectors of the blast pathogen. Nat Plants. 2018. 4:576–85. doi: 10.1038/s41477-018-0194-x.

95. De la Concepcion JC, Franceschetti M, MacLean D, Terauchi R, Kamoun S, Banfield MJ. Protein engineering expands the effector recognition profile of a rice NLR immune receptor. eLife. 2019. 8. doi: 10.7554/eLife.47713.

96. Win J, Kamoun S, Jones AME. Purification of effector-target protein complexes via transient expression in Nicotiana benthamiana. Methods in molecular biology (Clifton, NJ). 2011. 712:181–94. doi: 10.1007/978-1-61737-998-7_15.

97. Lobstein J, Emrich CA, Jeans C, Faulkner M, Riggs P, Berkmen M. SHuffle, a novel Escherichia coli protein expression strain capable of correctly folding disulfide bonded proteins in its cytoplasm. Microbial Cell Factories. 2012. 11:753. doi: 10.1186/1475-2859-11-56.

98. Studier FW. Protein production by auto-induction in high-density shaking cultures. Protein Expression and Purification. 2005. 41:207–34. doi: https://doi.org/10.1016/j.pep.2005.01.016.

99. Bozkurt TO, Schornack S, Win J, Shindo T, Ilyas M, Oliva R, et al. Phytophthora infestans effector AVRblb2 prevents secretion of a plant immune protease at the haustorial interface. Proc Natl Acad Sci U S A. 2011. 108:20832–7. doi: 10.1073/pnas.1112708109.

100. Petre B, Saunders DG, Sklenar J, Lorrain C, Win J, Duplessis S, et al. Candidate effector proteins of the rust pathogen Melampsora larici-populina target diverse plant cell compartments. Molecular Plant-Microbe Interactions. 2015. 28:689–700. doi: 10.1094/MPMI-01-15-0003-R.

101. Chambers MC, Maclean B, Burke R, Amodei D, Ruderman DL, Neumann S, et al. A cross-platform toolkit for mass spectrometry and proteomics. Nature biotechnology. 2012. 30:918–20. doi: 10.1038/nbt.2377.

102. Perez-Riverol Y, Csordas A, Bai J, Bernal-Llinares M, Hewapathirana S, Kundu DJ, et al. The PRIDE database and related tools and resources in 2019: improving support for quantification data. Nucleic Acids Res. 2019. 47:D442–D50. doi: 10.1093/nar/gky1106.

